# Bioenergetic mapping of ‘healthy microbiomes’ via compound processing potential imprinted in gut and soil metagenomes

**DOI:** 10.1101/2023.11.05.565728

**Authors:** Craig Liddicoat, Robert A. Edwards, Michael Roach, Jake M. Robinson, Kiri Joy Wallace, Andrew D. Barnes, Joel Brame, Anna Heintz-Buschart, Timothy R. Cavagnaro, Elizabeth A. Dinsdale, Michael P. Doane, Nico Eisenhauer, Grace Mitchell, Bibishan Rai, Sunita Ramesh, Martin F. Breed

**Affiliations:** College of Science and Engineering, Flinders University, Bedford Park, South Australia, Australia; Environmental Research Institute, University of Waikato, Hamilton, Aotearoa New Zealand; Swammerdam Institute for Life Sciences, University of Amsterdam, 1098 XH Amsterdam, The Netherlands; German Centre for Integrative Biodiversity Research (iDiv), 04103 Leipzig, Germany; Institute of Biology, Leipzig University, 04103 Leipzig, Germany; Manaaki Whenua – Landcare Research, Hamilton, Aotearoa New Zealand

**Keywords:** Metagenomics, compound processing potential, healthy microbiome, human health, bioenergetic mapping, van Krevelen diagram

## Abstract

Despite mounting evidence of their importance in human health and ecosystem functioning, the definition and measurement of ‘healthy microbiomes’ remain unclear. More advanced knowledge exists on health associations for compounds used or produced by microbes. Environmental microbiome exposures (especially via soils) also help shape, and may supplement, the functional capacity of human microbiomes. Given the synchronous interaction between microbes, their feedstocks, and micro-environments, with functional genes facilitating chemical transformations, there exists an intriguing opportunity to examine microbiomes in terms of their capacity to process compounds relevant to human health. Here we integrate functional genomics and biochemistry frameworks to derive new quantitative measures of the potential for human gut and environmental soil metagenomes to process major compound classes (e.g., lipids, carbohydrates) and selected biomolecules (e.g., vitamins, short-chain fatty acids) linked to human health. Metagenome functional potential profile data were translated into a universal compound mapping ‘landscape’ based on bioenergetic van Krevelen mapping of function-level meta-compounds and corresponding functional relative abundances, reflecting imprinted genetic capacity of microbiomes to metabolize an array of different compounds. We show that measures of ‘compound processing potential’ associated with human health and disease (examining atherosclerotic cardiovascular disease, colorectal cancer, type 2 diabetes and anxious-depressive behavior case studies), and displayed seemingly predictable shifts along gradients of ecological disturbance in plant-soil ecosystems (three case studies). Ecosystem quality explained 60–92% of variation in soil metagenome compound processing potential measures in a post-mining restoration case study dataset. With growing knowledge of the varying proficiency of environmental microbiota to process human health associated compounds, we might design environmental interventions or nature prescriptions to modulate our exposures, thereby advancing microbiota-oriented approaches to human health. Compound processing potential offers a simplified, integrative approach for applying metagenomics in ongoing efforts to understand and quantify the role and linkages of microbiota in environmental- and human-health.

## 1. Introduction

Microbial communities (microbiota), their feedstocks (substrates, nutrients) and environmental conditions (e.g., pH, redox potential, temperature, moisture, salinity) work in concert to drive microbially-mediated reactions essential to fueling life on Earth (Averill et al., 2022). Microbiomes (microbiota, genetic material and metabolites) are critical to the health and functioning of humans (Banerjee and van der Heijden, 2022; Gilbert et al., 2018; Round and Mazmanian, 2009) and ecosystems (Delgado-Baquerizo et al., 2020; Sokol et al., 2022). However, clear definitions of what constitutes a ‘healthy microbiome’ remain elusive (Eisenstein, 2020). More advanced knowledge exists on health associations for the compounds used or produced by microbes. Because microbes, their feedstocks and micro-environments interact synchronously—using functional genes to facilitate chemical transformations—this presents an intriguing opportunity to examine microbiomes through their potential to process compounds associated with human health.

Human microbiomes are supplemented and partly shaped by environmental microbial exposures (Banerjee and van der Heijden, 2022; Gilbert et al., 2018). Therefore, the involvement of environmental microbiomes in processing human health-associated compounds is also of interest. Transfer of environmental microbiota to humans may help supplement important functional capacity, protective microbiota, and immune-signaling agents, particularly in infants, but also in adults who have depleted microbiota due to antibiotic use, poor diet, lifestyle or other health incidents (Banerjee and van der Heijden, 2022; Brame et al., 2021; Flandroy et al., 2018; Roslund et al., 2022). We know that microbiota are shaped by the resources they utilize and the environments they inhabit (Fierer, 2017; Gilbert et al., 2018). If the functional composition of microbiota varies predictably along environmental gradients, then through design, management, and behavior we should be able to modulate our exposure to health-promoting versus disease-associated microbes. Soils, in particular, can represent a rich source of microbial diversity with potential to support human health (Banerjee and van der Heijden, 2022; Li et al.; Liddicoat et al., 2018; Liddicoat et al., 2020; Ottman et al., 2018; Sun et al., 2023). Microbiota in plant-soil systems are shaped by macro-scale factors including climate, soil characteristics, vegetation composition, diversity, land use and management (Delgado-Baquerizo et al., 2018; Fierer, 2017). With the prospect of cost-effectively encouraging health-promoting microbes, it is frequently asked, “What type of environment is best?” Yet, the attributes of health-promoting environmental microbiomes, including potential functional overlaps with human microbiomes, remain understudied.

Microbes typically operate as a community (Barton and Northup, 2011) where many taxa may be highly efficient at performing certain functions, but they lack functional capacity for stand-alone survival (Zengler and Zaramela, 2018). Complex cross-feeding and resource sharing in the extracellular space (Daisley et al., 2021), and involvement of multiple taxa in functional pathways, suggest that community-scale functional profiles (rather than specific microbial taxa) underpin the health-supporting capacity of microbiota. Indeed, it is this community-scale complexity that has hindered progress towards clear definitions of a healthy microbiome. Nevertheless, researchers want to better understand the assembly and structure of health-promoting microbiomes to improve the course of microbiome-associated diseases. Disease-associated microbiota are often characterized by a loss of diversity and overgrowth by opportunistic pathogens (Belizario and Napolitano, 2015; Round and Mazmanian, 2009), but it may be unclear whether they are the facilitators or followers of disease.

Many microbiome-associated diseases are linked to bioenergetic mechanisms (Daisley et al., 2021), with oxidation-reduction (redox) potential recognized as a key factor shaping microbial communities. The healthy anaerobic gut favors obligate anaerobes, whereas dysbiosis is often accompanied by increased oxygenation of the colonic epithelium and expanded abundance of oxygen-tolerant facultative anaerobic bacteria (Litvak et al., 2018; Rivera-Chávez et al., 2017). Oxygen is a highly electronegative element important in shaping electrochemical gradients, biochemical reactions, and gene expression (Wilson et al., 2020). Oxygen content varies in different types of organic matter (i.e., microbial feedstocks). Yet, the interrelationship between bioenergetic (electrochemical) status of compounds, microbial environments, microbiota development and human health appears to receive insufficient attention. In soils, redox potential varies with weather, vegetation, land use, management, drainage, organic-content, vicinity to roots, soil characteristics, and microbial activity (Hinsinger et al., 2009; Husson, 2013). At the molecular level, redox potential shapes what kind of molecules can be made and how energy is stored. Therefore, a compound-oriented examination of healthy microbiomes might capitalize on available knowledge linking compounds with human health, while also considering deterministic influences of bioenergetic (electrochemical) context.

Here, we examine the functional potential of gut and soil microbiota from a compound processing potential (CPP) viewpoint, to assess patterns in human health and disease, and with gradients of ecosystem maturity (or quality). Such an approach might discern health-versus disease-promoting microbiotas from the types of compounds they are attuned to consuming or producing (reflecting microbial feedstocks and metabolites respectively). Previous metabolome prediction frameworks (e.g., Diener et al., 2020; Garza et al., 2018; Mallick et al., 2019; Neveu et al., 2023) rely variously on supplementary metabolome training datasets, microorganism-specific genome-scale metabolic models, taxonomic abundance estimates, and assumed environments (e.g., human gut), to model short-term reaction rates. Here, we wanted to exploit community-scale compound-oriented information that might be embedded within metagenomes, regardless of taxa present. As functional potential profiling from whole genome sequencing (or shotgun metagenomics) does not directly measure functions performed, we characterized microbiota-linked CPP from DNA sequencing, without direct measurement of compounds. Our conceptual interest was to characterize the distribution of available community-scale genetic capacity of microbiomes to process an array of compound classes and specific biomolecules (compounds produced by living organisms), which may be present over a period of time and with variation in feedstocks and environmental conditions, consistent with dynamic influences that have shaped existing microbiota structure. We premised that the ease of transformation between health-associated compounds and other compounds that closely resemble them will depend on stoichiometric and energetic similarities, microbiota functional diversity, and environmental conditions.

We utilized a framework that integrates information about compounds, bioenergetics, and environmental conditions. The complexity-reducing van Krevelen (vK) coordinate space offers a simplified and intuitive bioenergetic framework for approximate mapping of compounds based on their carbon (C), oxygen (O) and hydrogen (H) content, while also reflecting energy density and principal axes that explain microbial distribution and chemical speciation (Fig. 1 and Fig. S1) (D’Andrilli et al., 2015; Wu et al., 2018). Compounds are mapped into vK space using their O:C and H:C molar ratios (x- and y-axis respectively). We surmised this framework could offer an exhaustive and intuitive mapping space to summarize the nature of microbiota-mediated functional reactions in a way that reflects the collective mean or dominant compound properties, reaction stoichiometries, and potential overlaps between dietary or environmental substrates, and key health-associated compounds.

**Fig. 1.**
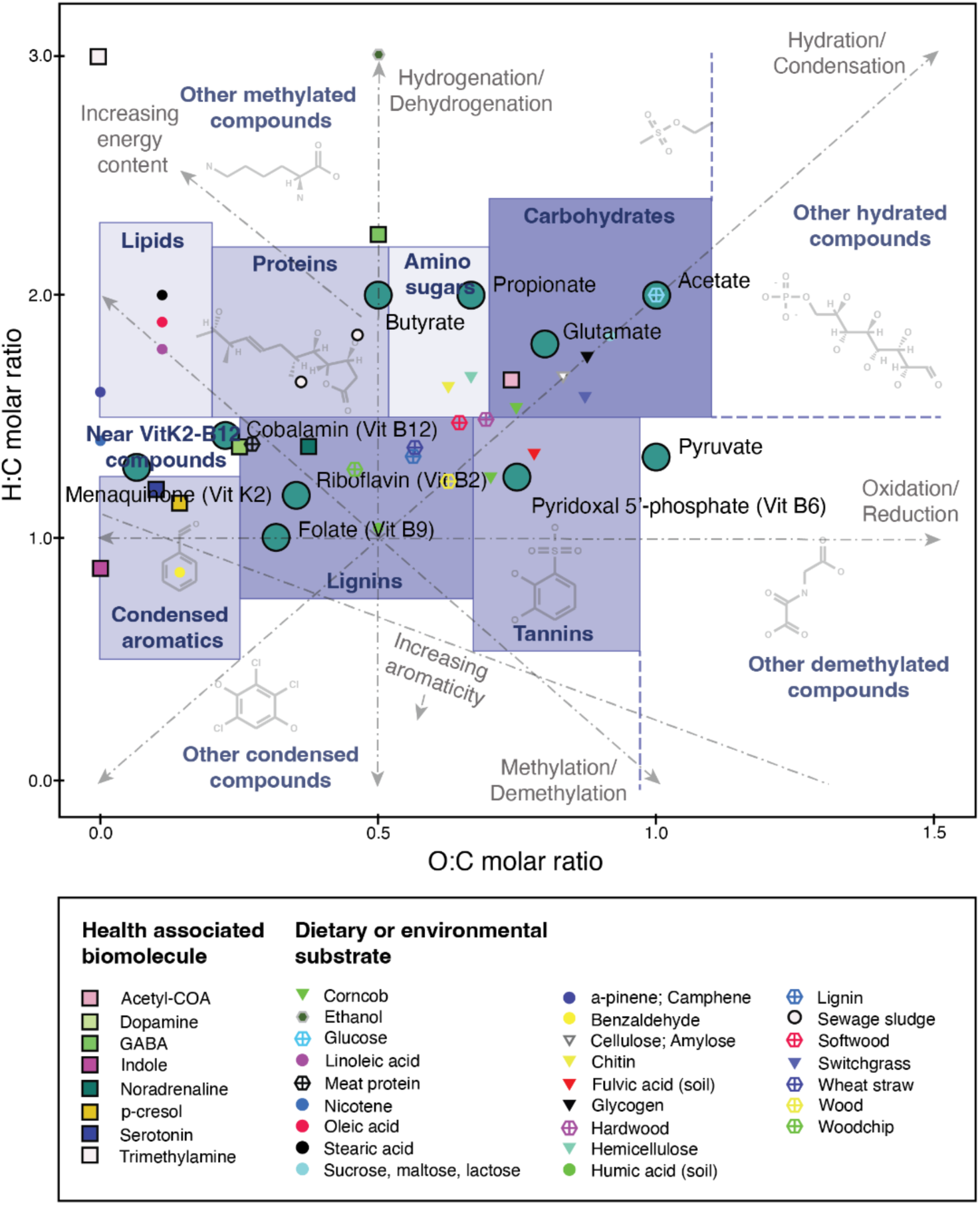
Van Krevelen (vK) coordinate space (adapted from D’Andrilli et al., 2015), displaying major compound classes (purple zones and text), key gradients (grey axes and text), focus biomolecules examined in this study (large dots), and additional example health-associated biomolecules, dietary or environmental substrates (legend) (see Tables S1, S2). vK zones were adapted from (Wu et al., 2018). Key gradients include oxidation-reduction (x-axis), hydrogenation-dehydrogenation (y-axis), hydration-condensation (top-right to bottom-left), methylation-demethylation (top-left to bottom-right) and increasing energy content (towards top-left).

Specifically, we combined SUPER-FOCUS functional profiling annotations (Silva et al., 2015), the comprehensive ModelSEED (Seaver et al., 2020) functional-biochemistry database system and vK coordinate mapping to assign functional potential relative abundances from human gut and soil sample metagenomes to overall mean function-level vK coordinates. This approach effectively mapped every SUPER-FOCUS function (where feasible via available corresponding database information), based on one or more associated chemical reactions, to a function-level ‘meta-compound’ represented in the two-dimensional vK space (Fig. S2, Methods). We aimed to: (1) investigate measures of microbiota CPP imprinted in human gut and soil metagenomes; (2) test for differences in human health and disease, and in disturbed, restored and natural ecosystems; and (3) identify compounds linked to human health with differing potential metabolism (i.e., CPP) by environmental microbiota due to differences in ecosystem quality. We hypothesized this universal bioenergetic compound mapping approach might identify CPP profiles, and possible overlaps in human and environmental metagenomes, that could inform the definition and future shaping of ‘healthy microbiomes’. We were also keen to explore whether CPP measures might enhance the interpretability and accessibility of metagenomics data to aid hypothesis building and prioritizing future research.

## 2. Materials and Methods

### 2.1. Case study datasets

We assessed metagenomics samples from four human health and disease datasets, comprising atherosclerotic cardiovascular disease (ACVD) (Jie et al., 2017), colorectal cancer (Zeller et al., 2014), type 2 diabetes (T2D) (Forslund et al., 2015) and problem (anxious-depressive) behaviors in children (Flannery et al., 2020); and three environmental soil datasets from ecological restoration and disturbed versus natural plant-soil systems (Barnes et al., 2020; Bissett et al., 2016; Sun and Badgley, 2019) (Table 1; further detail in Appendix A Supporting Information).

**Table 1.** Description of case study metagenome datasets.

| Case study focus | Main comparison variable or diagnosis groups (and sample/subject numbers<sup>a</sup>) | Metagenome functional profile characteristics (sample mean ± s.d.)<sup>b</sup> | Source data reference; Country of origin |
| --- | --- | --- | --- |
| **Human gut** |  |  |  |
| Atherosclerotic cardiovascular disease (ACVD) | ACVD ($n_F = 53$, $n_M = 157$) or normal healthy ($n_F = 101$, $n_M = 69$). Total $n = 380$. | Functions<sup>0</sup> $n = 21,117$ ($9301 \pm 3523$)<br>Functions<sup>vK</sup> $n = 10,829$ ($5051 \pm 1843$)<br>Total fxn rel abund<sup>vK</sup> = $63.2 \pm 3.5$ %<br>vK coordinates $n = 2535$ ($1509 \pm 502$) | SRA accession PRJEB21528 (Jie et al., 2017); China |
| Colorectal cancer | Colorectal cancer ($n_F = 24$, $n_M = 29$) or normal healthy ($n_F = 33$, $n_M = 27$). Total $n = 113$. | Functions<sup>0</sup> $n = 12,896$ ($2809 \pm 1924$)<br>Functions<sup>vK</sup> $n = 6932$ ($1750 \pm 1117$)<br>Total fxn rel abund<sup>vK</sup> = $66.7 \pm 2.7$ %<br>vK coordinates $n = 1954$ ($711 \pm 362$) | SRA accession PRJEB6070 (Zeller et al., 2014); France |
| Type 2 diabetes (T2D) and impaired glucose tolerance (IGT) | T2D with no Metformin treatment (T2D Met-, $n = 33$), T2D with Metformin (T2D Met+, $n = 20$), IGT ($n = 49$), or normal healthy ($n = 43$). Subjects are females only. Total $n = 145$. | Functions<sup>0</sup> $n = 19,099$ ($9916 \pm 2393$)<br>Functions<sup>vK</sup> $n = 9916$ ($4490 \pm 874$)<br>Total fxn rel abund<sup>vK</sup> = $64.4 \pm 2.2$ %<br>vK coordinates $n = 2393$ ($1431 \pm 202$) | SRA accession PRJEB1786 (Forslund et al., 2015); Sweden |
| Problem behaviors in children | First principal component (PC1) of anxious-depressive problem behaviors, examined as either numeric scores ($n_F =$ | Functions<sup>0</sup> $n = 20,599$ ($10282 \pm 2481$)<br>Functions<sup>vK</sup> $n = 10,322$ ($5507 \pm 1224$)<br>Total fxn rel abund<sup>vK</sup> = $54.9 \pm 1.3$ % | SRA accession PRJNA496479 (Flannery et al., |
| | 20, $n_M = 17$); or high/low PC1 groups<sup>c</sup> (high PC1 $n_F = 10$, $n_M = 8$; Low PC1 $n_F = 10$, $n_M = 9$). Total $n = 37$. | vK coordinates $n = 2459$ ($1645 \pm 281$) | 2020); United States |
| **Soils** |  |  |  |
| People Cities and Nature (PCaN) urban forest ecosystem restoration | Soil samples spanned young to old revegetation age, and remnant sites (treated as ordinal variables). Data were separated into pH-based groups: strongly acidic (pH < 4.5, 10-40 yr old, remnant, $n = 8$); and acidic-neutral soils ($4.5 < \text{pH} < 7$, 11-48 yr old, $n = 10$). Total $n = 18$. | Functions<sup>0</sup> $n = 36,324$ ($27,969 \pm 1011$)<br>Functions<sup>vK</sup> $n = 18,197$ ($14,690 \pm 396$)<br>Total fxn rel abund<sup>vK</sup> = $61.7 \pm 0.3$ %<br>vK coordinates $n = 3302$ ($2965 \pm 49$) | Aotearoa Genomic Data Repository project AGDR00045 (Barnes et al., 2020); Aotearoa New Zealand |
| Post-mining forest ecosystem restoration | Soil samples spanned revegetation ages of 6, 12, 22, 31 years, and unmined (UM) samples (treated as ordinal variables). Comprising five age-based groups, each with three replicates. Total $n = 15$. | Functions<sup>0</sup> $n = 30,125$ ($20,328 \pm 767$)<br>Functions<sup>vK</sup> $n = 15,576$ ($11,051 \pm 334$)<br>Total fxn rel abund<sup>vK</sup> = $61.9 \pm 0.3$ %<br>vK coordinates $n = 3076$ ($2537 \pm 54$) | MG-RAST project mgp16379 (Sun and Badgley, 2019); United States |
| Australian Microbiome Initiative (AMI) disturbed versus natural | Disturbed ($n = 29$) or natural ($n = 55$) soils, comprising temperate climate zone, surface (0-10cm) with 7.5–45% clay content (i.e., avoiding very sandy and very clayey soils). Total $n = 84$. | Functions<sup>0</sup> $n = 37,335$ ($22,959 \pm 1172$)<br>Functions<sup>vK</sup> $n = 18,551$ ($12,212 \pm 553$)<br>Total fxn rel abund<sup>vK</sup> = $60.8 \pm 1.0$ %<br>vK coordinates $n = 3326$ ($2700 \pm 69$) | AMI Data Portal [Data accessed Sep 2022] (Bissett et al., 2016)<sup>d</sup>; Australia |

Notes: <sup>a</sup>F = females, M = males. <sup>b</sup>Number of functions<sup>0</sup> from initial SUPER-FOCUS profiling, versus functions<sup>vK</sup> (fxn) with available compound information for mapping to vK coordinates. <sup>c</sup>PC1 of problem behaviors were analyzed as numeric values for visualizing and assessing CPP<sub>class</sub> and CPP<sub>density</sub> data; and high vs. low PC1 groups for CPP<sub>ASALR</sub> data. High PC1 values represent more problematic behavior. <sup>d</sup>AMI Data portal URL: https://data.bioplatforms.com/organization/australian-microbiome.

### 2.2. Compound processing potential (CPP) mapping approach

CPP measures were derived via the following steps (Fig. 2; further detail in Appendix A Supporting Information):

a. Shotgun metagenomics raw sequences were accessed, and bioinformatic steps were run on Flinders University DeepThought high-performance computing facility (Flinders_University, 2021).
b. Raw sequence data were inspected using FastQC (v0.11.9; Andrews, 2018) and quality control trimming performed using Fastp (v0.23.2; Chen et al., 2018).
c. Functional potential profiles were derived from good quality read 1 sequences using SUPER-FOCUS (Silva et al., 2015) functional annotation software, linked to the Diamond sequence aligner (v0.9.19; Buchfink et al., 2021) and version 2 100% identity-clustered reference database (100_v2; https://github.com/metageni/SUPER-FOCUS/issues/66). Where subjects/samples were represented by multiple sequence files, the combined SUPER-FOCUS outputs were normalized so that the total functional relative abundances summed to 100% in each subject/sample.
d. Every SUPER-FOCUS function (output row) was translated to one or more corresponding chemical reaction(s) using a purpose-built R-script algorithm based on ModelSEED database lookup tables (from https://github.com/ModelSEED/ModelSEEDDatabase; accessed 10 Aug 2022). The algorithm sought matches based on either: full matching of functional hierarchies (using subsystem-class, -subclass, - name and -role); detection of EC number; or matching of SUPER-FOCUS function name within ModelSEED lookup tables for reactions (reaction name or alias), subsystems (role), or reaction-pathways (external reaction name).
e. Every chemical reaction was converted to mean vK coordinates (O:C and H:C molar ratios), considering all C-containing reaction input and product compounds and weighted according to reaction stoichiometry (see example in Fig. 2.). Compounds not containing C were ignored due to undefined O:C and H:C ratios. Data for compounds were based on Hill system chemical formulae in protonated form.
f. Overall mean vK coordinates were calculated for each SUPER-FOCUS function via averaging one or more linked chemical reactions. The majority of SUPER-FOCUS functional relative abundances were successfully mapped to vK coordinates (sample ranges 52-84%, means 55-67%, Table 1, Table S3).
g. In vK coordinate space, we analyzed the spatial assignment of functional relative abundances to derive the following CPP data types:

1. CPP_class_: summed functional relative abundances mapping to major compound classes (based on pre-defined zones from Wu et al., 2018) (see Fig. 1, Table S1).
2. CPP_ASALR_: to address large variation in CPP_class_ values across case studies, we implemented a first-pass normalization aiming to account for microbial activity levels, here termed amino sugar adjusted log ratio (ASALR) data. Amino sugars have previously been used as a biomarker of microbial residue turnover (Hu et al., 2018) as they are major components of bacterial and fungal cell walls (peptidoglycan and chitin). CPP_class_ values were divided by the CPP_class_ value for amino sugars, followed by a variance-stabilizing log10-transformation.
3. CPP_density_: captured the density of functional relative abundances within radial buffers of varying proximity (radii of 0.05, 0.1, 0.15, 0.2, 0.25 vK units) to selected focus biomolecules (Fig. 1, Table S2, Appendix A Supporting Information). These comprised short-chain fatty acids: acetate, propionate, and butyrate; vitamins: riboflavin (B2), cobalamin (B12), pyridoxal 5’-phosphate (B6), folate (B9), menaquinone (K2); and glutamate and pyruvate. Functional relative abundances within radial buffers were summed then divided by the respective area in vK units^2^.
4. Compound-associated vK coordinates: Each SUPER-FOCUS functional row was translated to corresponding vK coordinates (as used for the above spatial assignments). Further supplementary analyses were undertaken at the vK coordinate level, including network analyses (in the T2D case study), differential abundance analysis (in the problem behavior case study), and weighted-mean vK coordinate analysis within major compound zones.

**Fig. 2.**
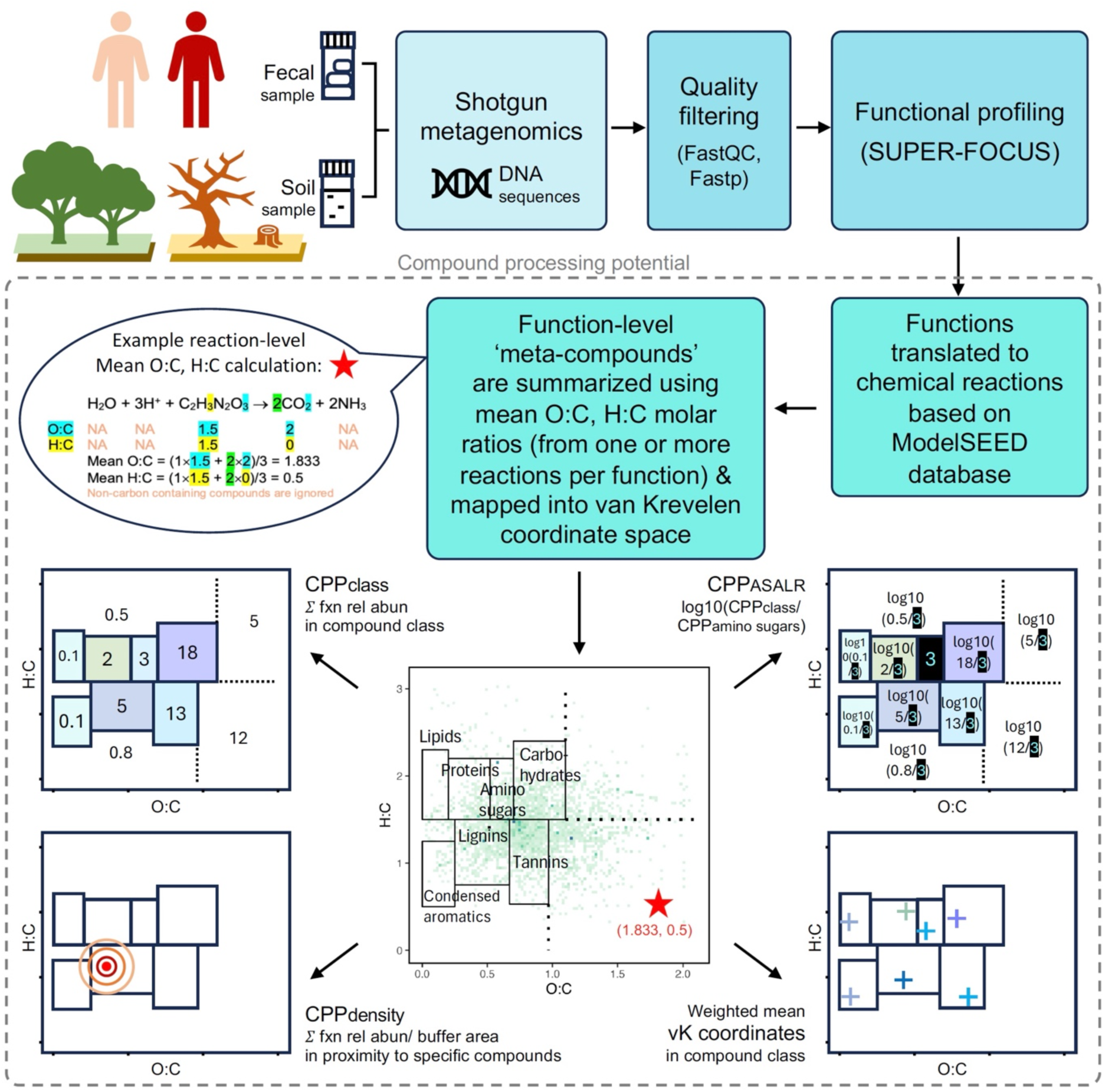
Flow chart of compound processing potential (CPP) analyses. The example reaction shows calculation of mean O:C and H:C molar ratios (e.g., 1.833, 0.5) based on stoichiometry of all carbon-containing compounds. Mean O:C and H:C values are then averaged across one or more chemical reactions per function, to derive function-level vK coordinates. Each function-level mapping into vK space represents a conceptual ‘meta-compound’ based on the collective elements and stoichiometry of all C-containing compounds involved in all reactions corresponding to a single microbiota-mediated function.

### 2.3. Data visualization and statistical analyses

Details of visualization and statistical testing using standard approaches are provided in Appendix A Supporting Information.

## 3. Results

### 3.1. Human health and disease

Gut metagenome CPP_class_ (Figs. S4, S8, S12, S18, Table S3) and CPP_density_ (Figs. S5-S6, S9-S10, S13-S14, S19-S20, Table S4) data produced strong associations in ACVD and colorectal cancer compared to normal subjects (detailed below). Many patterns observed in CPP_class_ data were reinforced in the putative activity-normalized CPP_ASALR_ measures (Fig. 3, Table S5), and this transformed data format showed stabilized variance across case study datasets. Interestingly, across all CPP_class_, CPP_density_, and CPP_ASALR_ measurements, when associations were found in both sexes they were always in the same direction (Tables S3-S5). In the T2D (female only) and problem behavior case studies, we observed far fewer relationships in the coarse CPP_class_, CPP_ASALR_ or biomolecule-focused CPP_density_ data. Therefore, we pursued network analyses and differential abundance analyses respectively in these case studies, as illustrative examples of more detailed supplementary analyses. Weighted mean vK-coordinate analyses produced striking associations in ACVD (Fig. S7 Tables S6-S7), but weaker effects in other case studies (Figs. S11, S15, S21, Tables S8-S12). The statistical test results described below are detailed in Appendix A.

**Fig. 3.**
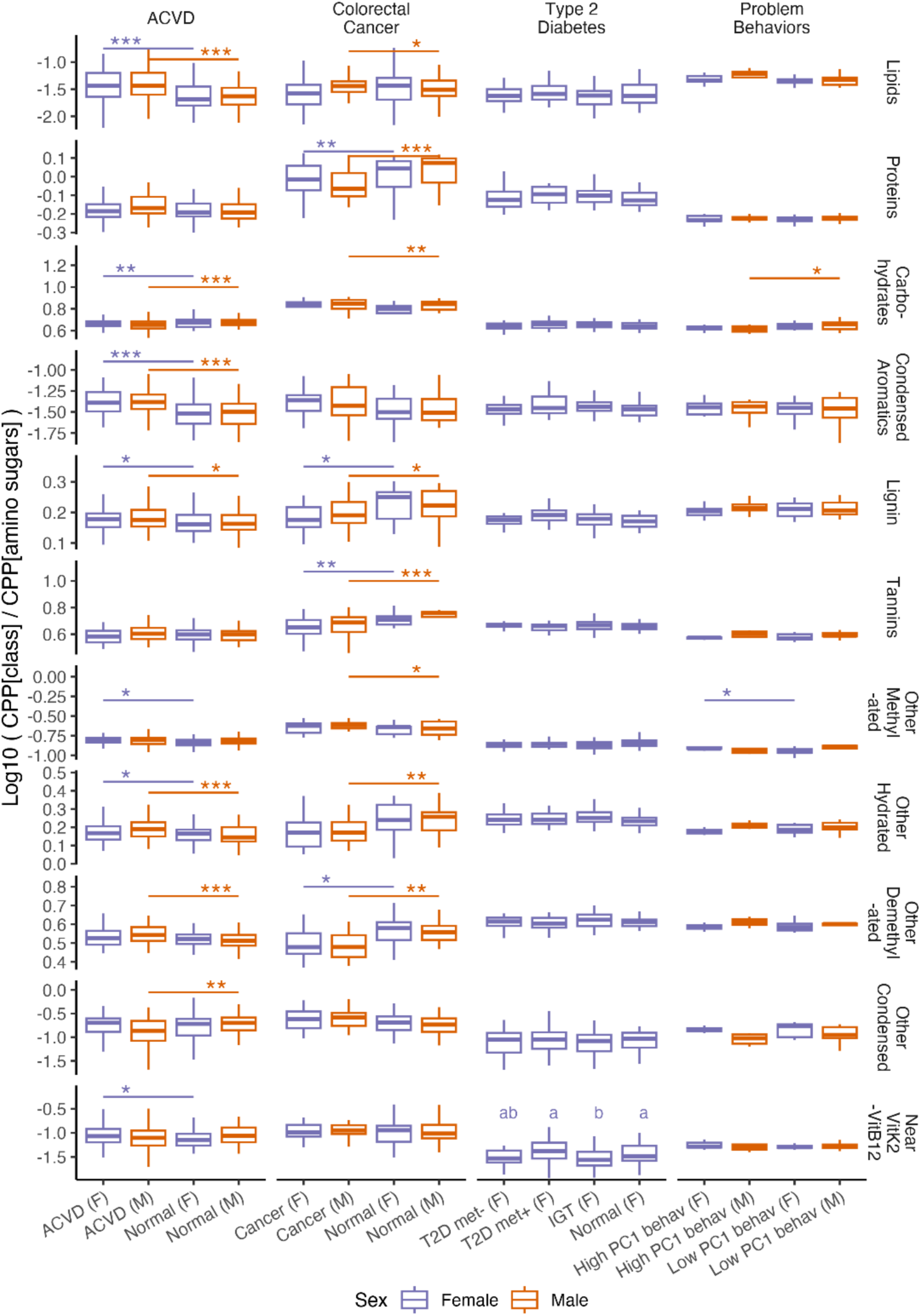
Amino sugar-adjusted log ratio compound processing potential (CPP_ASALR_), representing putative microbiota activity-normalized values, in normal healthy and diseased female (F) and male (M) subjects for atherosclerotic cardiovascular disease (ACVD), colorectal cancer, type 2 diabetes (T2D) with and without Metformin treatment (met +/-), impaired glucose tolerance (IGT), and high and low first principal component (PC1) problem behavior values. For visualization purposes outlying values are not shown. However, statistical tests were based on all data (Table S5). Sample sizes are detailed in Table 1. Tests for differences are performed within a single sex. In T2D data, groups not sharing a letter are different.

ACVD associated with CPP_class_ values, compared to normal subjects, in the form of increased potential metabolism of lipids (in females – F, and in males – M), proteins (M), condensed aromatics (F, M) lignin (F, M), tannins (M), other hydrated compounds (F, M), other demethylated compounds (M), and near vitamin K2-vitamin B12 compounds (F); contrasting with decreased potential metabolism of carbohydrates (F, M) and other condensed compounds (M). Despite all samples initially summing to 100% total functional relative abundances, there was less complete conversion to identifiable reactions and total sum CPP_class_ data in females only for ACVD compared to normal cases. Many of these patterns were reinforced in the CPP_ASALR_ data, with ACVD notably associated in both sexes with increased potential metabolism of lipids, condensed aromatics, lignin, and other hydrated compounds; but decreased potential metabolism of carbohydrates. For CPP_density_ measures within 0.05 vK unit radii, ACVD associated with increased potential metabolism of propionate (F), vitamin B12 (F), vitamin B6 (M), vitamin B9 (F, M), vitamin K2 (F, M) and pyruvate (M); but decreased potential metabolism of acetate (M) and glutamate (F, M).

Colorectal cancer also associated prominently with CPP_class_ values, specifically increased potential metabolism of lipids (M), amino sugars (F, M), condensed aromatics (M), lignin (M), other condensed compounds (F, M) and near vitamin K2-vitamin B12 compounds (M); but decreased potential metabolism of proteins (F, M) and tannins (F, M). Total sum CPP_class_ data were reduced in male colorectal cancer cases compared to normal. CPP_ASALR_ values in colorectal cancer subjects of both sexes were associated with decreased potential metabolism of proteins, lignin, tannins, and other demethylated compounds. For CPP_density_ measures within 0.05 vK unit radii, colorectal cancer associated with increased potential metabolism of vitamin B2 (M), vitamin B12 (F, M), vitamin B9 (F, M), and pyruvate (M). Weighted mean vK-coordinates varied between colorectal cancer and normal subjects with different centroids in condensed aromatics (F), lignin (M), and tannins (M); and different beta-dispersion or spread in proteins (F, M) and condensed aromatics (F, M).

In the female-only T2D case study results, we found CPP_class_ values decreased for other methylated compounds in IGT compared to normal subjects and decreased for near vitamin K2-vitamin B12 compounds in IGT compared to normal, with a reduced total sum CPP_class_ in normal compared to other subjects. No associations were found with CPP_density_ values. CPP_ASALR_ values were decreased for near vitamin K2-vitamin B12 compounds in IGT compared to normal and T2D Met+ subjects. From analysis of weighted mean vK-coordinates among diagnosis groups, a difference in centroids was found only in the compound class of proteins. Lacking interpretable results at the compound class or focus biomolecule levels, further investigation was performed via network analysis (Appendix A Supporting Information) on 20 subjects in each diagnosis group based on inferred correlations between functional relative abundances mapped to unique vK-coordinates. Network analyses were based on commonly observed vK coordinates (present in at least 60% of samples and minimum 2% sum functional relative abundance across samples). Network diagrams (Fig. 4) and structure dendograms (Fig. S16) display a transition in their complexity, number of nodes and fraction of negative edges (interactions) from simplest in normal and IGT subjects to most complex in T2D Met+ and T2D Met-subjects. Comparing network characteristics for the four groups (normal, IGT, T2D Met-, T2D Met+; n = groups of 20) to a bootstrapped (B = 1000) density distribution of randomly resampled networks (n = 20, drawn from the same pool of 80 subjects) (Fig. S17, Table S17) we found normal healthy subjects had the lowest fraction of negative edges and the highest degree centralization. Untreated disease, T2D Met-, had the lowest closeness centralization (graph-level inverse of average geodesic distance between nodes); and borderline significant results for the highest fraction of negative edges (negative correlations between vK-coordinates), lowest betweenness centralization (graph-level centrality based on broker positions connecting others), and lowest mean distance (average path length between nodes). In short, normal subjects appear to have far less correlations between vK-coordinates (fewer nodes / vertices), and for the nodes and links that are present they are largely positively correlated and highly interlinked. Whereas T2D Met-(untreated disease) is characterized by a much larger number of negatively correlated vK-coordinate nodes, which on average have shorter links, and are less well connected-up across the whole network.

**Fig. 4.**
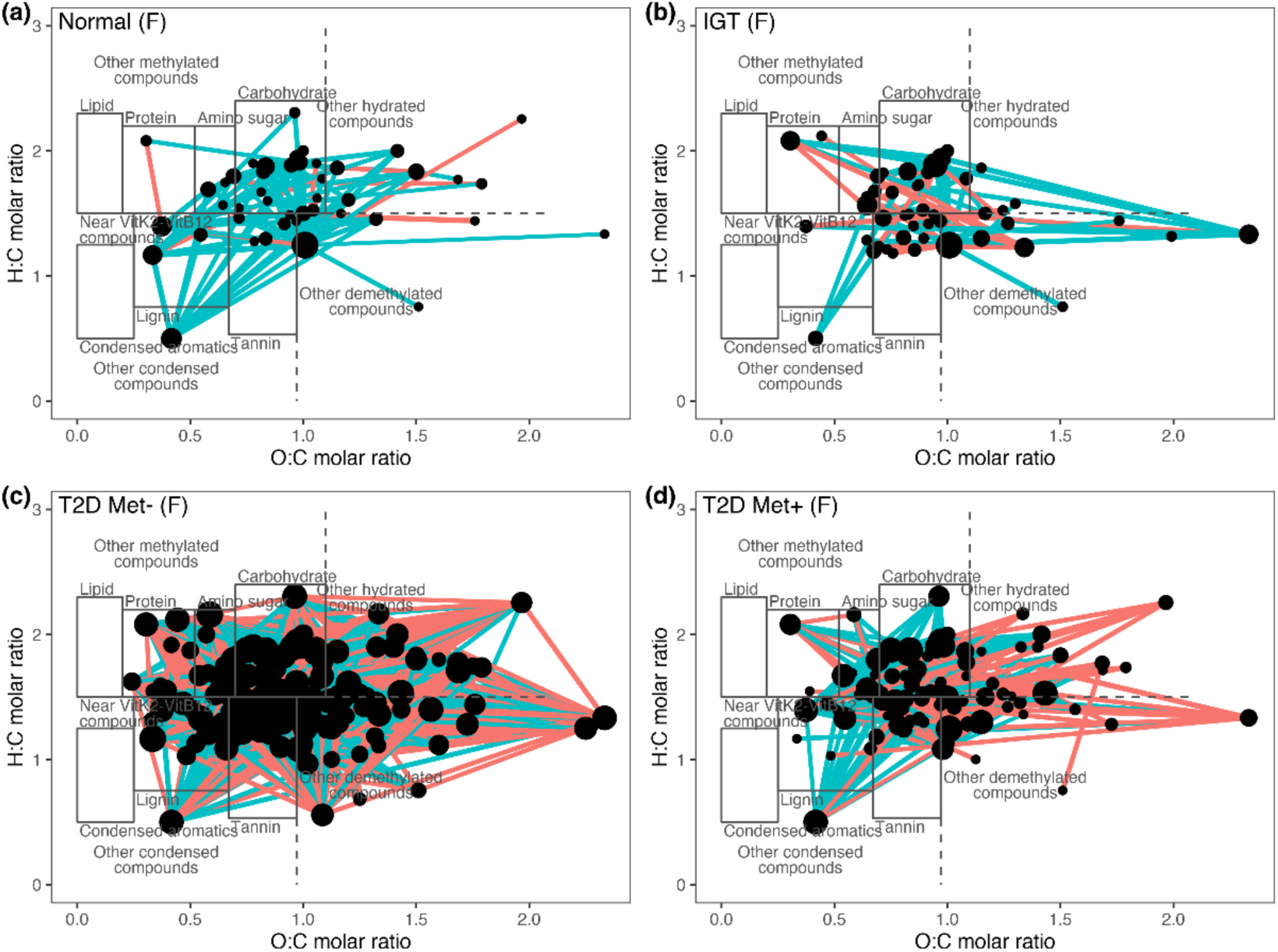
Network diagrams based on commonly observed vK coordinates for female subjects (n = groups of 20) with diagnoses: (a) normal, (b) impaired glucose tolerance (IGT), (c) type 2 diabetes without Metformin (T2D Met-), and (d) type 2 diabetes with Metformin (T2D Met+). Nodes are located according to unique compound-associated vK coordinates, with size reflecting node degree (number of linked significant correlations). Links between nodes display positive (aqua color) and negative (red color) correlations (p ≤ 0.05).

Problem behaviors displayed few significant associations in CPP_class_ values: increasing PC1 of problem behaviors associated with increased potential metabolism of lipids (M), amino sugars (F), other demethylated compounds (M), and near vitamin K2-vitamin B12 compounds (F); but decreased potential metabolism of carbohydrates (M). For CPP_density_ values, patterns were found in males only: increasing PC1 of problem behaviors associated with increased potential metabolism of vitamin B6 (M) and vitamin B9 (M); but decreased potential metabolism of acetate (M), vitamin B12 (M) and glutamate (M). For CPP_ASALR_ values, the comparison was made between groups of high PC1 versus low PC1 of problem behaviors (for consistent display with other case studies in Fig. 3). High PC1 values associated with increased potential metabolism of other methylated compounds (F); and decreased potential metabolism of carbohydrates (M). Weighted mean vK-coordinates between high PC1 and low PC1 of problem behaviors showed a difference in beta-dispersion within condensed aromatics for females only. To explore this dataset in more detail, we identified differentially abundant functions and vK-coordinates (i.e., aggregated functional relative abundances via bioenergetic mapping) in high PC1 versus low PC1 subjects, separately within each sex. In females, 22 differentially abundant functions, compared to only 2-vK coordinates (with 3 corresponding functions), were identified (Figs. S22, S24, Table S18, S20). In males, 6 functions compared to 2 vK-coordinates (with 8 corresponding functions) were identified (Figs. S23, S25, Table S19, S21). Not all functions could be mapped into vK coordinate space. Interestingly, there was no overlap in functions identified directly versus indirectly (from aggregation into vK-coordinates). This means that differential abundance analysis using vK-coordinates can provide entirely different focal areas for investigation compared to the standard function-level analysis. From the vK-coordinate level analysis, high PC1 (compared to low PC1) females exhibited increased potential metabolism of Uridine phosphorylase (EC 2.4.2.3) (fxn_14491) involved in pyrimidine conversions; and decreased aldehyde lyases dihydroneopterin phosphate phosphatase and dihydroneopterin aldolase (EC 4.1.2.25) (fxn_12938; fxn_12942). High PC1 males exhibited increased phosphoenolpyruvate carboxykinase (GTP) (EC 4.1.1.32) (fxn_2926) associated with pyruvate metabolism; and also increases in multiple functions (fxn_821; fxn_2958; fxn_2973; fxn_12703; fxn_12705; fxn_12786; fxn_12788) all involving alcohol dehydrogenase (EC 1.1.1.1) and acetaldehyde dehydrogenase (EC 1.2.1.10), with or without pyruvate-formate-lyase deactivase – involved in degradation of aromatic compounds, biphenyl, tryptophan, and pyruvate metabolism. Results and visualizations for the standard function-level analysis are included for comparison but are not discussed further.

### 3.2. Plant-soil systems

We found remarkable consistency in many observed patterns across the environmental soil case studies. Results reported here are for relative trends with increasing ecosystem maturity (i.e., older revegetation and natural samples) in CPP_class_ (Figs. S26, S30, S33, Table S3), CPP_density_ (Figs. S27-S28, S31-S32, S34-35, Table S4), and CPP_ASALR_ (Figure 5, Table S5). Below, we highlight trends found in at least two of three case studies.

**Fig. 5.**
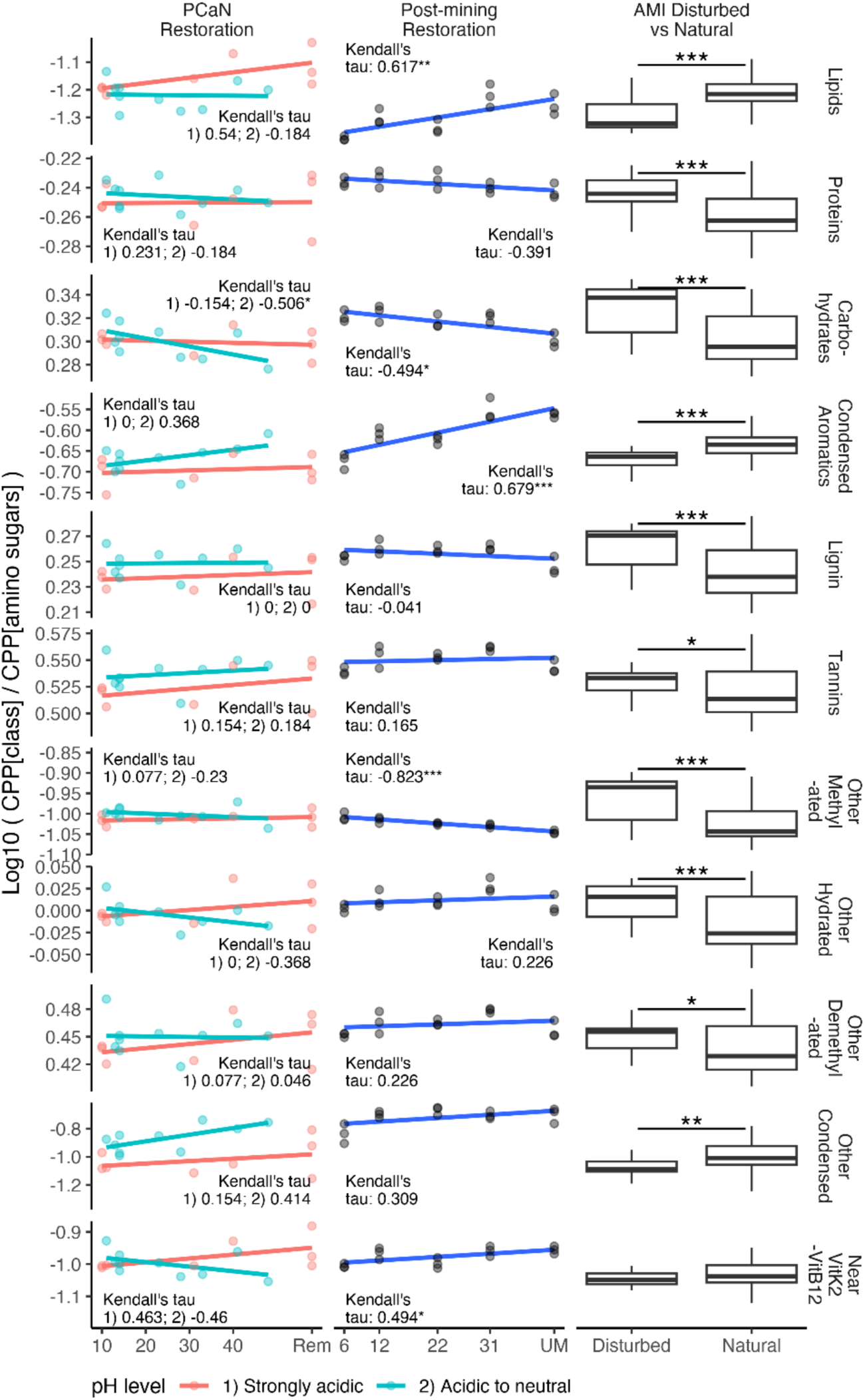
Amino sugar-adjusted log ratio compound processing potential (CPP_ASALR_), representing putative microbiota activity-normalized values, from People Cities and Nature (PCaN) urban restoration, post-mining restoration soils, and Australian Microbiome Initiative (AMI) disturbed versus natural soils. Linear trends were used for visualization purposes. However, Kendall’s tau correlation tests (suited to ordinal data) were applied. Sample sizes are detailed in Table 1.

In CPP_class_ data we observed: increased potential metabolism of lipids (post-mining, AMI) and condensed aromatics (post-mining, AMI); but decreased potential metabolism of proteins (post-mining, AMI), carbohydrates (post-mining, AMI, with marginal indications in both PCaN soil groups), lignin (post-mining, AMI), other methylated compounds (post-mining, AMI), and other hydrated compounds (AMI, PCaN acidic-neutral soils). Mixed or isolated results included potential metabolism: either decreased (AMI) or increased (PCaN strongly acidic soils) for tannins; decreased for other demethylated compounds (AMI); increased for other condensed compounds (AMI); and decreased (post-mining, PCaN acidic-neutral soils) or increased (PCaN strongly acidic soils) for near vitamin K2-vitamin B12 compounds. Total CPP_class_ compounds appeared to be less well characterized and mapped to functional reactions in AMI natural compared to disturbed samples, but more well characterized in older revegetation (compared to younger revegetation) within the PCaN strongly acidic soils.

CPP_ASALR_ results reinforced many patterns observed in the CPP_class_ data: we observed increased potential metabolism of lipids (post-mining, AMI, marginal in PCaN strongly acidic soils) and condensed aromatics (post-mining, AMI); but decreased potential metabolism of proteins (AMI, marginal in post-mining), carbohydrates (post-mining, AMI, PCaN acidic-neutral soils), and other methylated compounds (post-mining, AMI). Isolated results included, potential metabolism: increased for other condensed compounds (AMI); but decreased for lignin (AMI), tannins (AMI), other hydrated compounds (AMI), and other demethylated compounds (AMI). Differing from CPP_class_ results, for CPP_ASALR_ potential metabolism of near vitamin K2-vitamin B12 compounds increased (post-mining).

CPP_density_ results were also quite consistent across case studies: we observed increased potential metabolism of vitamin B9 (AMI, marginal in post-mining and PCaN strongly acidic soils) and vitamin K2 (post-mining, AMI, marginal in PCaN strongly acidic); but decreased potential metabolism of acetate (post-mining, AMI, PCaN acidic-neutral soils), propionate (AMI), vitamin B2 (post-mining, marginal in PCaN strongly acidic soils), vitamin B12 (post-mining, AMI, PCaN strongly acidic soils), vitamin B6 (post-mining, AMI), glutamate (post-mining, AMI, PCaN acidic-neutral soils), and pyruvate (AMI). While butyrate CPP_density_ values were low and indistinguishable across all samples using 0.05 vK unit radial buffers, for an example comparison we tested near butyrate CPP_density_ using a larger 0.1 vK unit buffer in the AMI soils and found increased levels in natural compared to disturbed soils (Fig. S36).

Post-mining and AMI samples displayed significant and mostly consistent directional shifts in weighted mean vK-coordinate centroids across all compound classes (vK mapping zones) considered, while PCaN acidic to neutral soils also displayed shifts within carbohydrates and condensed aromatics (Fig. 6, Fig. S29, Tables S13-S16). Ecosystem maturity (/quality) explained 60–92% of the variation in weighted mean vK-coordinates in post-mining soils (Fig. 6a, Table S15). The following general patterns emerged with increasing ecosystem maturity: lipids became more reduced (lower oxygen content), proteins became more hydrated, amino sugars became more dehydrogenated or condensed, carbohydrates became more condensed (post-mining, PCaN acidic-neutral) or reduced (AMI), condensed aromatics became more condensed, lignin showed mixed trends (dehydrogenation in post-mining, reduction in AMI), and tannins became more demethylated. Except for carbohydrates and amino sugars, these trends largely represented an outward extension of sample profile mapping into vK coordinate space with older ecosystems.

**Fig. 6.**
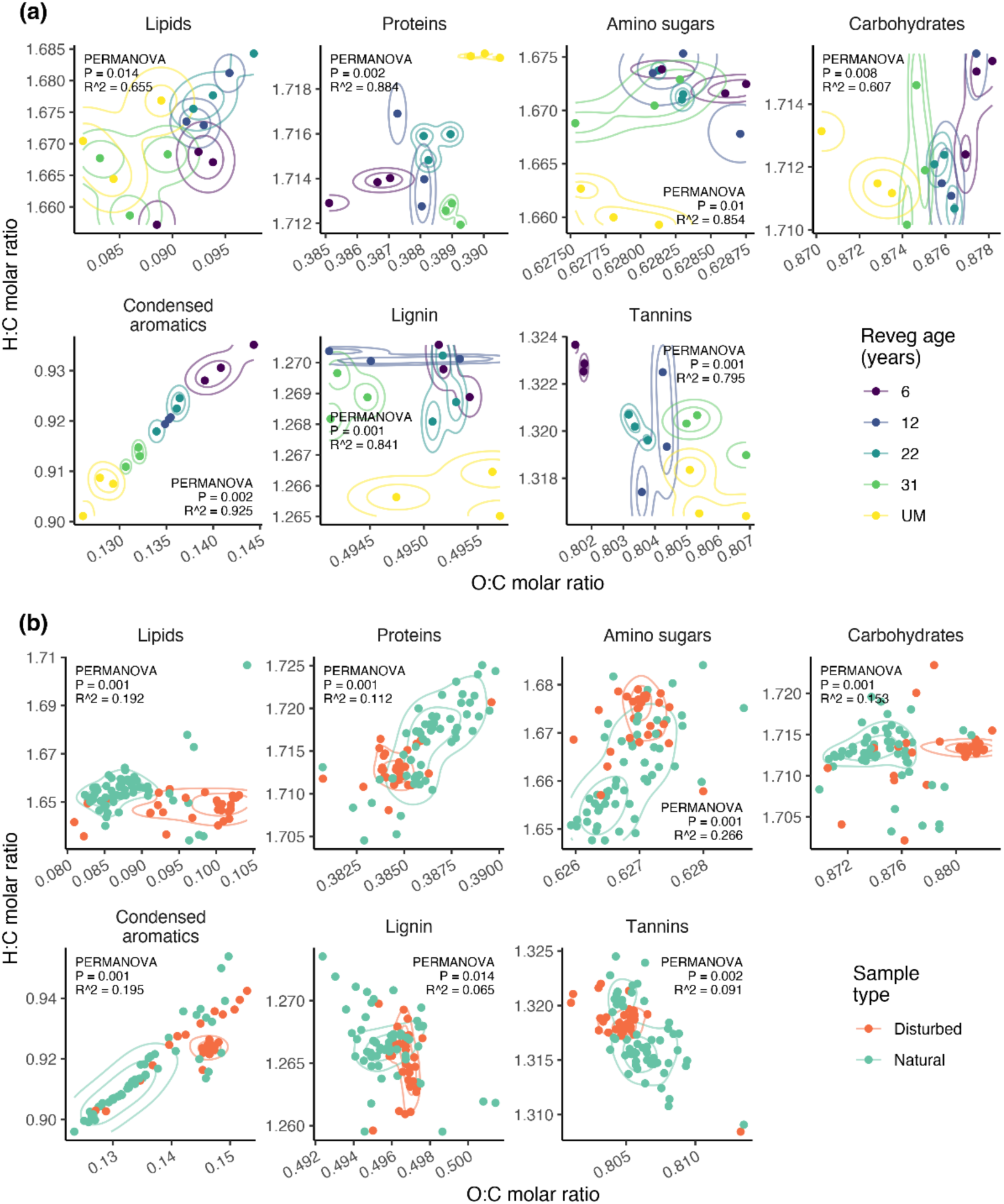
Weighted-mean van Krevelen coordinates within select compound classes display significant shifts with maturity of plant-soil ecosystems. Patterns are from (a) post-mining forest ecosystem restoration soil samples (n = age-based groups of 3), and (b) AMI disturbed (n = 29) vs natural (n = 55) soil samples (b). PERMANOVA and beta-dispersion results are in Table S15-S16.

### 3.3. Consistency in bioenergetic ‘topography’ mapping

CPP_density_ plots visualizing local polynomial regression fitting (‘loess’) smoothed vK mapping profiles for increasing radii buffer areas displayed striking consistency of form within each compound class, with alternative expressions in either gut (Figs. S5, S9, S13, S19) or soils (Figs. S27, S31, S34).

## 4. Discussion

We show a meaningful bioenergetic basis to the development and differentiation of community-scale microbiomes. Our CPP metrics quantified putative shifts in a microbial community’s relative proficiency to process different types of compounds across gradients of human- and environmental-health. We found significant patterns of CPP association at varying resolutions: within major classes of compounds, near focus biomolecules, and for unique vK coordinates. In the environmental soil metagenomes, we observed surprising consistency in CPP profiles, which suggests they may link to plant-soil system conditions in coherent and predictable ways. Our findings align with the notion that microbiota are shaped by the bioenergetic status of prevailing substrates and micro-environments, and this information is simultaneously recorded in their metagenomes. From a methodological perspective, our compound-focused bioenergetic mapping approach demonstrates new pathways for assessing and interpreting microbial systems, capable of supporting ongoing efforts to define healthy microbiomes. For example, aggregating functions via vK-coordinates can provide entirely different focal areas for investigation compared to standard function-level analyses.

### 4.1. Patterns in human health

#### 4.1.1. Atherosclerotic cardiovascular disease (ACVD)

We found strong links in both sexes between ACVD and increased potential gut microbial metabolism of lipids, condensed aromatics, lignin, other hydrated compounds, vitamin B9 and vitamin K2; and decreased potential metabolism of carbohydrates and glutamate. Globally, ACVD is the leading cause of death, with established treatment involving lipid-lowering statins, while human gene editing approaches to disrupt lipid processing are being trialed (Li and Wu, 2022; Vaduganathan et al., 2022).

Links between high-fat diets and ACVD are well established (Law, 2000). Gut microbiota can contribute to ACVD by metabolizing the dietary lipid phosphatidylcholine, with subsequent production of harmful trimethylamine oxide (Albenberg and Wu, 2014). Polycyclic aromatic hydrocarbons are known risk factors in ACVD (Burstyn et al., 2005; Mallah et al., 2021). Lignin (different to lignan) is a complex and ubiquitous structural plant polymer, considered predominantly insoluble fiber, with content ranging from 1-2 g/100g in vegetables, fruits and cereals, up to 30-40 g/100g in nut shells and stone fruit kernels (Tao et al., 2020). Lignin inhibits the enzymatic activity of α-glucosidase, delaying carbohydrate digestion and absorption, with potential for low post-meal blood sugar levels (Chen et al., 2015). Acute hypoglycemia (low blood sugar) can trigger cardiac events (Frier et al., 2011), with greater adverse risks in subjects with significant comorbidities (e.g., T2D, ACVD). Our finding for decreased carbohydrates in ACVD might align with these impacts on blood sugar. Alternatively, we speculate that result might be symptomatic of a more Westernized diet (high in animal protein, sugar, starch, and fat – and lower in carbohydrate content than a plant-rich diet; Albenberg and Wu, 2014).

Excessive vitamin B9 (folate) is associated with ACVD risk via a non-linear u-shaped dose-response relationship (Xu et al., 2023). Non-linear u-shaped dose-response relationships are common in biological systems (i.e., hormesis, or deficiency-sufficiency-toxicity) (Calabrese et al., 2007). Similarly, our results linking increased vitamin K2 with ACVD appear contrary to recent opinion (Essa et al., 2021), although non-linear u-shaped dose-responses have also been observed (Bellinge et al., 2021). Vitamin K2 is commonly found in fermented foods which are less common in Western diets (McFarlin et al., 2017). Here, the ACVD case study was based on Chinese subjects whose diets potentially contained higher quantities of fermented foods including vitamin K2. Possibly, these subjects were more susceptible to adverse effects if excessive levels of vitamin K2 were reached.

Our results linking reduced glutamate with ACVD appear contrary to findings from large US cohort studies which found higher glutamate levels, and lower glutamine:glutamate ratios, correlated with increased ACVD risk (Ma et al., 2017; Zheng et al., 2016). Dietary proteins are a major source of glutamate (Loï and Cynober, 2022). However, some ethnic populations may have inadequate protein in their diets (Liu et al., 2013). Interestingly, in female adults from rural western China with inadequate (and largely plant-derived) protein intake, increasing animal protein associated with reduced risk of hypertension (Liu et al., 2013). Together, these observations suggest that a u-shaped dose-response may also operate for animal-based proteins, glutamate, and ACVD risk.

#### 4.1.2. Colorectal cancer

In the colorectal cancer subjects (from France) we found consistent associations across CPP metrics in both sexes with increased potential metabolism of amino sugars, vitamin B12 and B9; and decreased potential metabolism of proteins and tannins. Colon cancer has been associated with low dietary fiber, low fruit and vegetable consumption, and high red meat consumption (Law, 2000). Our results were consistent with reports for anticancer activity, including protective effects against colorectal cancer, from some tannins or polyphenols (e.g., components in green and black tea, resveratrol in red wine and grapes) (Alam et al., 2018). Unfortunately, our data did not distinguish between animal- and plant-based protein. However, red meat is widely consumed in France, with 41% of males and 24% of females consuming above guideline levels (Mota et al., 2021). Amino sugars are sugar molecules with at least one hydroxyl group substituted by an amino group. In biological systems, they are formed by catalytic activity acting on amino acids (glutamate, glutamine – building blocks of protein) to transfer an amino functionality to a sugar phosphate or sugar nucleotide (Skarbek and Milewska, 2016). Therefore, both glucose (sugar) and amino acids contribute to amino sugar formation. Meanwhile, metabolism of both glucose and amino acids plays a key role in colorectal cancer development (Zhang et al., 2023). Possibly, our finding of increased potential metabolism of amino sugars with colorectal cancer may reflect dysregulated activity of glucose and amino acids with the product of their interaction (amino sugars) recorded by the gut microbiome. Consistent with our findings, vitamin B9 (folate or folic acid) and vitamin B12 supplementation have been associated with increased risk of colorectal cancer (Oliai Araghi et al., 2019).

#### 4.1.3. Type 2 diabetes (T2D)

In the T2D case study, our lack of clear findings linked to major compound classes or focus biomolecules seems consistent with reports that T2D is a complex, multifaceted, highly heterogeneous polygenic disease with uncertain etiology (Pearson, 2019). We found untreated (Met-) T2D exhibited an anomalous and complex CPP network, including a high number of negative correlations (indicating negative feedbacks) ranging widely across vK coordinate space (i.e., covering a spectrum of compounds and bioenergetic status). Diagnosis of T2D is based on elevated blood glucose, primarily arising from insulin resistance and inadequate insulin secretion (Forslund et al., 2015). However, a clear diagnostic test for T2D is lacking, except by exclusion of other causes (Pearson, 2019). A range of factors including genetics, dietary habits, sedentary lifestyle, and gut microbiota are involved in disease development (Forslund et al., 2015). Possibly, with more detailed examination, diagnostic relationships (e.g., correlations, ratios) might be uncovered in relative abundance patterns of compound-associated vK-coordinates underpinning the anomalous T2D Met-network.

#### 4.1.4. Problem behaviors

In the problem behavior case study, different compound associations were observed in female and male children. Increased problem behavior (higher PC1) in females associated with increased potential metabolism of amino sugars, other methylated compounds, and near vitamin K2-vitamin B12 compounds. From differential abundance analysis, high PC1 females exhibited decreased dihydroneopterin aldolase—an enzyme involved in converting dihydroneopterin (a molecule involved in folate biosynthesis) into other compounds (Deng et al., 2000). Excessive serum levels of dihydroneopterin have been associated with major depression (Kusunoki et al., 1999). Possibly, in case study subjects, reduced levels of dihydroneopterin-degrading enzyme have promoted accumulation of dihydroneopterin in association with problem behaviors. High PC1 females also exhibited increased uridine phosphorylase, an enzyme involved in pyrimidine metabolism that converts uridine to uracil (Cao and Pizzorno, 2004), therefore possibly degrading uridine levels in those subjects. Uridine is linked to energy metabolism and glutamate-mediated excitatory neurotransmission in the brain, and supplemental uridine treatments have been used to reduce depressive symptoms in adolescents (Douglas G. Kondo, 2011).

In males, increased PC1 associated with increased potential metabolism of lipids, vitamins B6 and B9; and decreased potential metabolism of carbohydrates, acetate, vitamin B12, and glutamate. Vitamins B12 (cobalamin) and B9 (folate) are recognized precursors involved in forming key neurotransmitters dopamine, noradrenaline (norepinephrine), and serotonin (Hutto, 1997). These three neurotransmitters occur in the vicinity of vitamin B12 and K2 in vK coordinate space. Vitamin B12 deficiency has been associated with depressive disorders in older subjects (Henning Tiemeier et al., 2002). Glutamate is the major excitatory neurotransmitter of the healthy mammalian brain, and an abundant free amino acid important in multiple metabolic pathways, which requires regulation at optimal levels in extracellular fluids (Zhou and Danbolt, 2014). High PC1 males also exhibited increased phosphoenolpyruvate carboxykinase— an enzyme involved in cataplerosis, or removal of intermediate 4- and 5-carbon compounds from the TCA cycle (Owen et al., 2002; Yang et al., 2009). These intermediates are removed because they cannot be fully oxidized for energy metabolism within the TCA cycle, but are converted elsewhere to glucose, fatty acids or amino acids (Owen et al., 2002). High PC1 males also exhibited increases in alcohol dehydrogenase, acetaldehyde dehydrogenase, and pyruvate-formate-lyase deactivase, variously involved in pyruvate metabolism, degradation of aromatics and biphenyl, and tryptophan catabolism. Key processes of energy metabolism involving glucose, lipids, protein and the TCA cycle (via keystone molecules pyruvate, acetyl-CoA, and glutamate) have been implicated in major depressive disorder, although precise pathways of pathogenesis are still unclear (Gu et al., 2021).

### 4.2. Patterns in plant-soil systems

Our findings point to generalizable patterns with older ecosystems for increasing microbial CPP associated with lipids, condensed aromatics, vitamin B9, and vitamin K2; and decreasing CPP associated with proteins, carbohydrates, lignin, other methylated compounds, other hydrated compounds, acetate, vitamin B12, vitamin B6, and glutamate. Drivers of shifting CPP in soils are expected to include changing: (1) composition of biota and biotic materials including plants, organic debris, and re-assembly of invertebrate and microbial communities, and (2) soil abiotic conditions due to plant-soil feedbacks (e.g., pH, nutrients, organic carbon content, temperature, moisture regime) (van der Putten et al., 2016; Wardle et al., 2004). This includes macro-environmental influences with development of vegetation structure and canopy cover (e.g., shading, rainfall interception, altered drainage). CPP values also likely reflect a dynamic balance between resource availability and use by microbiota.

For example, we might expect greater accumulation of lignin in soils of older ecosystems due to plant inputs such as dead roots, bark, leaf litter, and other structural plant residues. However, we observed reduced CPP for lignin in these sample types. Fungi are major lignin degraders (Janusz et al., 2017) and fungal communities vary with ecosystem disturbance and abiotic conditions (Rodriguez-Ramos et al., 2021). Interestingly, our results were counter to expectations for elevated fungal decomposition of lignin in older ecosystems. Reforestation with native mixed-species can produce higher levels of recalcitrant soil organic matter (Cunningham et al., 2015) (e.g., humic acid which is hard to decompose and maps to lignin in vK space). Our CPP metrics are relative and compositional (based on functional relative abundances summing to a maximum of 100%), so it may be that in relative terms, the metabolic foci of microbiota are shifted to processing other materials. Or possibly, structural plant materials may be more accessible for degradation in disturbed (e.g., agricultural) soil environments, depending on plant residue management, nutrient availability and other factors.

We expect some CPP quantities are driven primarily by plant material inputs. For example, soils from more mature ecosystems in temperate climates, represented in samples from AMI and post-mining (in the Appalachian Plateau, southwestern Virginia USA; Avera et al., 2015; Sun and Badgley, 2019), displayed a positive relationship with CPP for lipids and condensed aromatics. These two compound classes are represented in plant-based essential oils and volatile, aromatic organic compounds. Oils are found in high densities in much of the fire-adapted Australian flora (unlike New Zealand flora) (Bowman et al., 2012). Increased CPP for lipids might also arise due to increased density of energy storage linked to primary production, or more active plant signaling in response to abiotic stress (Hou et al., 2016). High levels of lipids and condensed aromatics in mature ecosystem soils could also be a result of increasing plant investment into defensive mechanisms via antimicrobial essential oils (Hammer et al., 1999), and volatile and aromatic secondary defense compounds induced by herbivory (typically by invertebrates) (Erb and Kliebenstein, 2020).

Shifting weighted mean vK coordinates across many compound classes (in AMI and post-mining) suggests broad changes in the composition of microbial substrates with more mature ecosystems. The changing composition of the microbiota itself may contribute to this. Carbohydrates are of interest due to the potential contribution of plant-based material to human diet, and CPP_class_ values for carbohydrates were consistently assigned the largest sum of functional relative abundances in the human gut samples. With more mature ecosystems, CPP for carbohydrates decreased in relative terms, but weighted vK coordinates suggest carbohydrate CPP shifts towards favoring processing materials with reduced oxygen content per unit of carbon. There is likely to be global variation in environmental soil CPP driven by soil abiotic factors and changing biota (vegetation and animals), previously outlined.

### 4.3. Potential environment-human health links

This work opens new avenues for investigating environment-human health connections as environments vary in their production of human health-associated compounds. Moreover, varying environmental microbiota exposures may supply modulating CPP profiles for colonizing or transient impacts to human microbiomes (e.g., skin, airway, gut), which are intimately linked to our health. We show CPP patterns imprinted in environmental soil metagenomes are linked with the maturity (or quality) of plant-soil systems and abiotic factors such as soil pH. We demonstrate that CPP ‘landscapes’ (or the range of meta-compounds) involved in functions that can be performed by soil and gut metagenomes show a large degree of overlap with similar distributions of functional relative abundances across vK space (Fig. S2). We also show a similar spectrum of CPP measures that are impacted by ecosystem quality (e.g., increased potential metabolism of lipids, condensed aromatics, vitamin B9, vitamin K2; and decreased potential metabolism of proteins, carbohydrates, lignin, acetate, vitamin B12, vitamin B6, and glutamate) are also variously linked to human health and disease. However, we urge caution in attempting to directly translate CPP trends in plant-soil environments to infer possible implications for human health. We stress that non-linear, u-shaped dose-response relationships, as discussed earlier (Calabrese et al., 2007), are common and relevant in the context of dietary and environmental exposure-human health links. With potential for u-shaped dose-response effects (i.e., deficiency, sufficiency, toxicity), we might expect environmental microbiota CPP to have varying influences in different geographic/cultural populations. Also, the gut represents a more tightly controlled micro-environment (redox, pH, etc.) unsuited to many environmental microbes.

Example evidence for potential environment-human transfer of microbial CPP comes from Endomicrobia species found in oral microbiota of indigenous peoples from central Australia (Handsley-Davis et al., 2022). Endomicrobia species provide energetic advantage for cellulose digestion in the guts of termites and wood-eating insects—and transfer to humans has occurred likely through use of termites and termite mounds in traditional food and medicine (Handsley-Davis et al., 2022). Speculatively, our results suggest that if broad supplementation of human microbiota CPP capacity is required (spanning a range of health-supporting biomolecules), this may require exposure to multiple types of environments. Alternatively, to address particular microbiota-associated disease states, specific environment types may provide more targeted microbiota CPP supplementation. As knowledge builds of the varying proficiency of environmental microbiota to process human health associated compounds in different qualities of ecosystems, we might design environmental interventions or nature prescriptions to modulate human-microbial exposures, thereby advancing microbiota-oriented approaches to human health.

### 4.4. Limitations

There are important limitations in this study. CPP metrics do not measure actual compounds; rather, they quantify conceptual ‘meta-compounds’ or assemblies of elements based on function-level summary mean O:C and H:C ratios consistent with compounds of interest. Quantification occurred via mapping into vK space and aggregating functional relative abundances into major compound classes, near focus biomolecules, or at unique vK coordinates, to assess CPP structural profiles of metagenomes. Conceptually, mean function-level attributes (O:C and H:C ratios) represent a mid-point of chemical transformation mediated by microbiota (i.e., interpreted as ‘X% of functions were involved in processing compound/biomolecule type Y’). Stoichiometry rules determine that reaction inputs and products will have balanced O, H, and C atomic counts. However, different mean O:C and H:C ratios can arise due to uncounted O and H atoms in non-C containing species (O:C, H:C values become undefined). For future work, the CPP mapping algorithm could be readily adjusted to separately target reaction inputs, or products, or individual chemical species.

Our coarse compound classes did not distinguish (for example) plant versus animal proteins or high-fiber versus low-fiber carbohydrates. Finer-resolution vK mapping zones would increase the precision of results. We could not discern CPP differences for butyrate between sample types using 0.05 vK radii. Butyrate is often present at low concentrations in the gut compared to other SCFAs, with rapid consumption by colonocytes (den Besten et al., 2013). The volatility of butyrate (Wagner et al., 2017) may make it susceptible to loss from soils. Our butyrate CPP_density_ profiles spanning large to small vK radii may depict source-sink dynamics. Compound mapping using O, H, and C content enable exhaustive and compartmented mapping within vK coordinate space, however this represents a simplified, imperfect approach. Multi-element compound mapping would offer increased precision (Rivas-Ubach et al., 2018) and may be developed to provide exhaustive and compartmented mapping across a range of compound types. CPP measures used here were relative, not absolute. Mapping of SUPER-FOCUS functions was incomplete (sample functional relative abundances ranged from 52-84%, with means 55-67%). This may be improved with future algorithm refinement.

Further work is required to refine our first-pass ASALR normalization, to examine CPP-disease links in wider geographic/cultural populations, and to explore other potential explanatory variables not considered in our analyses. Like many microbiome studies, our analyses do not permit causal insight to interpret whether increased or decreased CPP may facilitate or follow disease. For example, excessive CPP may produce metabolites at toxic levels, or degrade substrates leading to deficiency. Reduced CPP measures might correspond to dietary deficiencies or suppression of functional pathways due to dysregulated environmental conditions. Nonetheless, observed CPP trends may assist hypothesis-building and prioritizing mechanistic research. We suggest that future work might address these limitations.

## 5. Conclusions

Our assessment of microbial metagenomes using the compound processing potential framework offers a new approach to examine potential overlapping functional capacities between human and environmental microbiomes. Our findings reveal a meaningful bioenergetic basis to the development and differentiation of microbiomes in human health and disease, and with variation in ecosystem quality. We show that ecosystem quality can shape environmental microbiota in seemingly predictable ways, with varying proficiency to process a range of compounds that are associated with human health and disease—as measured by the functional potential relative abundances imprinted in their collective DNA. This work informs the ongoing development of microbiota-oriented approaches in human health, including the potential for nature-based solutions to help address global healthcare challenges.

## Data availability

All data used in this study are available from repositories listed in Table 1. Code used to support this study is available from: https://github.com/liddic/compound_potential

## Acknowledgments

We acknowledge the authors of previously published metagenomics datasets used here which enabled comparison of diverse samples spanning human and environmental health. We acknowledge First Nations Peoples and traditional custodians of the lands from which the case study datasets originate. This work was supported by the People, Cities and Nature research program (Aotearoa New Zealand Ministry of Business, Innovation and Employment, grant UOWX2101), Marsden Fund Fast-Start grant funding (Royal Society Te Apārangi), funding from the German Research Foundation iDiv ([DFG]–FZT 118, 202548816; Ei 862/29-1; Ei 862/31-1; Ei 862/27-1; HE 8266/4-1), and the Australian Research Council (grant numbers LP190100051, LP190100484). We acknowledge the contribution of the Australian Microbiome consortium in the generation of data used in this publication. The Australian Microbiome Initiative is supported by funding from Bioplatforms Australia and the Integrated Marine Observing System (IMOS) through the Australian Government’s National Collaborative Research Infrastructure Strategy (NCRIS), Parks Australia through the Bush Blitz program funded by the Australian Government and BHP, and the CSIRO.

