## Supplementary Information for "Bioenergetic mapping of ‘healthy microbiomes’ via compound processing potential imprinted in gut and soil metagenomes"

**This PDF file includes:**

Supporting Information
Figures S1 to S36
Tables S1 to S21
Supplementary References

**Other supporting materials for this manuscript include the following:**

Code used to support this study is available from:
https://github.com/liddic/compound_potential

#### Supporting Information

##### Case study metagenomics datasets.

Case study data used in the article are described below. For pragmatic reasons some case study datasets comprise a subset of the parent study data based on completeness of metadata, public availability of sequences, and sufficiency of a coherent illustrative dataset within reasonable geographic or environmental bounds (further described below). This approach reflected our focus to examine patterns in compound processing potential (CPP) with human health and disease, and across a spectrum of ecosystem maturity/condition, which could be achieved without re-analyzing entire case study parent datasets. Unless otherwise stated, raw shotgun metagenomics fastq sequence files were downloaded and processed as described in the following section.

Atherosclerotic cardiovascular disease (ACVD) metagenomics data were sourced from (Jie et al., 2017). Here we used a total of 380 samples (210 ACVD, 170 normal healthy) that were both complete with sex metadata and available for public download from the National Centre for Biotechnology Information (NCBI) Sequence Read Archive (SRA, accession PRJEB21528).

Colorectal cancer metagenomics data were sourced from (Zeller et al., 2014, SRA accession PRJEB6070) where we considered their French study population only. Subjects with diagnoses for adenoma (benign noncancerous tumor) were excluded. Sequence reads for one normal healthy male failed to pass the specific QC parameters used in our pipeline, resulting in 60 normal healthy and 53 cancer subjects (113 total samples) used here.

Type 2 diabetes (T2D) metagenomics data were sourced from (Forslund et al., 2015, SRA accession PRJEB1786) where we considered the Swedish female-only cohort samples from the parent study. These 145 total samples comprised 53 subjects with T2D (with and without Metformin treatment) and 92 nondiabetic individuals (43 normal healthy, 49 with impaired glucose tolerance).

Problem behavior metagenomics data were sourced from (Flannery et al., 2020, SRA accession PRJNA496479). Multiple standardized measures of problem behavior in children—comprising CBCL Aggressive Behavior, CBCL Anxious Depressed, CBCL Depressive Problems, CBCL Internalizing Behavior, CBQ Anger Frustration, CBQ Fear, CBQ Inhibitory Control, CBQ Impulsivity, and CBQ Sadness (where CBCL = Child Behavior Checklist, and CBQ = Children's Behavior Checklist; Flannery et al., 2020)—were simultaneously considered via principal components analysis (see Fig. S3). The first principal component (PC1) of problem behaviors was ultimately used as the health response variable. PC1 values correlated with reported measures of depression ($r = 0.86$ in females, $r = 0.72$ in males). Due to incomplete cases (missing data) in three subjects for the behavior measures used to define PC1, only 37 samples (20 females, 17 males) were used, from 40 subjects in the parent study.

People Cities and Nature (PCaN) urban forest restoration metagenomics samples used here represent pilot-scale metagenomics survey data collected in Nov 2019 to Feb 2020 from across Aotearoa New Zealand (Barnes et al., 2020, AGDR project AGDR00045) spanning revegetation ages from 10 to 48 years old ('young' < 15 years, n = 8; 'old' > 15 years, n = 7), as well as remnant sites (n = 3). These data were analyzed in pH-based classes: strongly acidic (n = 8, pH range 3.4 to 4.3), and acidic-neutral soils (n=10, pH range 4.6 to 7). One site (DunSiH, 31 years old) was not included from the parent dataset due to missing pH data. Sample locations are described in (Mitchell, 2022).

Post-mining forest ecosystem restoration data were sourced from (Sun and Badgley, 2019), using the entire parent study dataset. Raw fastq files were not available, so the SUPER-FOCUS analysis proceeded from screened fasta files downloaded directly from MG-RAST. Soil samples were considered to represent ordinal (semi-quantitative) treatment values with revegetation ages of 6, 12, 22 and 31 years, together with unmined (UM) samples.

Soil data representing disturbed versus natural plant-soil systems were sourced from (Bissett et al., 2016; Australian Microbiome Initiative, AMI). On 15 September 2022 we queried available metagenomes on the Bioplatforms Australia AMI Data Portal (https://data.bioplatforms.com/bpa/otu/metagenome) via contextual filters. Specifically, we searched for soil samples (Am Environment = Soil), within the temperate climate zone (Gen Env Feature = Temperate), from surface depths (Depth Upper [m] <= 0), and with clay content (Clay [%]) in the range 7.5–45%, which was selected to avoid either very sandy or very clayey soils. From 104 samples returned, we excluded ambiguous samples with land use/management (Env Local Scale) recorded as '3.2.1 Native/exotic pasture mosaic', '1.1.5 Habitat/species management area', '1.3.4 Rehabilitation', '1.3.3 Residual native cover', '1.1.6 Protected landscape'. The remaining 84 samples were classed into groups of either 'Natural' (n = 55, including Env Local Scale = '1.1 Nature conservation', '1.1.3 National park', and '6.5.1 Marsh/wetland conservation'), or 'Disturbed' (n = 29, including Env Local Scale = '3.3.1 Cereals', '3.3.6 Cotton', '3.4.1 Tree fruits', '3.4.4 Vine fruits', '4.2.3 Irrigated legume/grass mixtures', and '4.5 Irrigated seasonal horticulture'). Land use/management criteria used to define natural versus disturbed soils followed the approach used in (Liddicoat et al., 2019).

**Bioinformatic processing and SUPER-FOCUS functional profiling.**

Scripts detailing the bioinformatic processing and SUPER-FOCUS functional profiling are available at: https://github.com/liddic/compound_potential

Briefly, Linux shell scripts and Python (v3.8.5; https://www.python.org/) scripts were used to download previously published publicly available case study datasets from the NCBI Sequence Read Archive (SRA) (https://www.ncbi.nlm.nih.gov/sra) or MG-RAST (https://www.mg-rast.org/). Linux shell, Snakemake (v5.22.0; Molder et al., 2021) and Python scripts were used for quality assessment using FastQC (v0.11.9; Andrews, 2018) and quality control/trimming using Fastp (v0.23.2; Chen et al., 2018). Using shell scripts and Python scripts, good quality R1 sequences were then used to generate functional relative abundance profiles from each sequence through use of SUPER-FOCUS (v0.0.0; https://github.com/metageni/SUPER-FOCUS; Silva et al., 2015) software, linked to the Diamond sequence aligner (v0.9.19; Buchfink et al., 2021) and version 2 100% identity-clustered reference database (100_v2; https://github.com/metageni/SUPER-FOCUS/issues/66). Where subjects/samples were represented by multiple sequence files, the combined SUPER-FOCUS outputs were normalized so that the total functional relative abundances summed to 100% in each subject/sample. Operations involving sequence downloading, QA/QC, and SUPER-FOCUS functional-profiling were performed on the Flinders University DeepThought high performance computing facility (Flinders_University, 2021). Further data analyses were performed using R (v4.2.2; R_Core_Team,

2022) in the RStudio environment (v2022.12.0+353; https://github.com/rstudio/rstudio). Within R, functional profile data was handled using the microbiome data management framework provided by the R phyloseq package (v1.44.0; McMurdie and Holmes, 2013).

**Compound processing potential (CPP) mapping.**

Further to the Method outlined in the main paper, in each dataset we observed an anomalous number of overlapping van Krevelen (vK) coordinates at O/C = 0.5206427, H/C = 1.400701 (in the Lignin zone) corresponding to the poorly specified SUPER-FOCUS functional output termed 'hypothetical protein' – which triggered a match to multiple reactions in several thousand occasions. Although, typically at the per-sample level these hypothetical protein vK coordinates summed to only 0.045-0.06% functional relative abundance. Due to their poor description these hypothetical proteins were excluded from subsequent analyses.

We mapped the entire ModelSEED Compound database (33,992 compounds) into vK coordinate space to validate the molar ratio assignment subroutine of the CPP mapping algorithm (Fig. S1). Reassuringly, this achieved equivalent distributions to previous vK mapping of metabolite databases (Rivas-Ubach et al., 2018). Similarly, following CPP mapping all case study datasets were visualised as 2-dimensional density distributions of all compound-associated vK coordinates, displaying notable similarities in their coverage of vK space (Fig. S2).

**Data visualization and statistical testing.**

In each case study dataset we performed a standard set of analyses.

**$CPP_{class}$ data.** $CPP_{class}$ values for grouped data (ACVD, colorectal cancer, T2D, AMI) were presented as boxplots and testing for differences between disease and normal sex-based groups, and in AMI disturbed versus natural soils, used t-tests, except when groups had unequal variance in which case the Wilcoxon rank sum test was used. Testing across multiple groups of female-only T2D data first applied Levene's test to confirm common variance, followed by analysis of variance testing and post-hoc Tukey tests; unless we found unequal variances, in which case we used Kruskal-Wallis rank sum testing, followed by post-hoc pairwise Dunn tests (if applicable). In the case of ACVD, data were 95% Winsorized to contain outlying values. For ordinal scale independent variable data (PCaN, post-mining) we tested for trends with $CPP_{class}$ values using Kendall's tau rank-based correlation coefficient, which is suited to assessing ordinal associations (values of 1 or -1 indicate strongest correlations). Problem behavior data were expressed and tested for effects via the first principal component (PC1) of nine covarying behaviours (SI Appendix Fig. S3). Associations between $CPP_{class}$ values and the PC1 of problem behaviors (which are both numeric variables) were assessed using Pearson r correlation tests.

**$CPP_{density}$ data.** $CPP_{density}$ values were visualized at log10-transformed scale (following zero-replacement with half the smallest non-zero value for the respective biomolecule-analysis sub-group) using the geom_smooth() trendline in R ggplot2 package (v3.4.2; Wickham, 2016) with 'loess' (local polynomial regression) smoothing. These trendlines indicate the local 'topography' of vK coordinate space adjacent to the biomolecules of interest. Differences in log10($CPP_{density}$) values were tested only using measures calculated within the closest radius (0.05 vK units) using the same testing approach as described for $CPP_{class}$ data.

**$CPP_{ASALR}$ data.** $CPP_{ASALR}$ values were calculated and displayed as boxplots for ACVD, colorectal cancer, T2D, problem behaviours and AMI data. For $CPP_{ASALR}$ testing, problem behavior PC1 data were split into two groups, i.e., low PC1 < median(PC1), and high PC1 > median(PC1). The remaining PCaN and post-mining soil datasets were visualized using linear trend lines. Pairwise difference tests or Kendall's tau ordinal correlation tests were then applied as above. In T2D $CPP_{ASALR}$ data, testing for differences across multiple groups first applied Levene's test to confirm common variance, followed by analysis of variance testing and post-hoc Tukey tests (if applicable).

**Weighted mean vK coordinates.** Weighted mean compound-associated vK coordinates were plotted to visualize differences and shifting weight of functional relative abundance occurring within $CPP_{class}$ zones, to explore effects linked to health versus disease, and with changes in ecosystem status. These plots were prepared for all case studies. Plots included 2-d kernel density estimation, displaying density distributions via contour lines, using the geom_density_2d() function in R ggplot2. Testing for differences in centroids between sample groups, within each $CPP_{class}$ zone, was performed using permutational multivariate analysis of variance (PERMANOVA), followed by testing for homogeneity of group dispersions. Specifically, these respective tests used the functions adonis2() and betadisper(), both from the R vegan package (v2.6.4; Oksanen et al., 2020).

Additional analyses were performed on a case-by-case basis for illustrative purposes.

**Network analysis.** T2D diagnosis groups showed little effects via coarse-level examination using $CPP_{class}$ and $CPP_{ASALR}$ data, therefore we explored example network analysis using the CPP mapping framework with functional relative abundances aggregated at the level of unique vK coordinates. We performed network analysis using equal sample sizes equivalent to the smallest diagnosis group (T2D Met+, n = 20). Twenty samples each were randomly selected from T2D Met-, IGT, and Normal groups. Correlations between function relative abundances summed at unique compound-associated vK coordinates were determined using the compositionally-aware method of CCLasso (Fang et al., 2015). To avoid emphasizing correlations from sparse and less common vK coordinates, CCLasso correlation matrices were derived using only vK coordinates with non-zero abundances in at least 60% of samples and a minimum of 2% total functional relative abundance summed across samples. Significant correlations ($p \leq 0.05$) only were retained and translated to networks using the R igraph package (v1.4.2; Csardi and Nepusz, 2006), then displayed with nodes (vK coordinates) plotted in vK coordinate space, and displaying positive and negative correlations, and degrees (number of correlations/links between nodes). Testing for differences between networks of the four diagnosis groups was performed by comparison with bootstrap (B = 1000) randomized network probability density distributions. The following network characteristics were assessed: number of vertices (nodes), number of edges (links), edge density, fraction of negative edges, degree centralization, closeness centralization, betweenness centralization, mean distance, and modularity.

**Differential abundance testing.** In the behavior case study, we performed differential abundance testing using ANCOM-BC software (v2.0.1; Lin and Peddada, 2020), based on conventional 'fine-resolution' SUPER-FOCUS functional relative abundance outputs, as well as for data aggregated (summed functional relative abundances) at unique compound-associated vK coordinates, to examine differences in these approaches.

**Focus biomolecules in this study.**

In gut microbiome bioenergetics and chronic metabolic diseases, Daisley et al. (2021 highlight key human health-associated biomolecules (compounds produced by living organisms) found within extracellular resources shared by microbes. Short-chain fatty acids (SCFAs) acetate, propionate, and butyrate benefit host metabolism, intestinal barrier function, systemic anti-inflammatory effects, and contribute up to 10% of daily energy requirements (den Besten et al., 2013). B group vitamins: riboflavin (B2), cobalamin (B12), pyridoxal 5'-phosphate (B6), and folate (B9) are critical in electron transport and represent precursors to a variety of enzyme cofactors essential to the tricarboxylic acid (TCA) cycle, fatty acid oxidation, and other metabolic pathways (Daisley et al., 2021). Menaquinone (Vitamin K2) is a critical electron carrier in bacteria and considered essential in humans for calcium regulation (Daisley et al., 2021). Other keystone health-linked biomolecules include glutamate and pyruvate. Glutamate is the major excitatory neurotransmitter of the healthy mammalian brain, and an abundant free amino acid important in multiple metabolic pathways, which requires regulation at optimal levels in extracellular fluids (Zhou and Danbolt, 2014). Glutamate is sensed luminally in the intestinal mucosa, triggering vagus nerve (gut-brain axis) activity (Torii et al., 2013). Pyruvate is a critical intermediate involved in human energy metabolism, where dysregulation is associated with cancer, heart failure, and neurodegeneration (Gray et al., 2014).

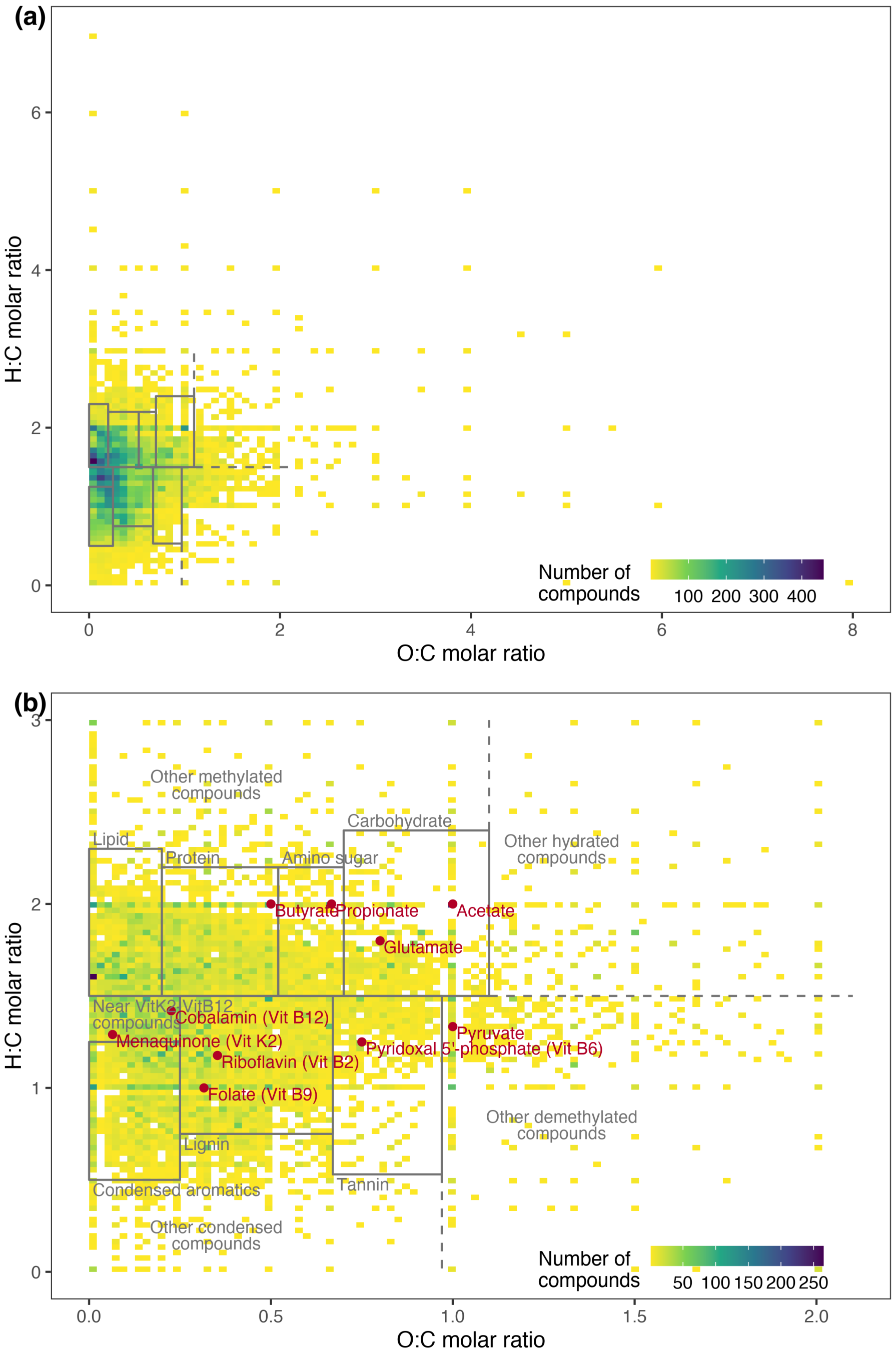


**Fig. S1.** Density map for all ModelSEED database compounds (n = 33,992) mapped to van Krevelen (vK) coordinate space, (a) displaying full-extent; and (b) limited (zoomed-in) extent detailing vK-zones for major compound classes and vK-coordinates for focus biomolecules in this study.

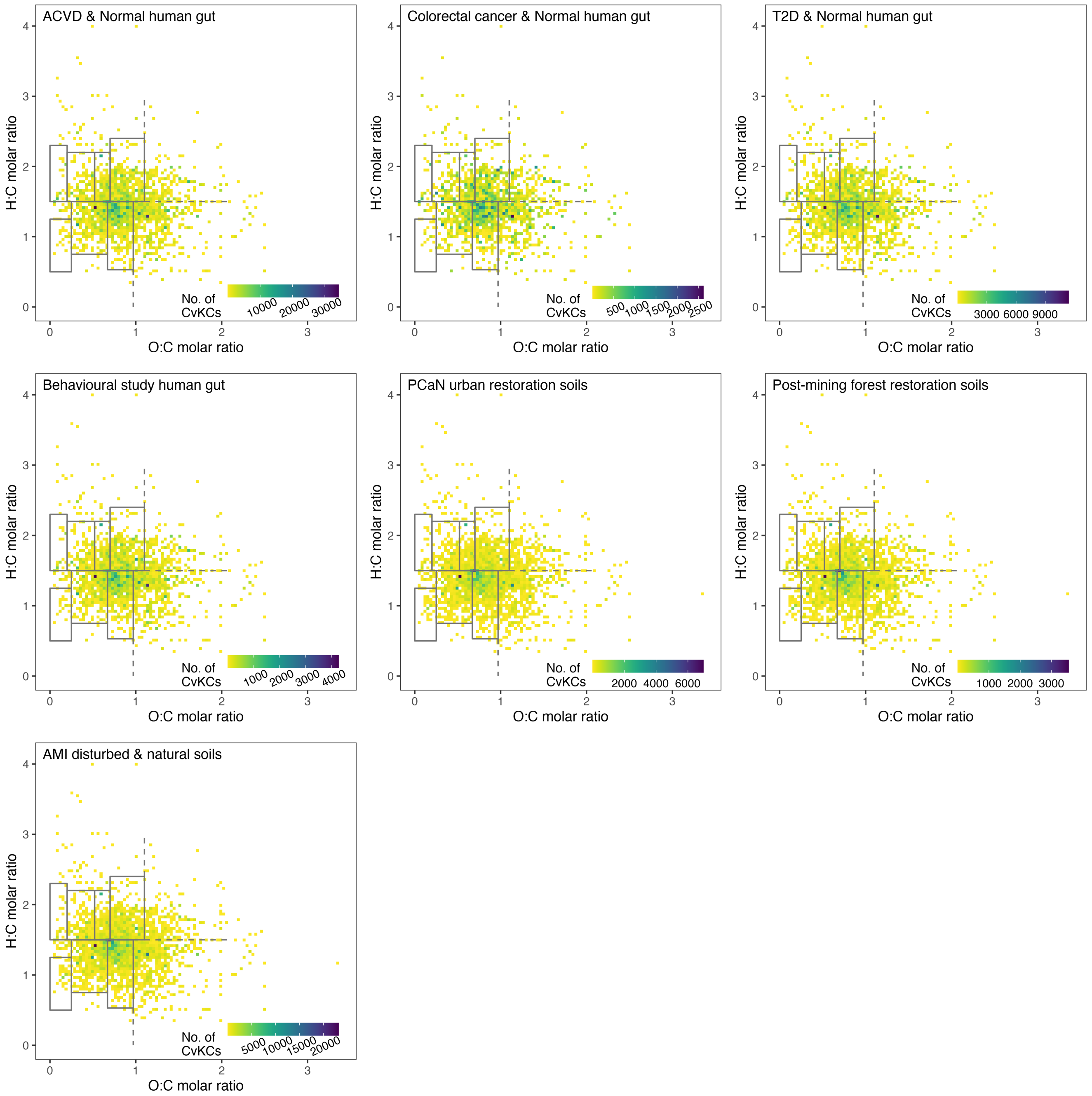


**Fig. S2.** Distribution of entire case study datasets mapped to van Krevelen (vK) coordinate space. Functional relative abundance datasets derived using SUPER-FOCUS software analysis of shotgun metagenomics samples were mapped to functional reaction abundance-weighted mean compound-associated van Krevelan coordinates (CvKCs).

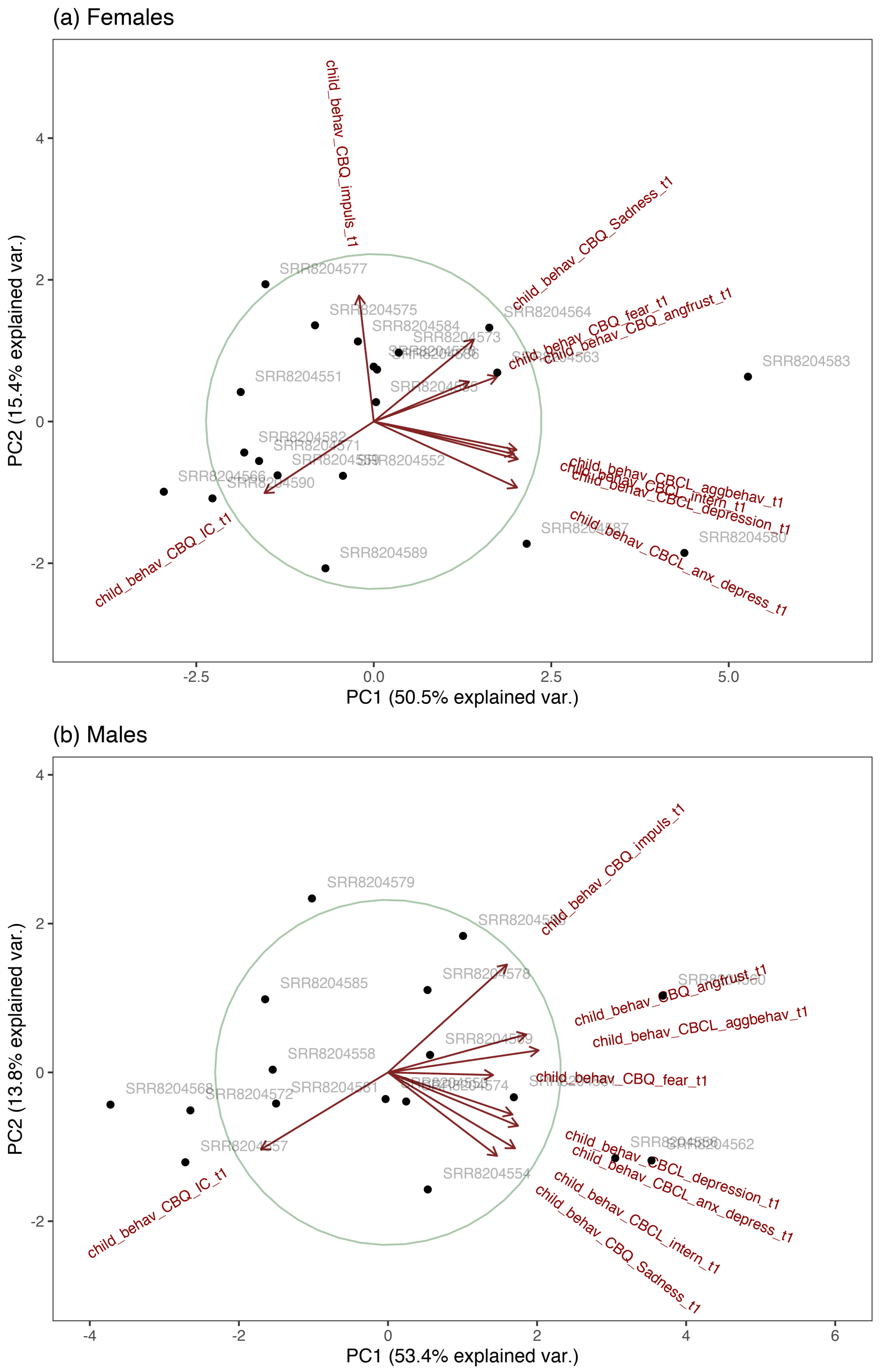


**Fig. S3.** Principal components analysis (PCA) of (a) female (n = 20) and (b) male (n = 17) child problem behaviors from (Flannery et al., 2020). Within each sex, the PCA was based on measures of aggressive behavior, anxious depressed behavior, depressive problems, internalizing behavior, anger frustration behavior, fear, inhibitory control, impulsivity, and sadness. By way of example, measures of depression showed strong to very strong correlation (r = 0.86 in females, r = 0.72 in males) with the first principal component (PC1) of these nine covarying behaviors.

Atherosclerotic Cardiovascular disease (ACVD) case study data (raw sequence data from Jie et al., 2017)

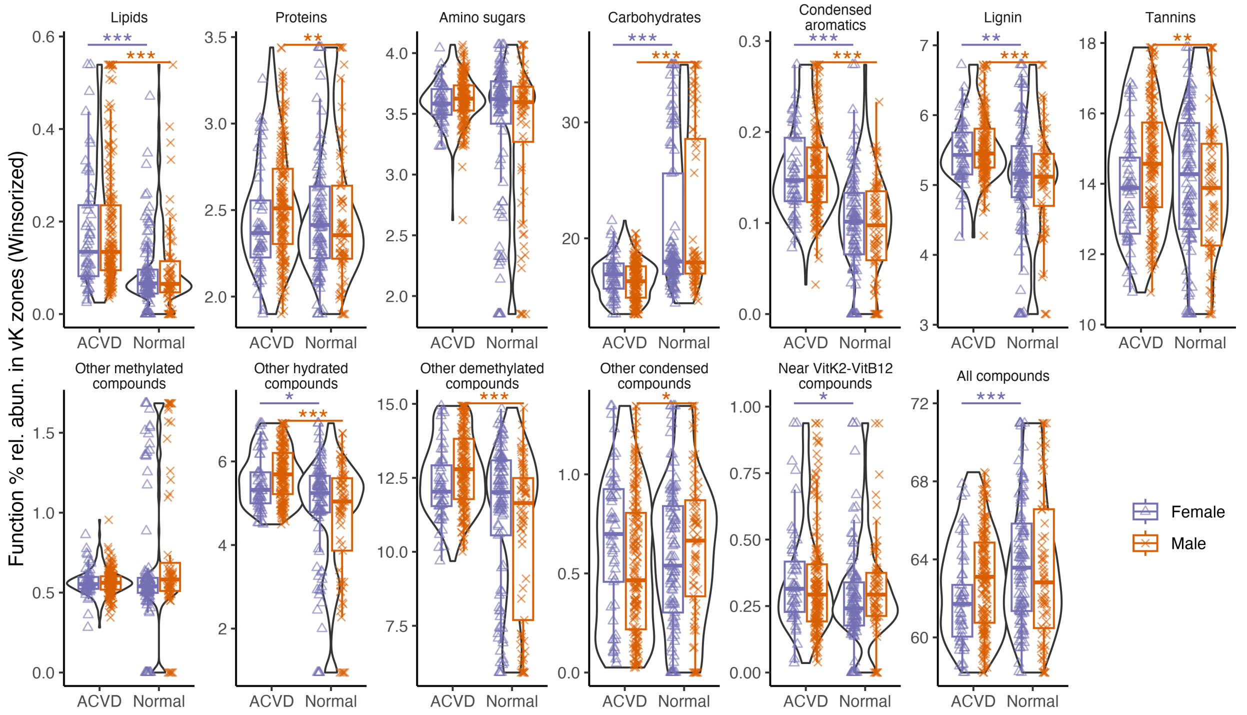


**Fig. S4.** Compound processing potential (CPP$_{class}$) in ACVD and normal subjects across compound classes, based on mapping of % functional relative abundances to compound-associated vK zones (ACVD-females n = 53, ACVD-males n = 157, normal-females n = 101, normal-males n = 69). Data were 95% Winsorized to limit outlying values.

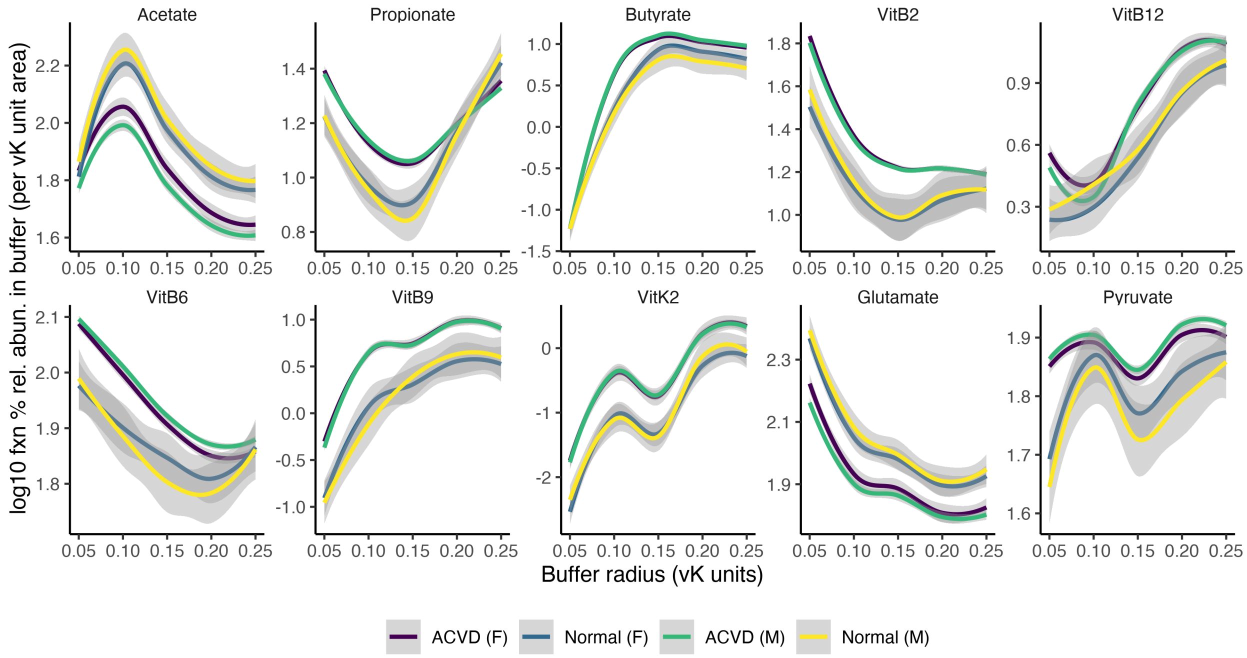


**Fig. S5.** Compound processing potential densities (CPP$_{density}$) in proximity to key health-associated biomolecules, based on log10 % function relative abundance mapped to radial buffers surrounding selected biomolecules, normalized (divided) by buffer area (ACVD-females n = 53, ACVD-males n = 157, normal-females n = 101, normal-males n = 69).

ACVD case study data (continued)

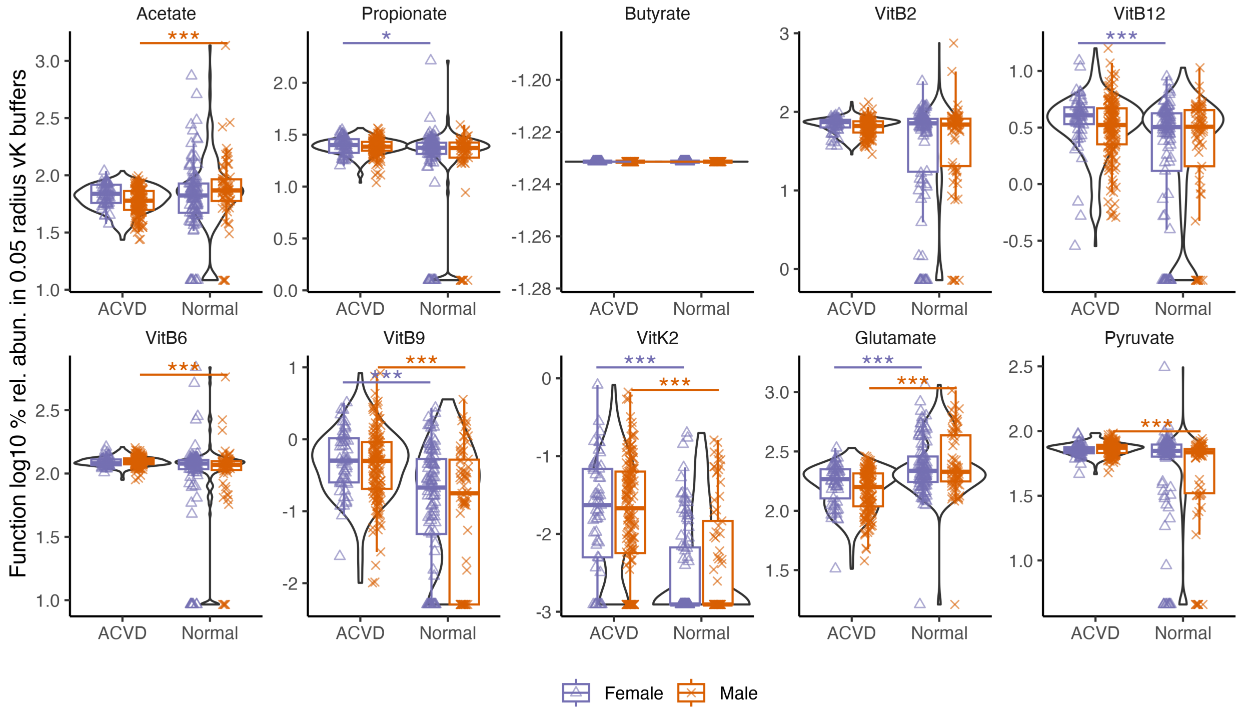


**Fig. S6.** Compound processing potential density ($CPP_{density}$) corresponding to key health-associated biomolecules, based on log10 % functional relative abundance mapped to the smallest radial buffer (0.05 vK units) normalized (divided) by buffer area (ACVD-females n = 53, ACVD-males n = 157, normal-females n = 101, normal-males n = 69).

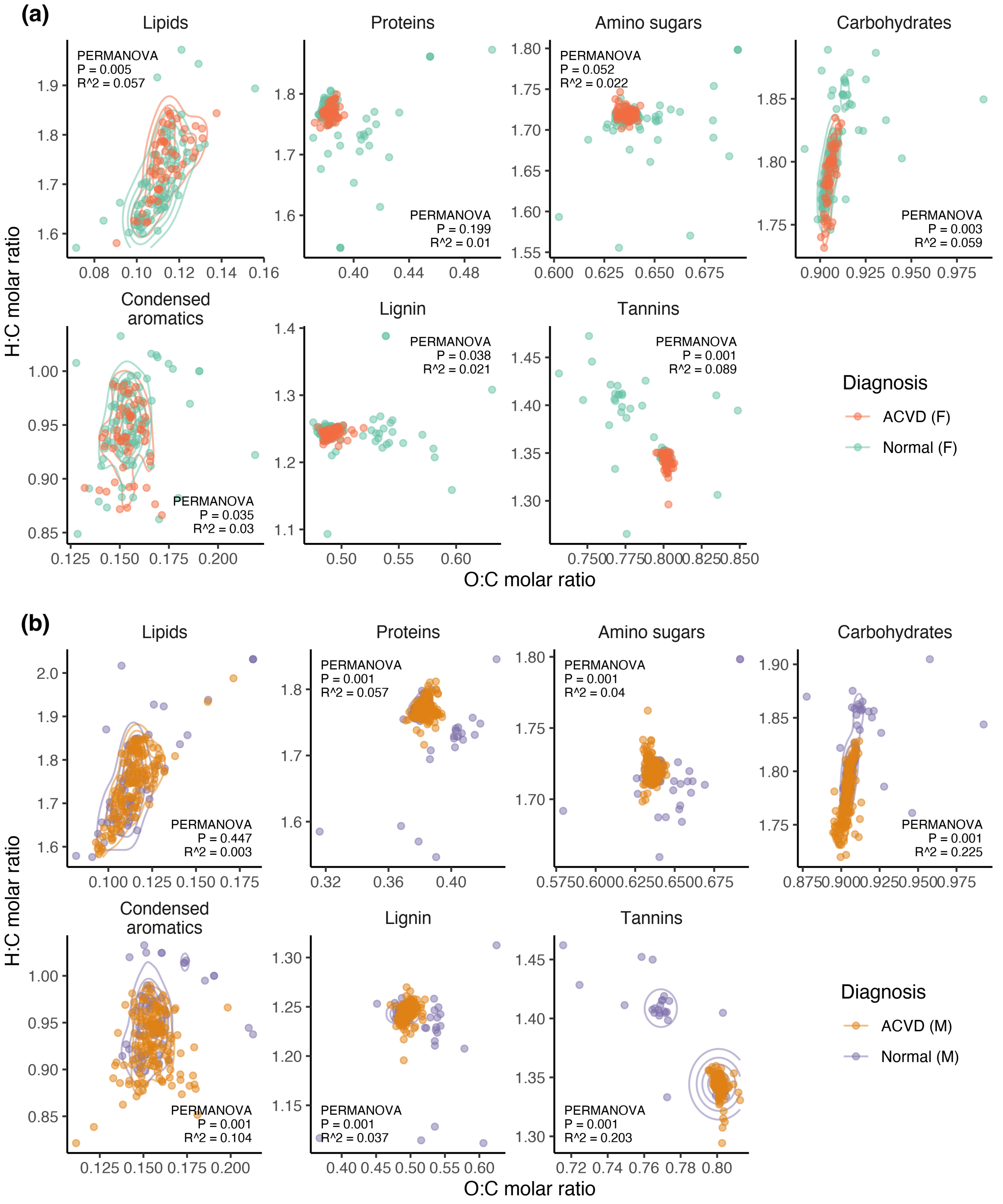


**Fig. S7.** Weighted mean vK coordinates within compound classes in ACVD and normal subjects in (a) females (ACVD n = 53, normal n = 101), and (b) males (ACVD n = 157, normal n = 69).

Colorectal cancer case study data (raw sequence data from Zeller et al., 2014)

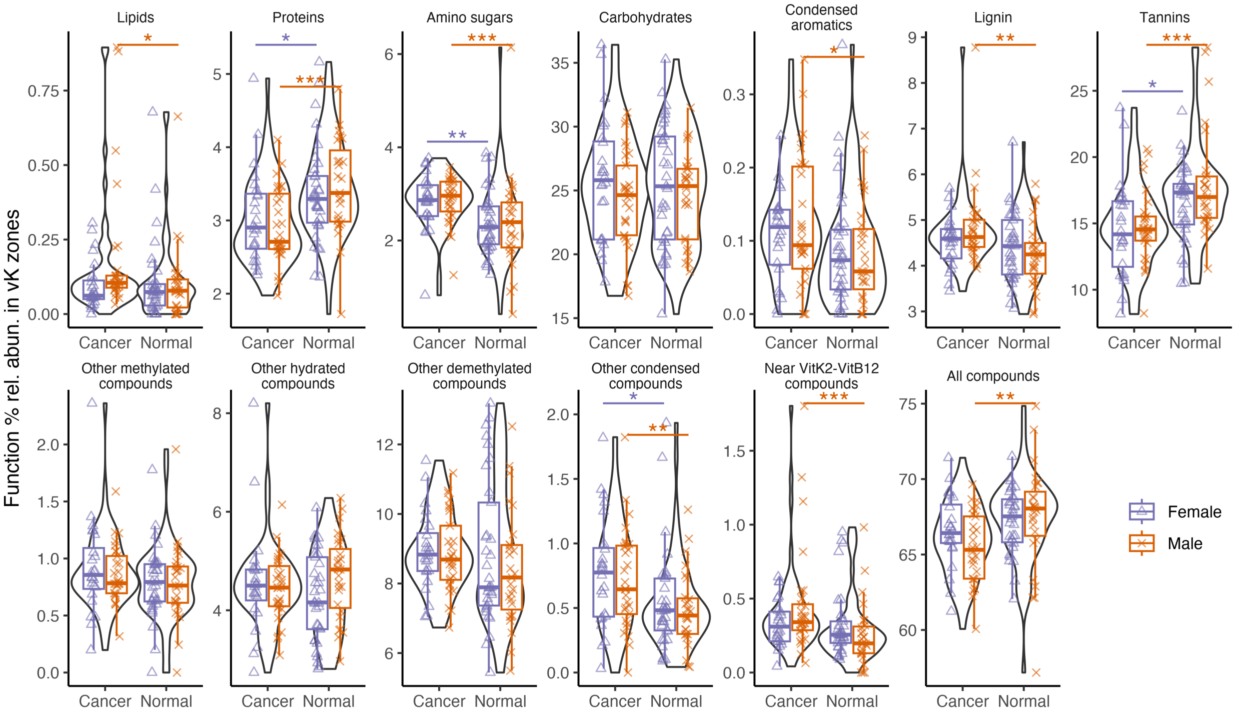


**Fig. S8.** Compound processing potential ($CPP_{class}$) in colorectal cancer and normal subjects across compound classes, based on mapping of % functional relative abundances to compound-associated vK zones (cancer-females n = 24, cancer-males n = 29, normal-females n = 33, normal-males n = 27).

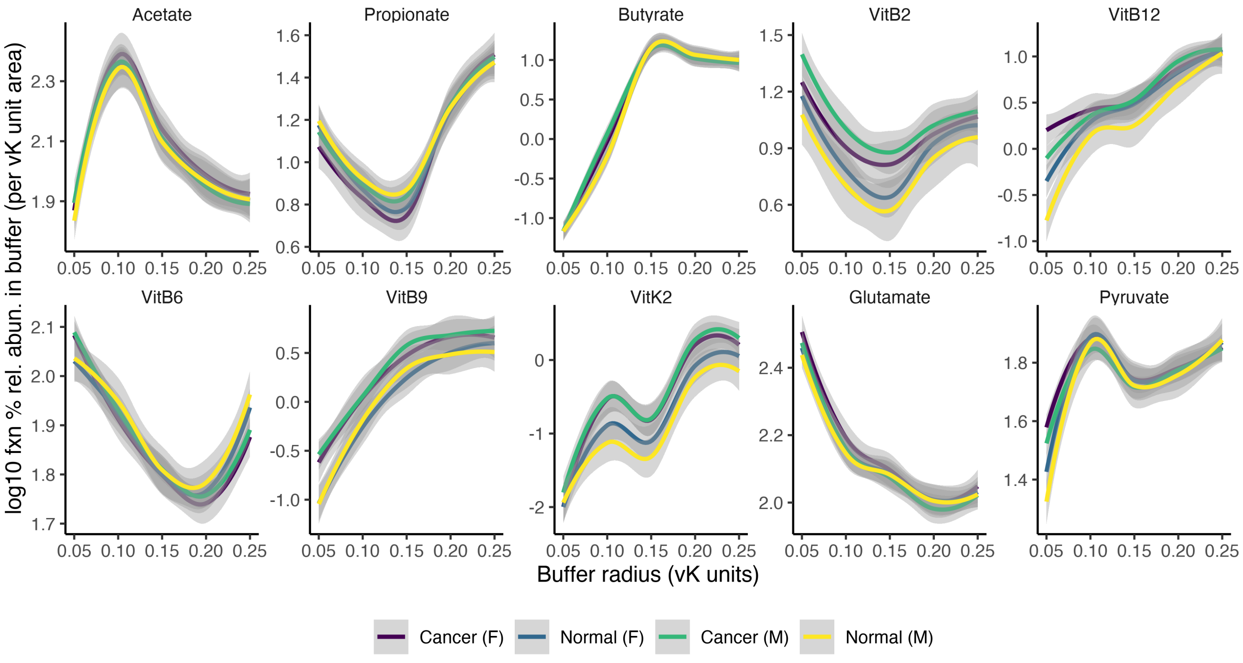


**Fig. S9.** Compound processing potential densities ($CPP_{density}$) in proximity to key health-associated biomolecules, based on log10 % function relative abundance mapped to radial buffers surrounding selected biomolecules, normalized (divided) by buffer area (cancer-females n = 24, cancer-males n = 29, normal-females n = 33, normal-males n = 27).

Colorectal cancer case study data (continued)

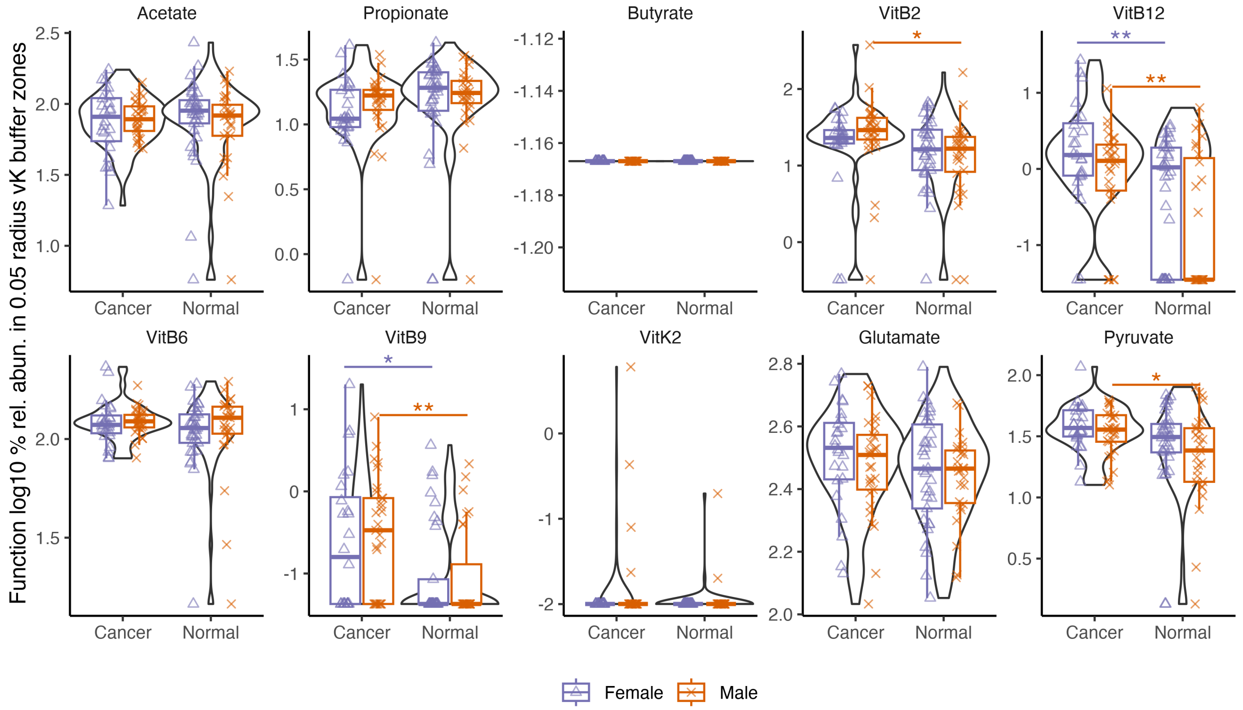


**Fig. S10.** Compound processing potential density ($CPP_{density}$) corresponding to key health-associated biomolecules, based on log10 % functional relative abundance mapped to the smallest radial buffer (0.05 vK units) normalized (divided) by buffer area (cancer-females n = 24, cancer-males n = 29, normal-females n = 33, normal-males n = 27).

Colorectal cancer case study data (continued)

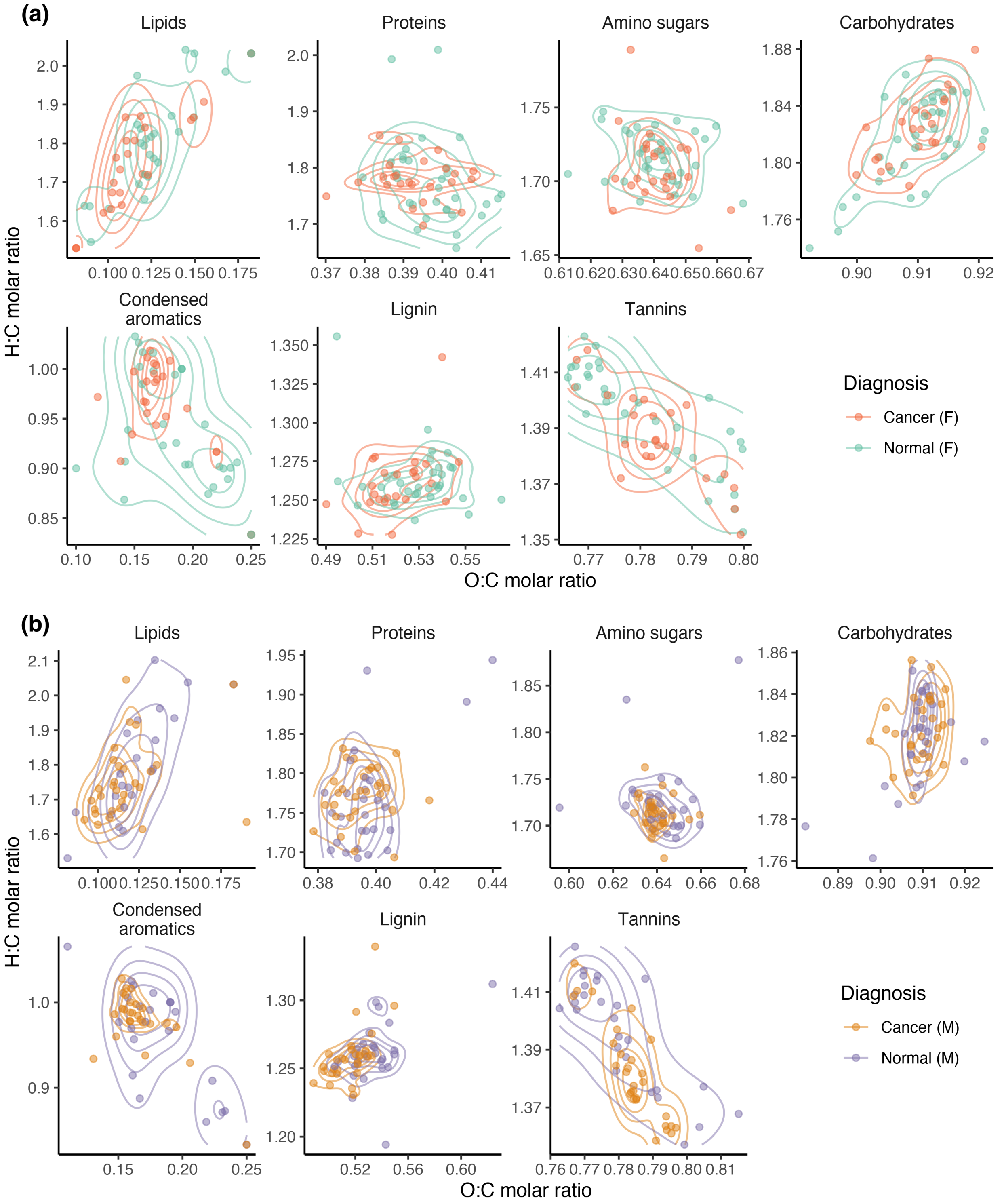


**Fig. S11.** Weighted mean vK coordinates within compound zones in colorectal cancer and normal subjects in (a) females (cancer n = 24, normal n = 33), and (b) males (cancer n = 29, normal n = 27).

Type 2 diabetes (T2D) case study data – female only (raw sequence data from Forslund et al., 2015)

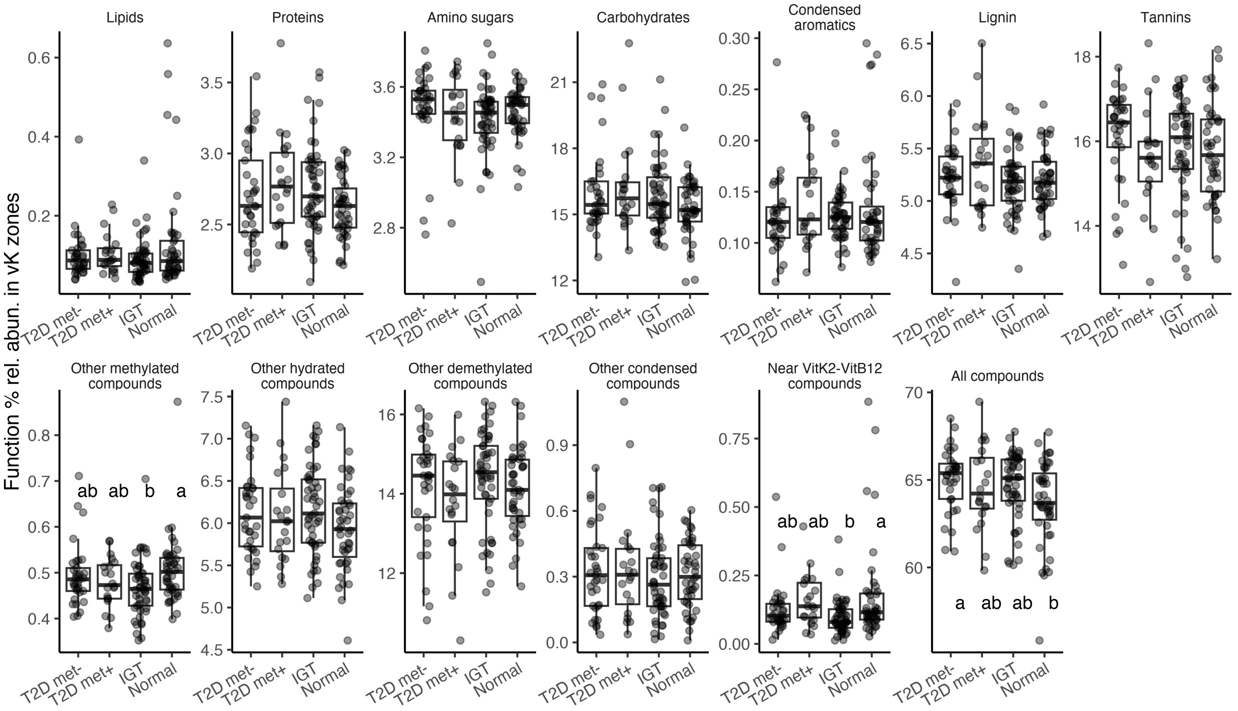


**Fig. S12.** Compound processing potential ($CPP_{class}$) in female only subjects diagnosed with type 2 diabetes with no Metformin (T2D met-, n = 33), type 2 diabetes with Metformin (T2D met+, n = 20), impaired glucose tolerance (IGT, n = 49), and normal (n = 43). Based on mapping of % functional relative abundances to compound-associated vK zones.

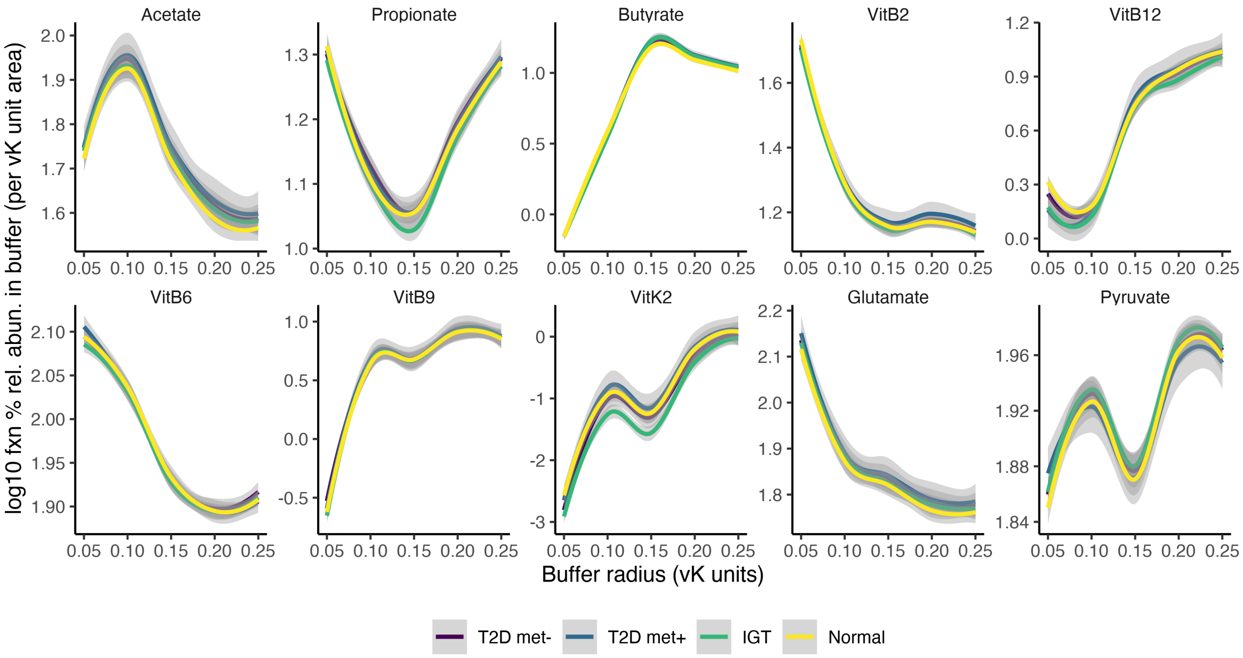


**Fig. S13.** Compound processing potential densities ($CPP_{density}$) in proximity to key health-associated biomolecules, based on log10 % function relative abundance mapped to radial buffers surrounding selected biomolecules, normalized (divided) by buffer area (T2D met- n = 33, T2D met+ n = 20, IGT n = 49, normal n = 43).

Type 2 diabetes (T2D) case study data – female only (continued)

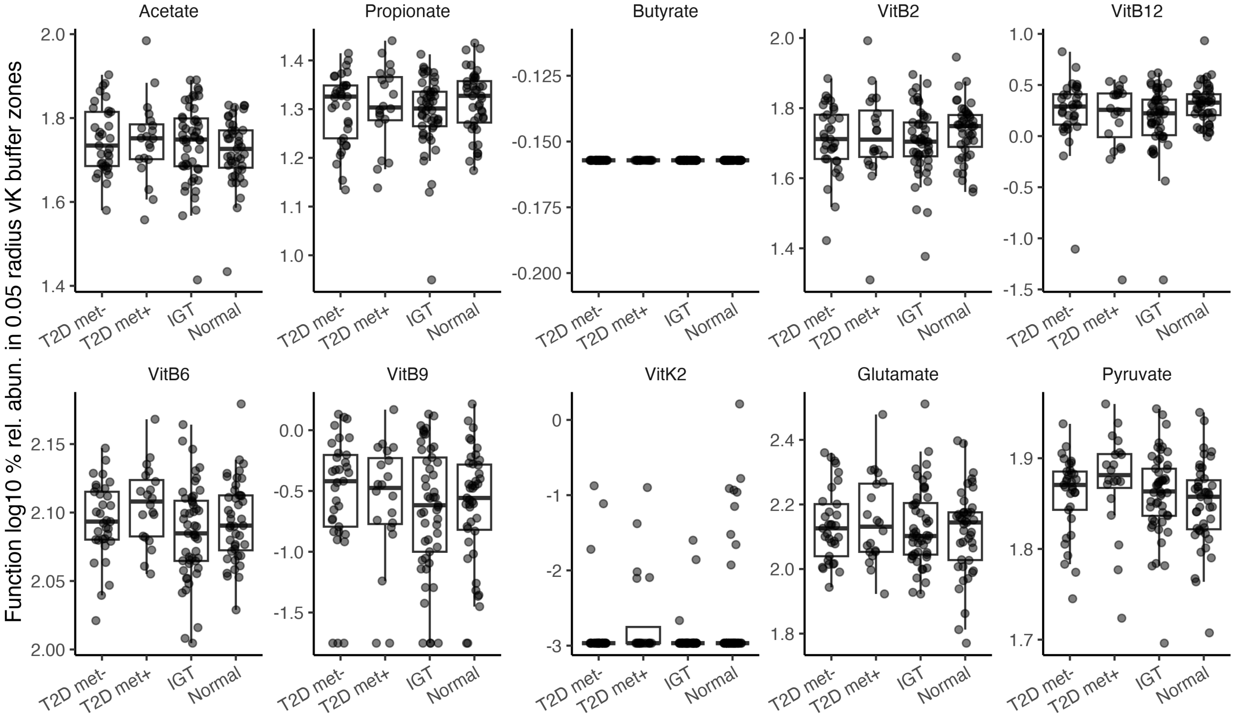


**Fig. S14.** Compound processing potential density ($CPP_{density}$) corresponding to key health-associated biomolecules, based on log10 % functional relative abundance mapped to the smallest radial buffer (0.05 vK units) normalized (divided) by buffer area (T2D met- n = 33, T2D met+ n = 20, IGT n = 49, normal n = 43).

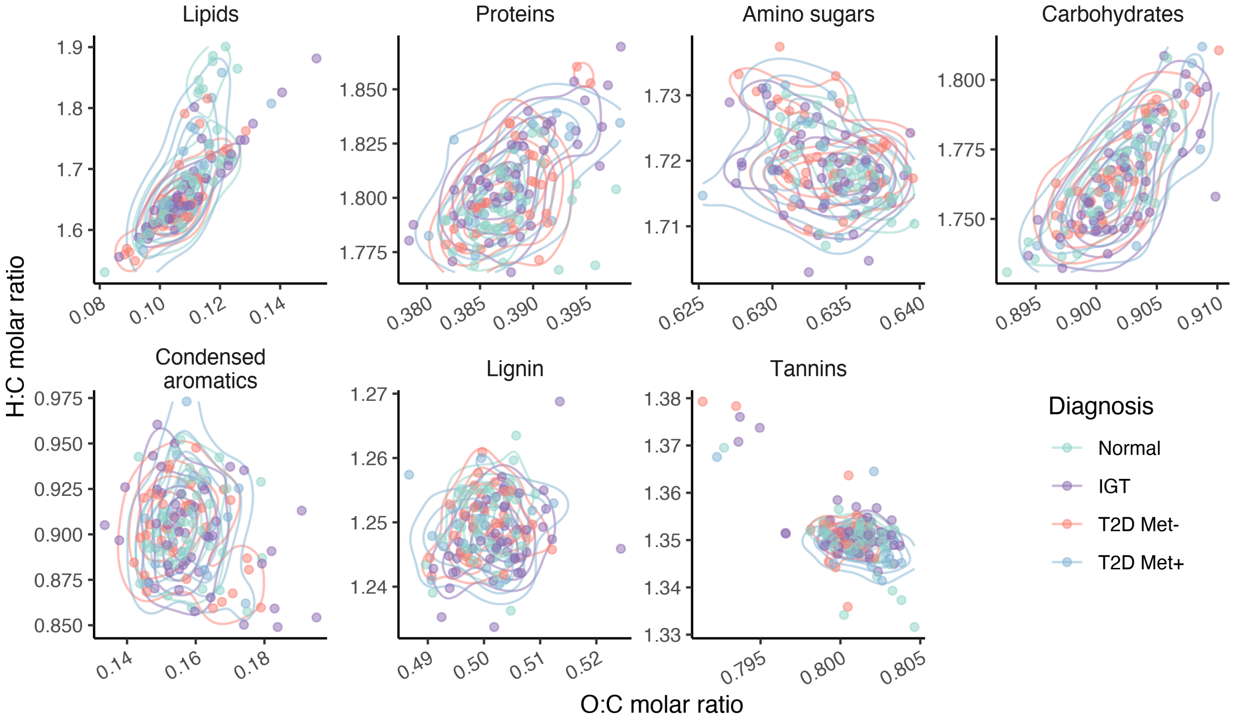


**Fig. S15.** Weighted mean vK coordinates within compound zones in female only subjects with diagnoses of T2D met- (n = 33), T2D met+ (n = 20), IGT (n = 49), normal n = 43.

Type 2 diabetes (T2D) case study data – female only (continued)

**(a)** Normal

(48 vertices, 96 edges)

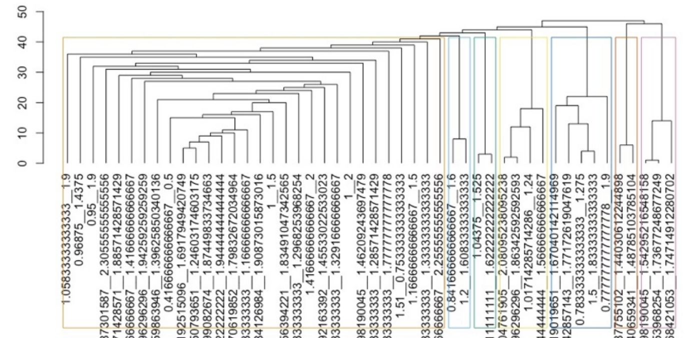

**(b)** IGT

(49 vertices, 67 edges)

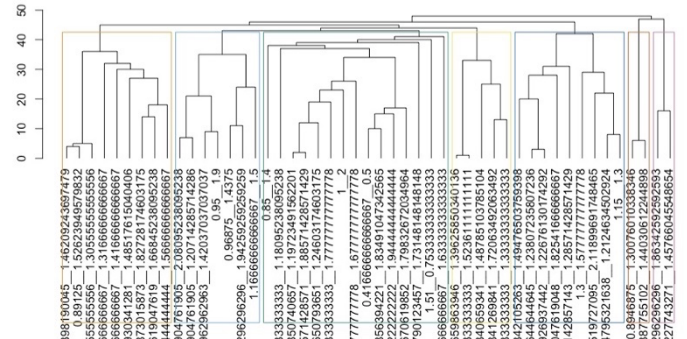

**(c)** T2D Met-

(173 vertices, 4028 edges)

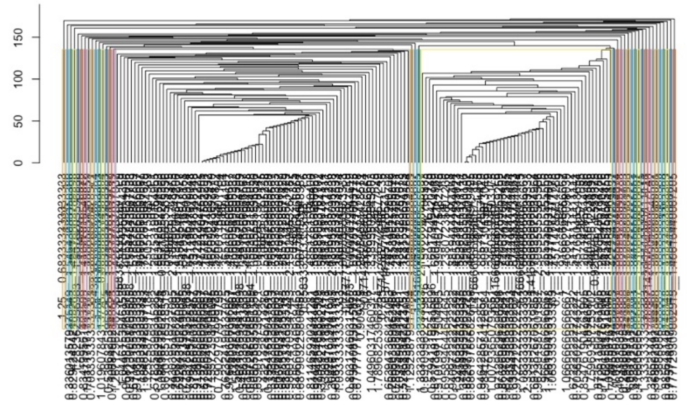

**(d)** T2D Met+

(103 vertices, 352 edges)

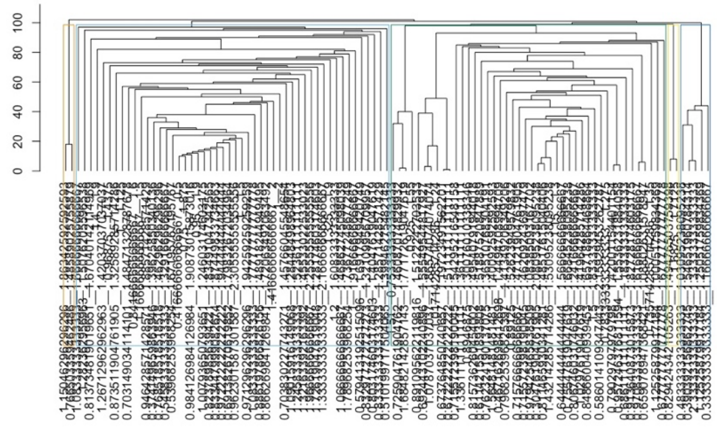

**Fig. S16.** Network structure dendogram plots based on correlations inferred among compound-associated vK coordinate profiles of 20 female subjects in each diagnosis group comprising (a) normal, (b) IGT, (c) T2D Met-, and (d) T2D Met+.

Type 2 diabetes (T2D) case study data – female only (continued)

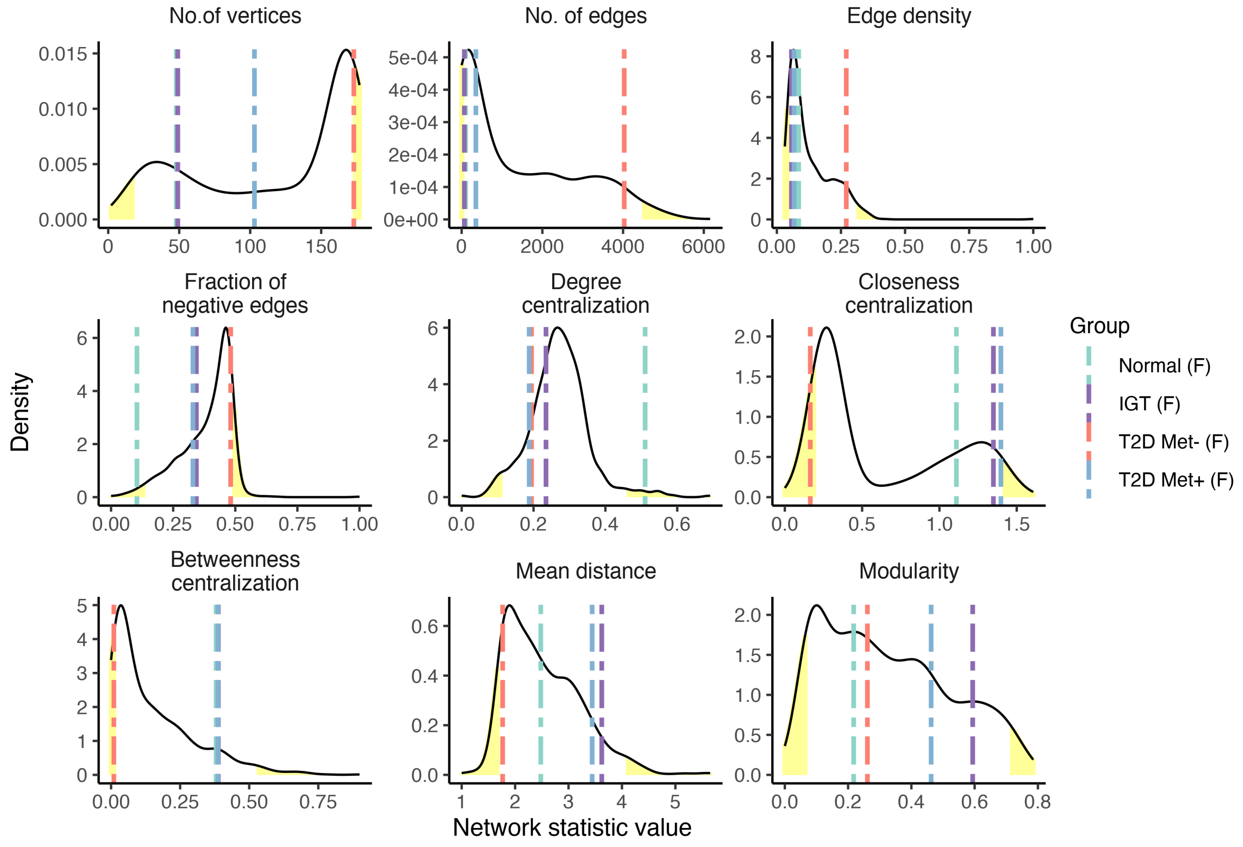


**Fig. S17.** Network characteristics from compound processing potential mapping in the type 2 diabetes (T2D) case study. Networks are based on correlations inferred among compound-associated van Krevelen coordinate profiles of 20 female subjects in each diagnosis group, including normal, impaired glucose tolerance (IGT), and T2D with and without Metformin treatment (Met+, Met-). Characteristics of diagnosis groups (vertical lines) are compared to randomized network probability density distributions generated from the same pipeline, but instead use bootstrap (B = 1000) resampling of 20 random subjects from the total pool of 80 subjects. Statistics are summarized in Table S17.

Problem behavior case study data (raw sequence data from Flannery et al., 2020)

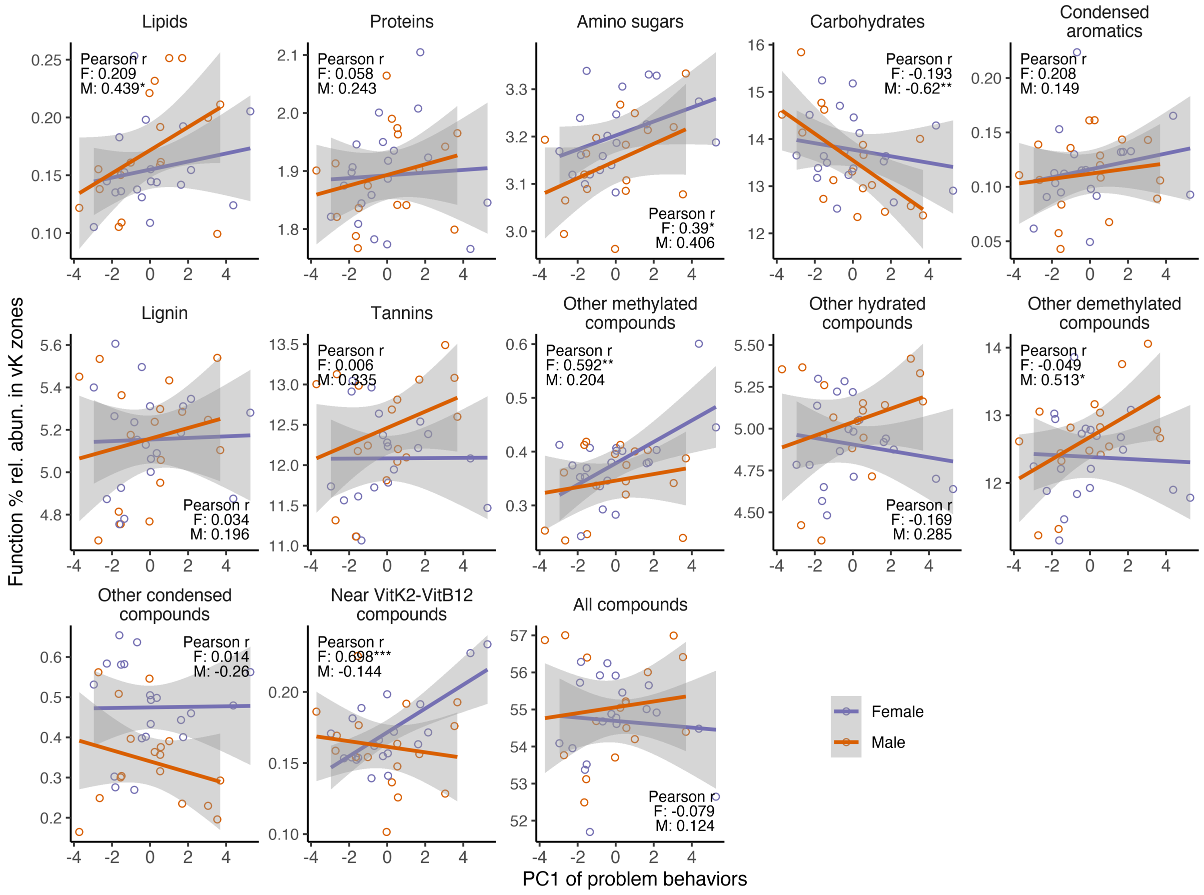


**Fig. S18.** Variation in compound processing potential ($CPP_{class}$) versus PC1 of problem behaviors in female (n = 20) and male (n = 17) children. Based on mapping of % functional relative abundances to compound-associated vK zones. The first principal component (PC1) for each sex is derived from behavioral scores outlined in Fig. S3.

Problem behavior case study data (continued)

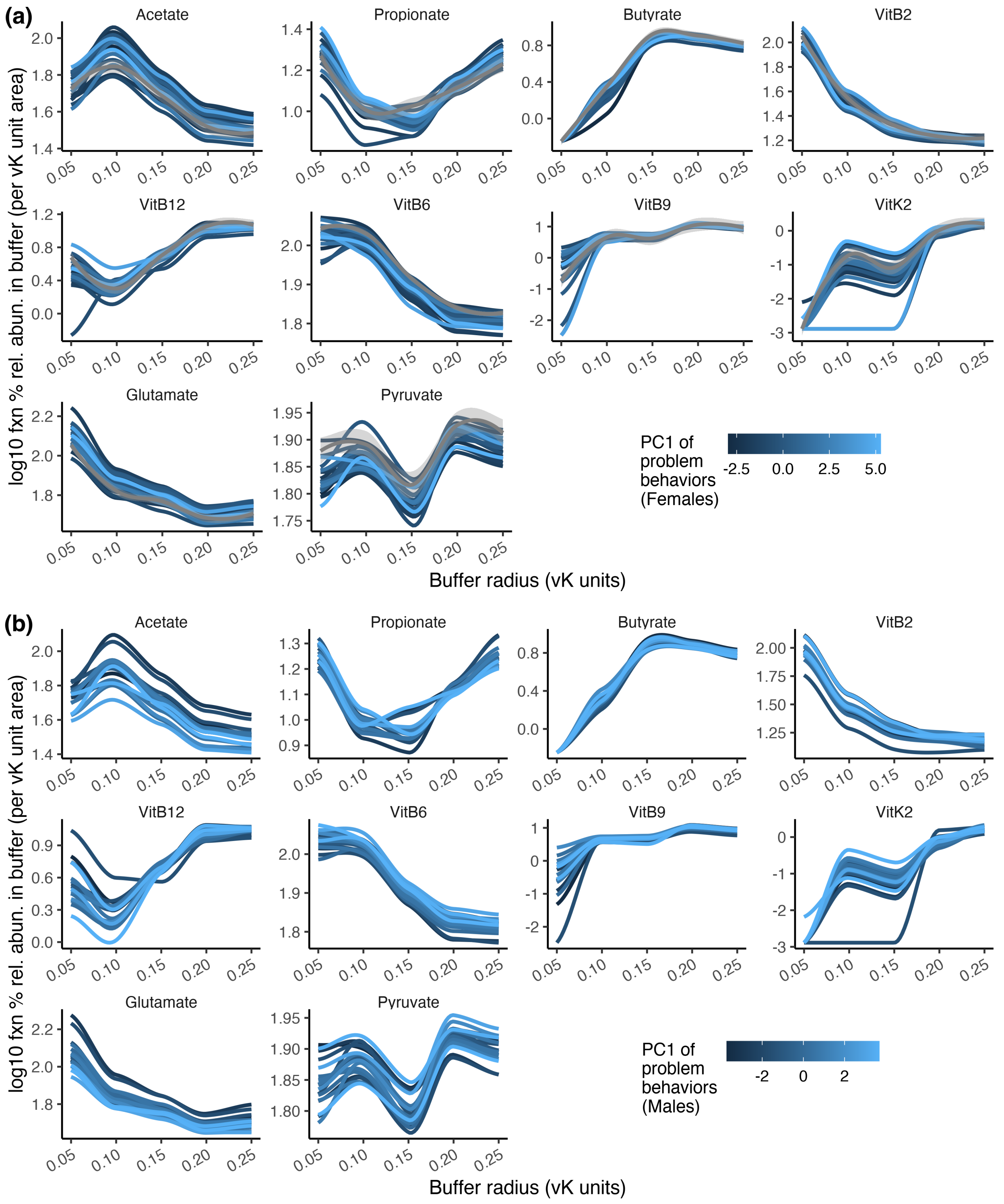


**Fig. S19.** Compound processing potential densities ($CPP_{density}$) in proximity to key health-associated biomolecules in (a) female (n = 20), and (b) male (n = 17) children. Based on log10 % function relative abundance mapped to radial buffers surrounding selected biomolecules, normalized (divided) by buffer area. Lines denote variation in values of the PC1 of problem behaviors.

Problem behavior case study data (continued)

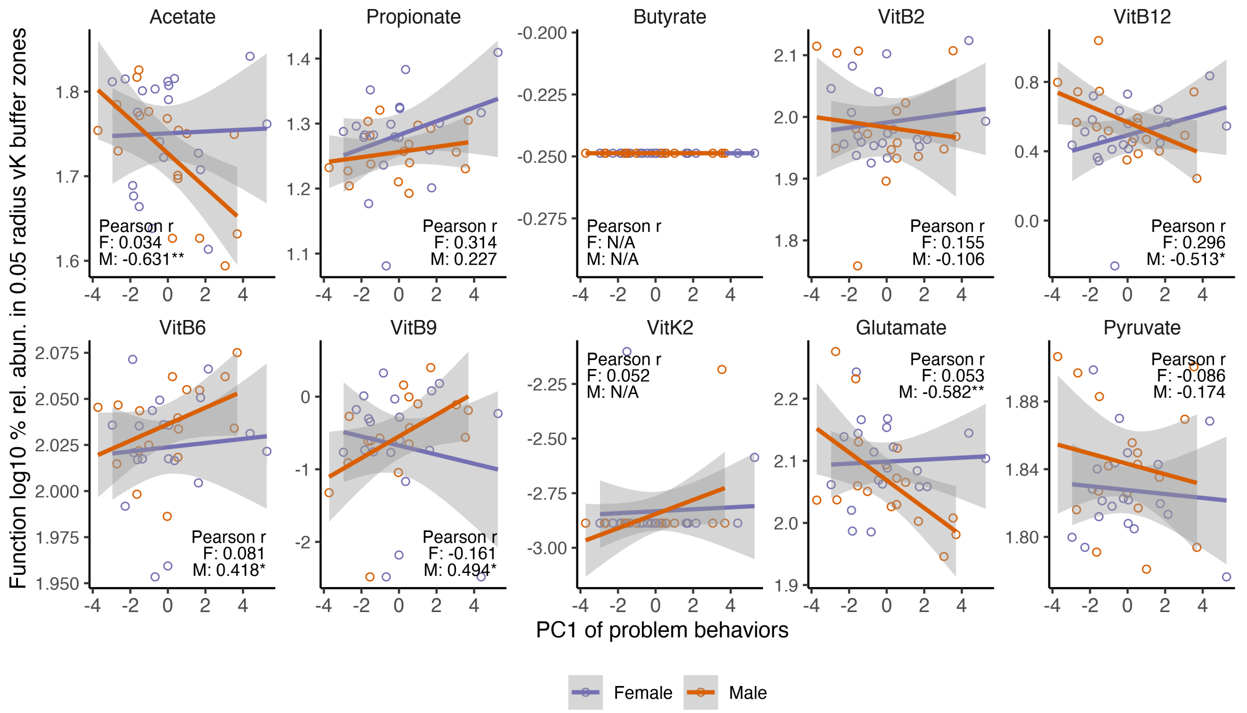


**Fig. S20.** Compound processing potential density ($CPP_{density}$) corresponding to key health-associated biomolecules in female (n = 20) and (b) male (n = 17) children. Based on log10 % functional relative abundance mapped to the smallest radial buffer (0.05 vK units) normalized (divided) by buffer area. The x-axis expresses variation in PC1 of problem behaviors, which are underpinned by measures outlined in Fig. S3.

Problem behavior case study data (continued)

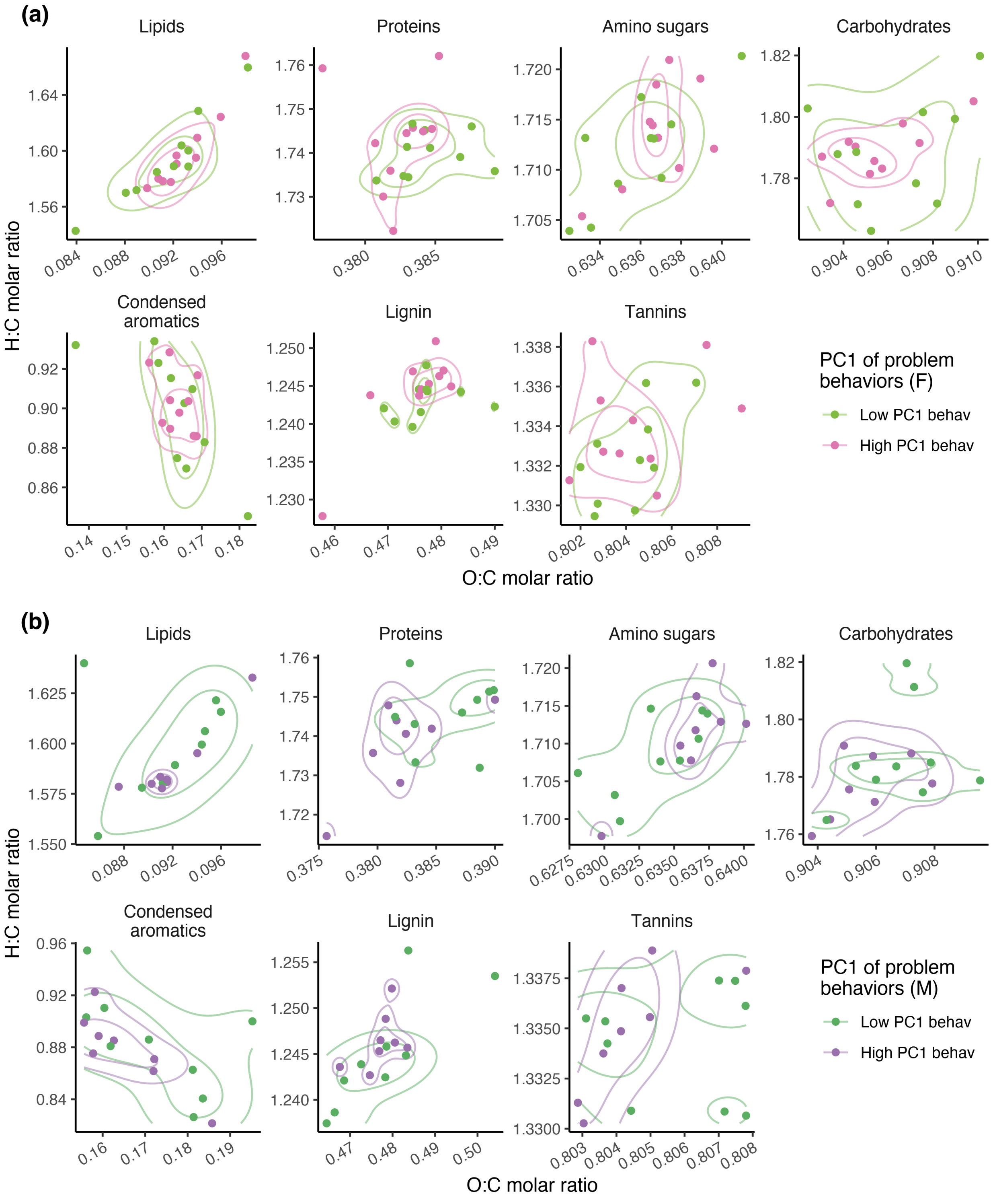


**Fig. S21.** Weighted mean vK coordinates within compound zones in (a) female (n = 20), and (b) male (n = 17) children. Data are expressed as groups of low or high PC1 of problem behaviors, where PC1 values are underpinned by measures outlined in Fig. S3.

Problem behavior case study data (continued)

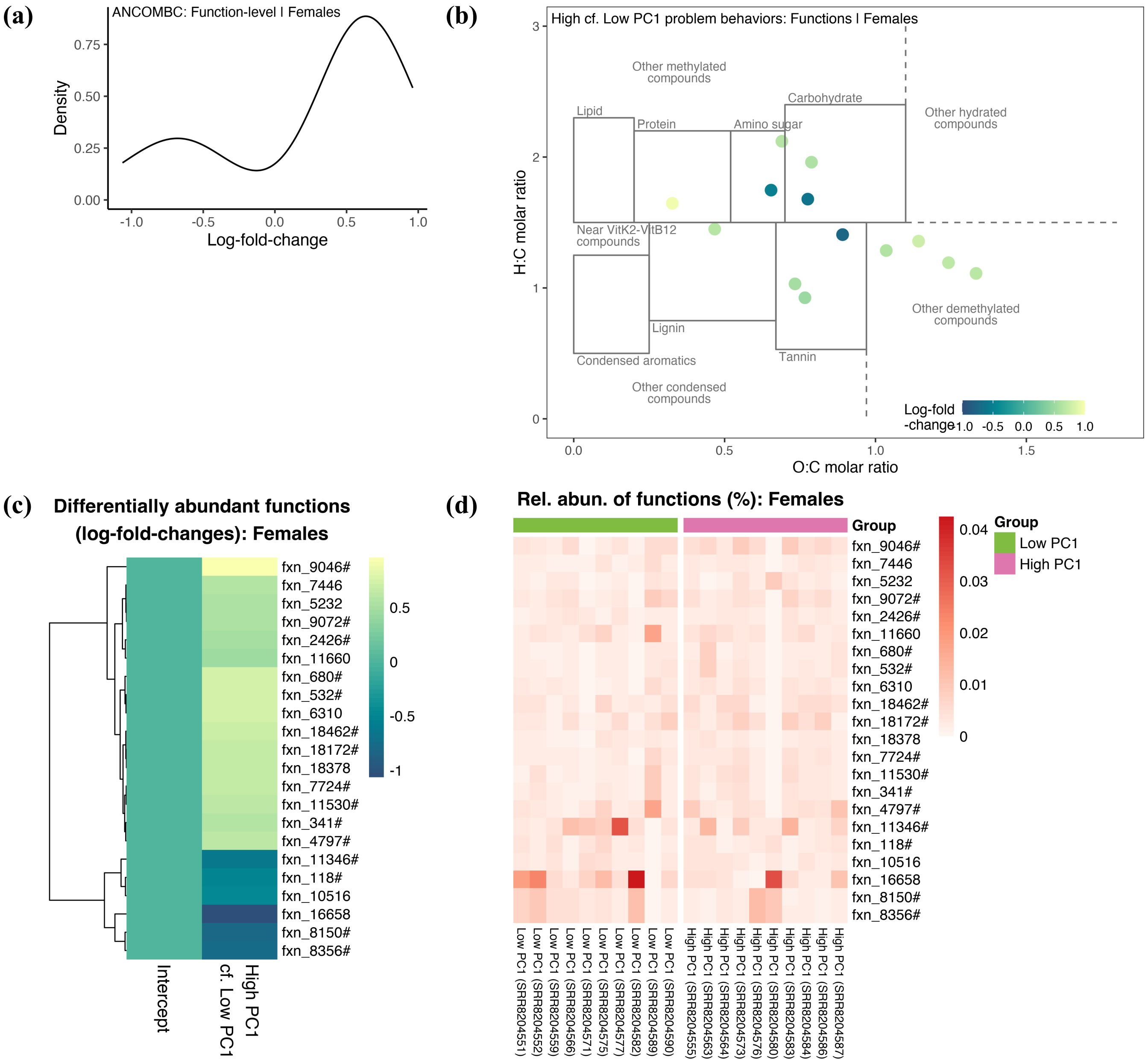


**Fig. S22.** Differentially abundant functions linked to problem behaviors in female children. Plots display (a) density distribution, (b) mapping to vK space, and (c) heatmap of log fold changes; and (d) relative abundances of functions, all comparing high PC1 (n = 10) to low PC1 (n = 10) of problem behaviors. Based on ANCOMBC analysis at function level which identified 22 differentially abundant functions, 15 of which (denoted by #) can be visualized in vK space.

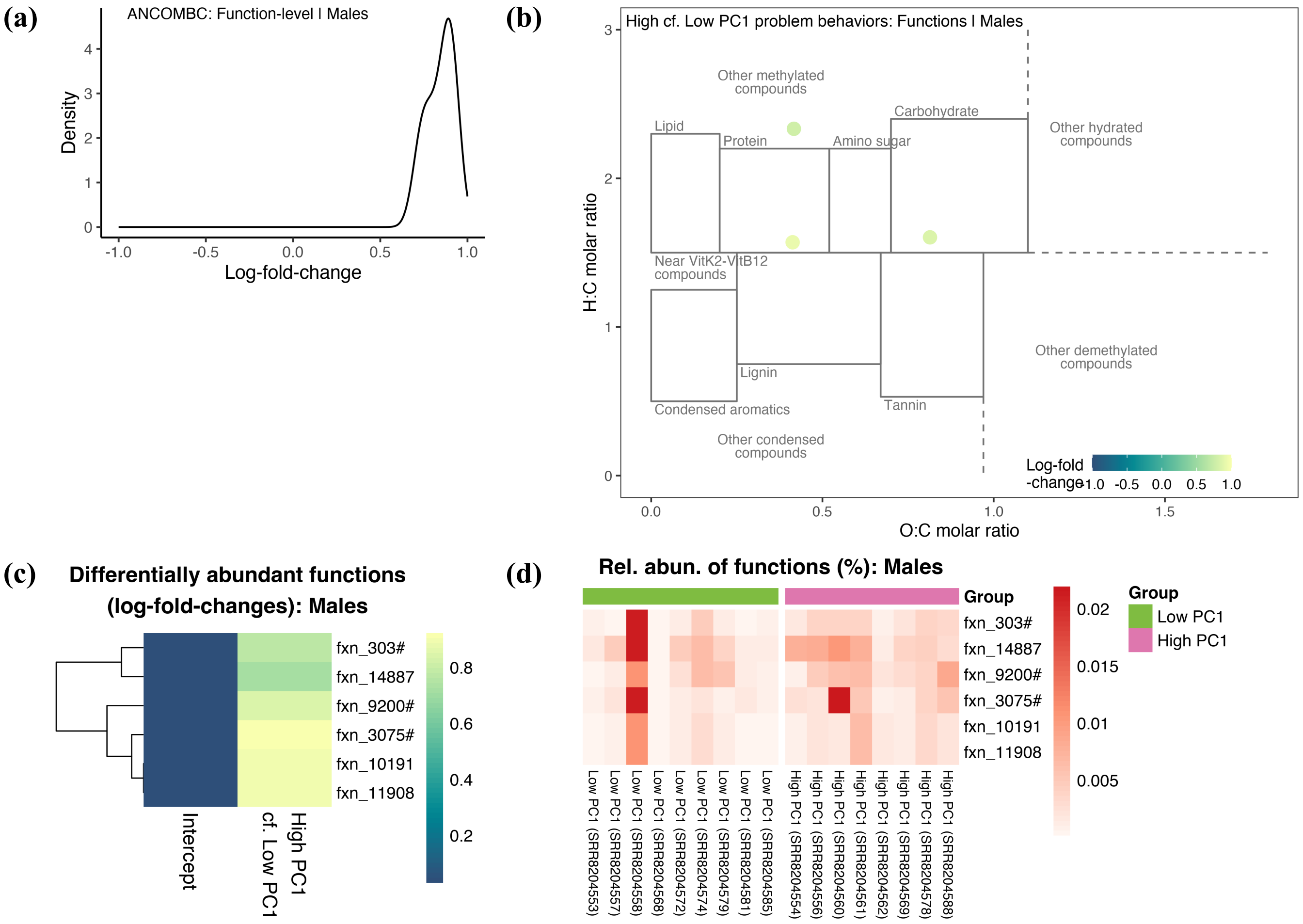


**Fig. S23.** Differentially abundant functions linked to problem behaviors in male children. Plots display (a) density distribution, (b) mapping to vK space, and (c) heatmap of log fold changes; and (d) relative abundances of functions, all comparing high PC1 (n = 8) to low PC1 (n = 9) of problem behaviors. Based on ANCOMBC analysis at function level which identified 6 differentially abundant functions, 3 of which (denoted by #) can be visualized in vK space.

Problem behavior case study data (continued)

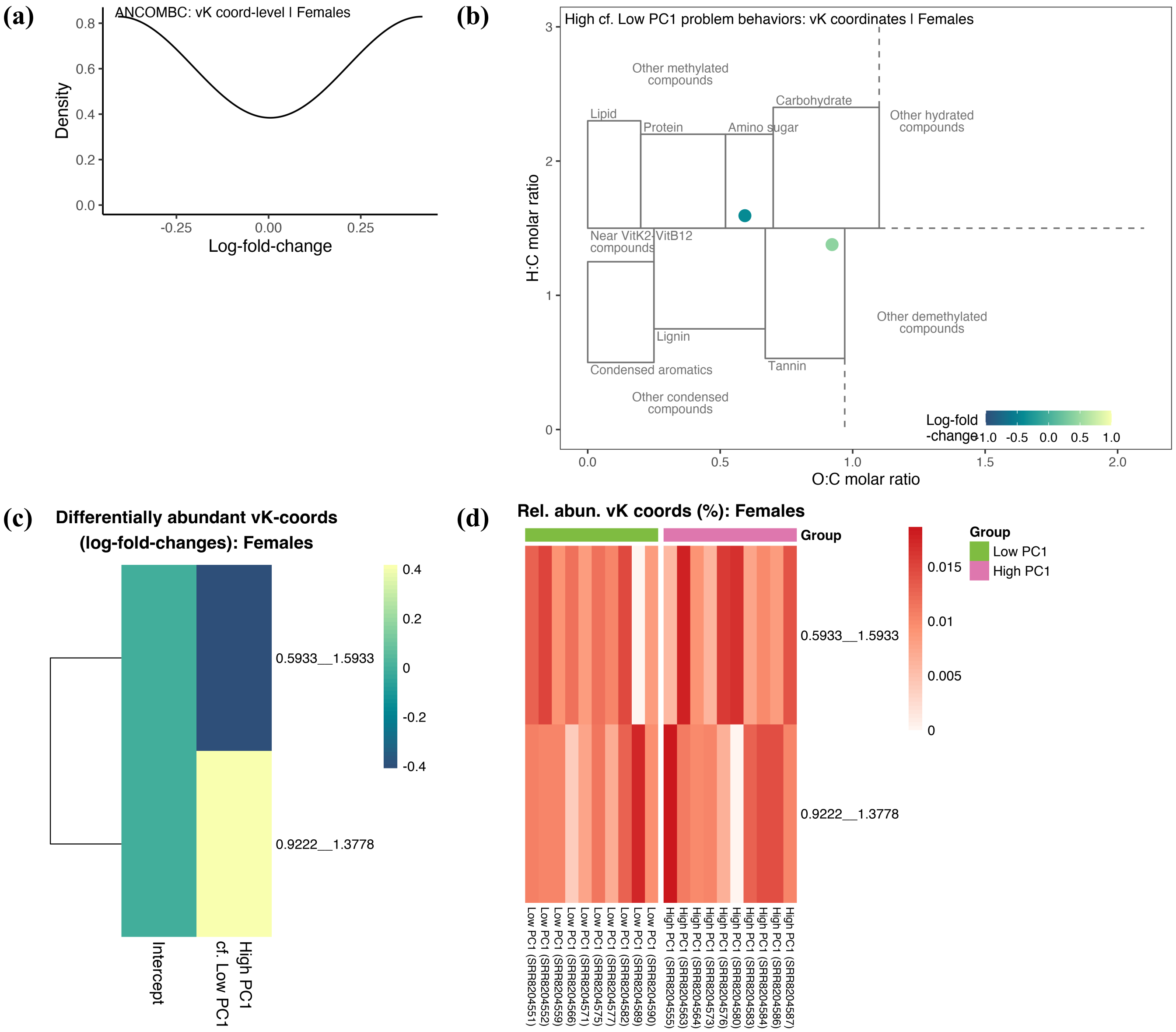


**Fig. S24.** Differentially abundant vK-coordinates linked to problem behaviors in female children. Plots display (a) density distribution, (b) mapping to vK space, and (c) heatmap of log fold changes; and (d) relative abundances of vK-coordinates, all comparing high PC1 (n = 10) to low PC1 (n = 10) of problem behaviors. Based on ANCOMBC analysis at vK-coordinate level.

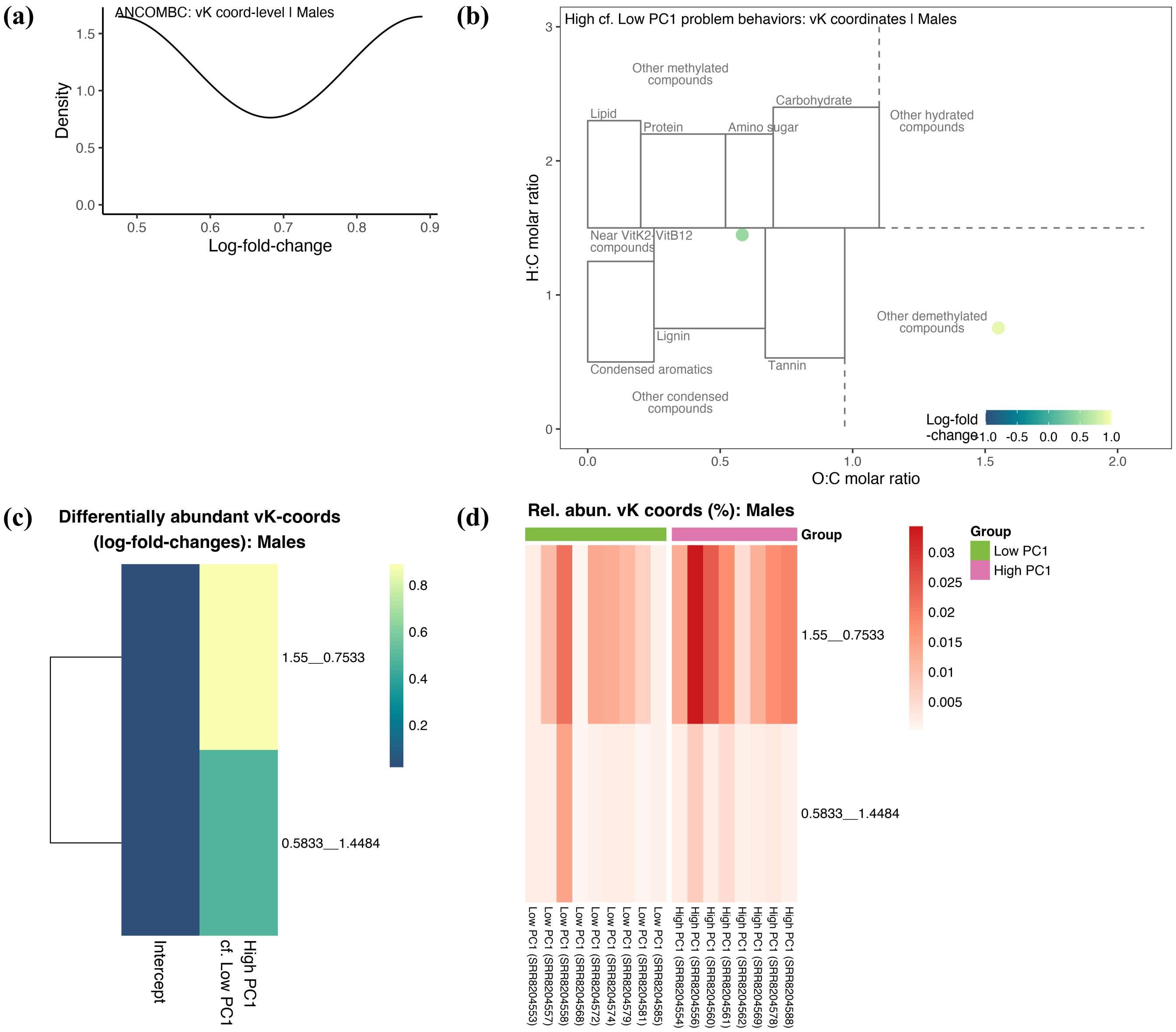


**Fig. S25.** Differentially abundant vK-coordinates linked to problem behaviors in male children. Plots display (a) density distribution, (b) mapping to vK space, and (c) heatmap of log fold changes; and (d) relative abundances of vK-coordinates, all comparing high PC1 (n = 8) to low PC1 (n = 9) of problem behaviors. Based on ANCOMBC analysis at vK-coordinate level.

People Cities and Nature (PCaN) urban ecosystem restoration case study data (raw sequence data from Barnes et al., 2020)

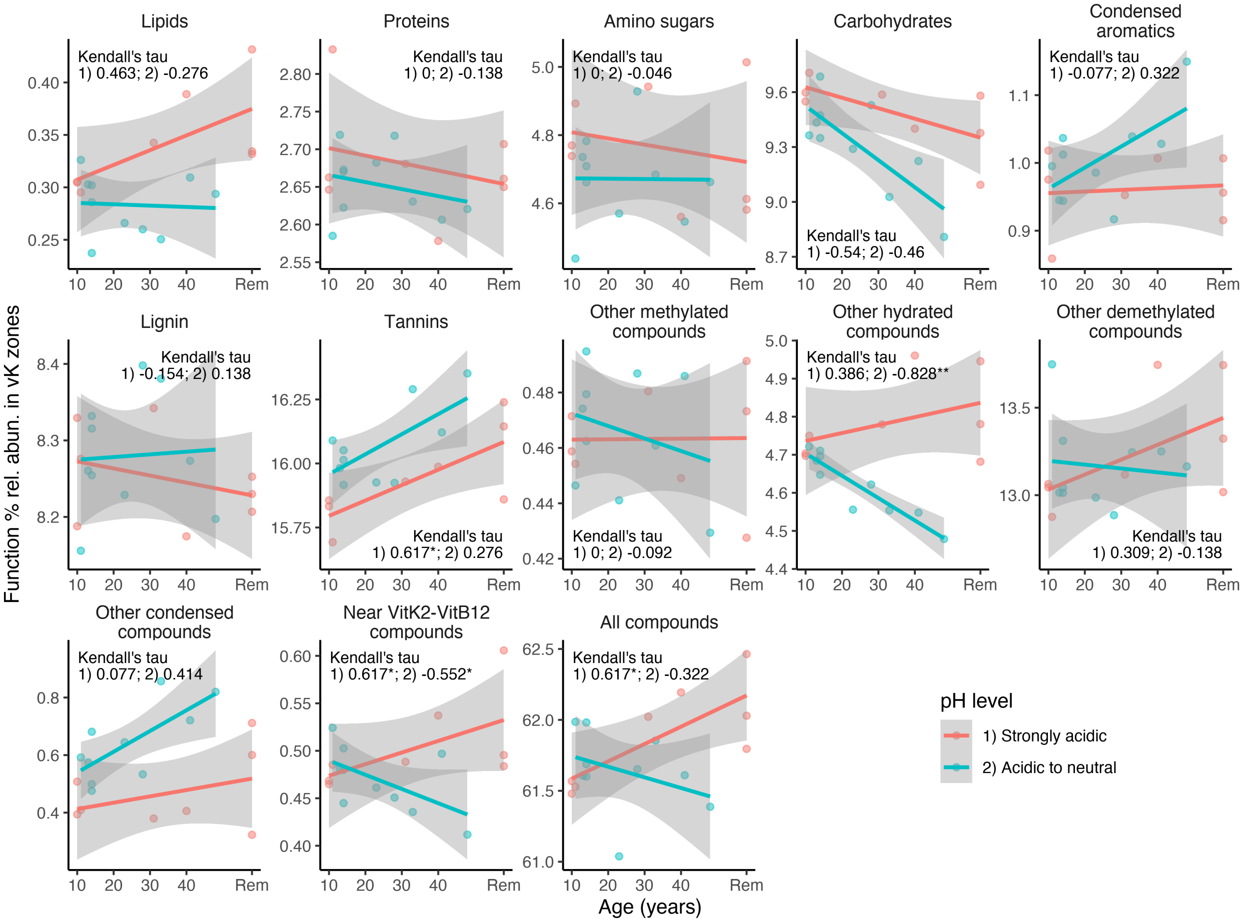


**Fig. S26.** Variation in compound processing potential ($CPP_{class}$) versus revegetation age (treated as ordinal data) at PCaN urban ecosystem restoration sites, across compound classes. Based on mapping of % functional relative abundances to compound-associated vK zones. Sites were analyzed separately for samples that were strongly acidic (pH < 4.5, n = 8) or acidic to neutral (4.5 < pH < 7, n = 10). Testing for significant trends was performed using Kendall's tau correlation values but visualized using linear trendlines.

PCaN urban ecosystem restoration case study data (continued)

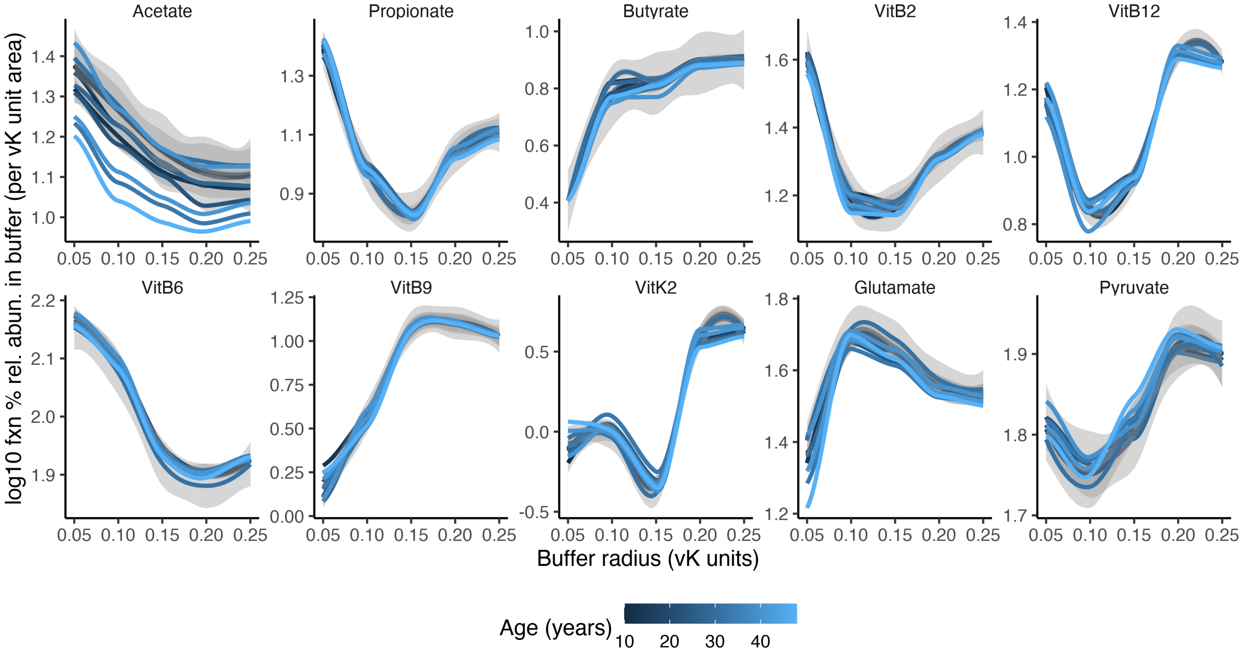


**Fig. S27.** Compound processing potential densities ($CPP_{density}$) in proximity to key health-associated biomolecules at PCaN urban ecosystem restoration sites, based on log10 % function relative abundance mapped to radial buffers surrounding selected biomolecules, normalized (divided) by buffer area. Lines denote variation in revegetation age (excluding remnants).

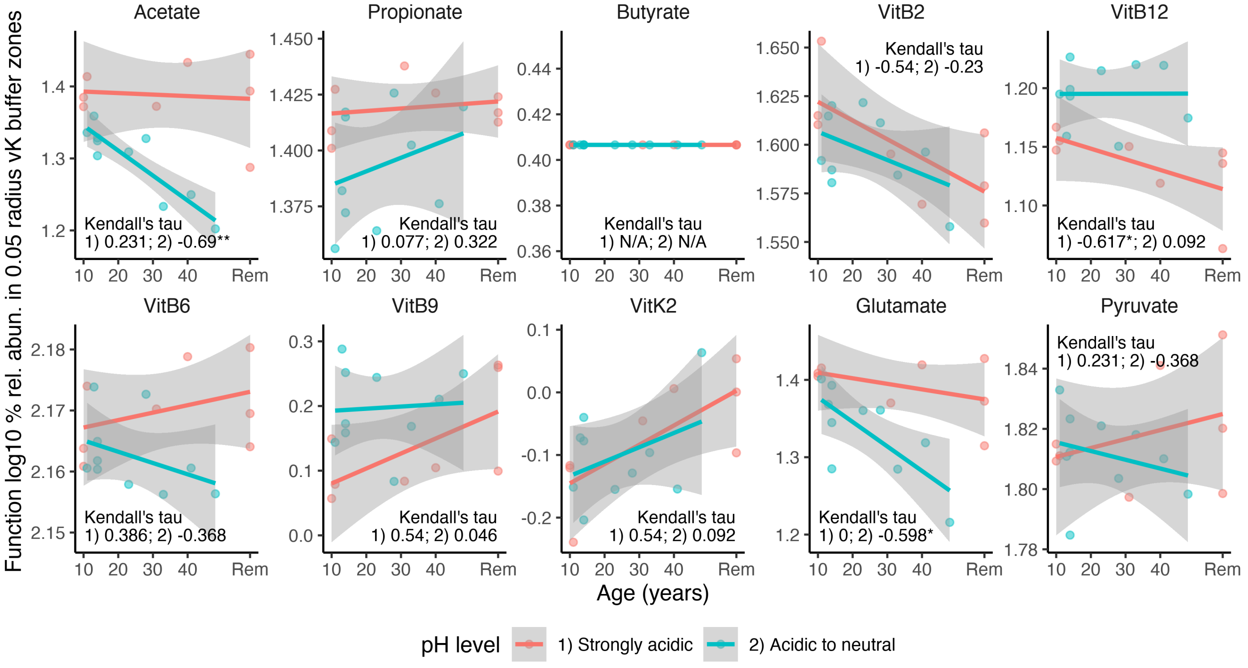


**Fig. S28.** Compound processing potential density ($CPP_{density}$) corresponding to key health-associated biomolecules at PCaN urban ecosystem restoration sites. Based on log10 % functional relative abundance mapped to the smallest radial buffer (0.05 vK units) normalized (divided) by buffer area. Revegetation age (ordinal data) is expressed on the x-axis. Sites were analyzed separately for samples that were strongly acidic (pH < 4.5, n = 8) or acidic to neutral (4.5 < pH < 7, n = 10). Testing for significant trends was performed using Kendall's tau correlation values but visualized using linear trendlines.

PCaN urban ecosystem restoration case study data (continued)

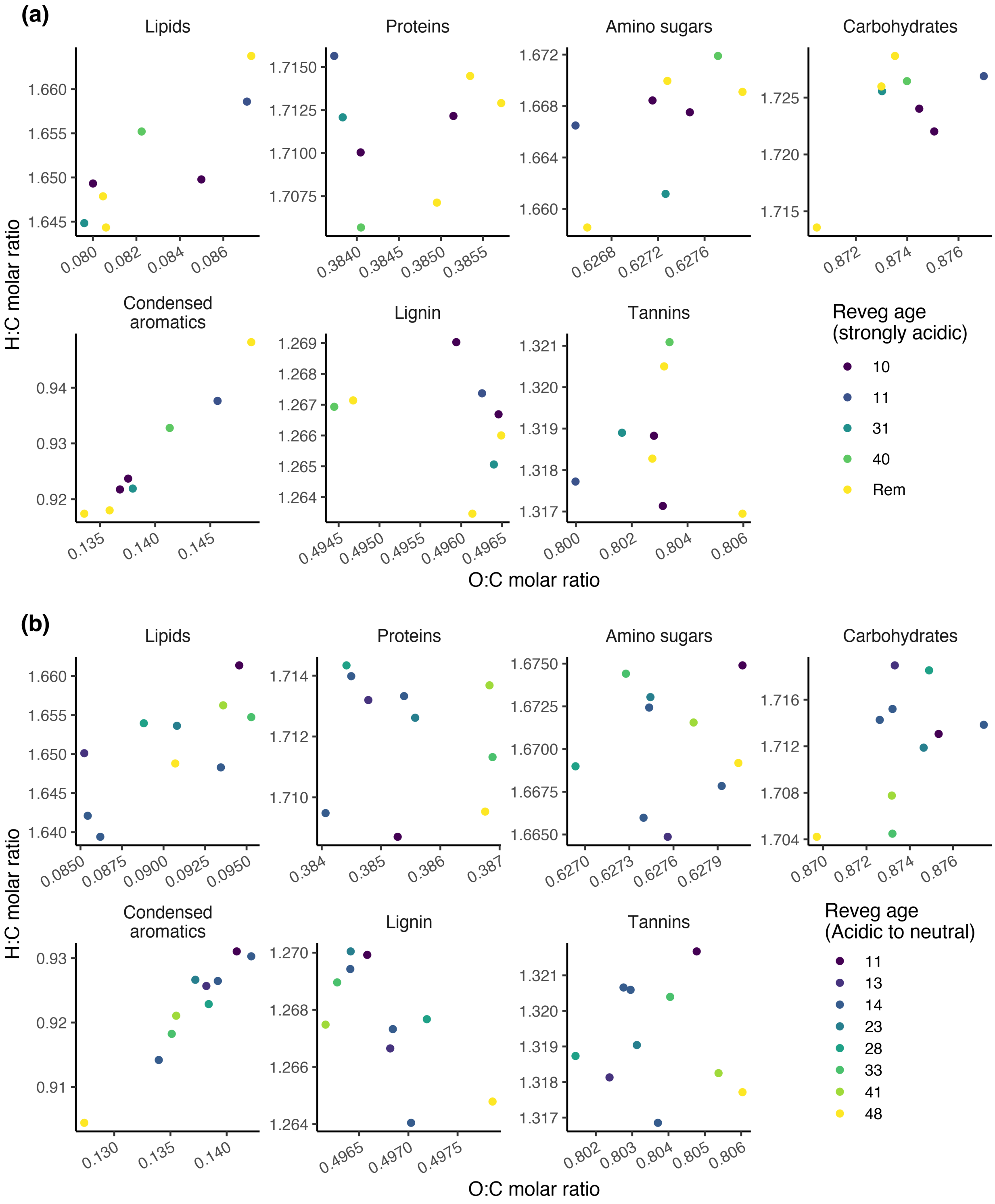


**Fig. S29.** Weighted mean vK coordinates within compound zones at PCaN urban ecosystem restoration sites, in (a) strongly acidic (pH < 4.5, n = 8) or (b) acidic to neutral (4.5 < pH < 7, n = 10) soils.

Post-mining forest restoration case study data (raw sequence data from Sun and Badgley, 2019)

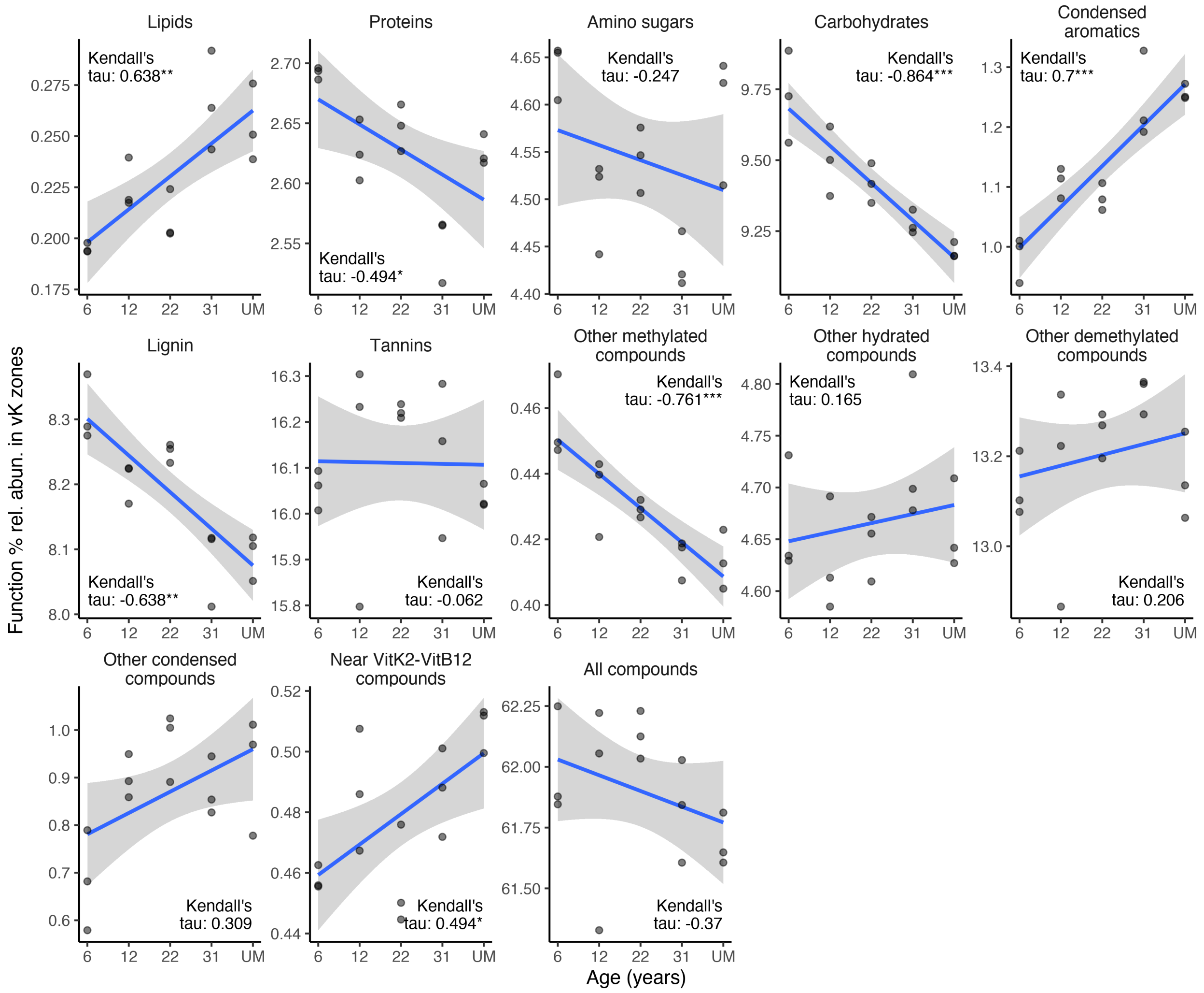


**Fig. S30.** Variation in compound processing potential ($CPP_{class}$) versus revegetation age (treated as ordinal data) at post-mining forest ecosystem restoration sites (n = groups of 3, across 5 revegetation age groups), across compound classes. Based on mapping of % functional relative abundances to compound-associated vK zones. Testing for significant trends was performed using Kendall's tau correlation values but visualized using linear trendlines.

Post-mining forest restoration case study data (continued)

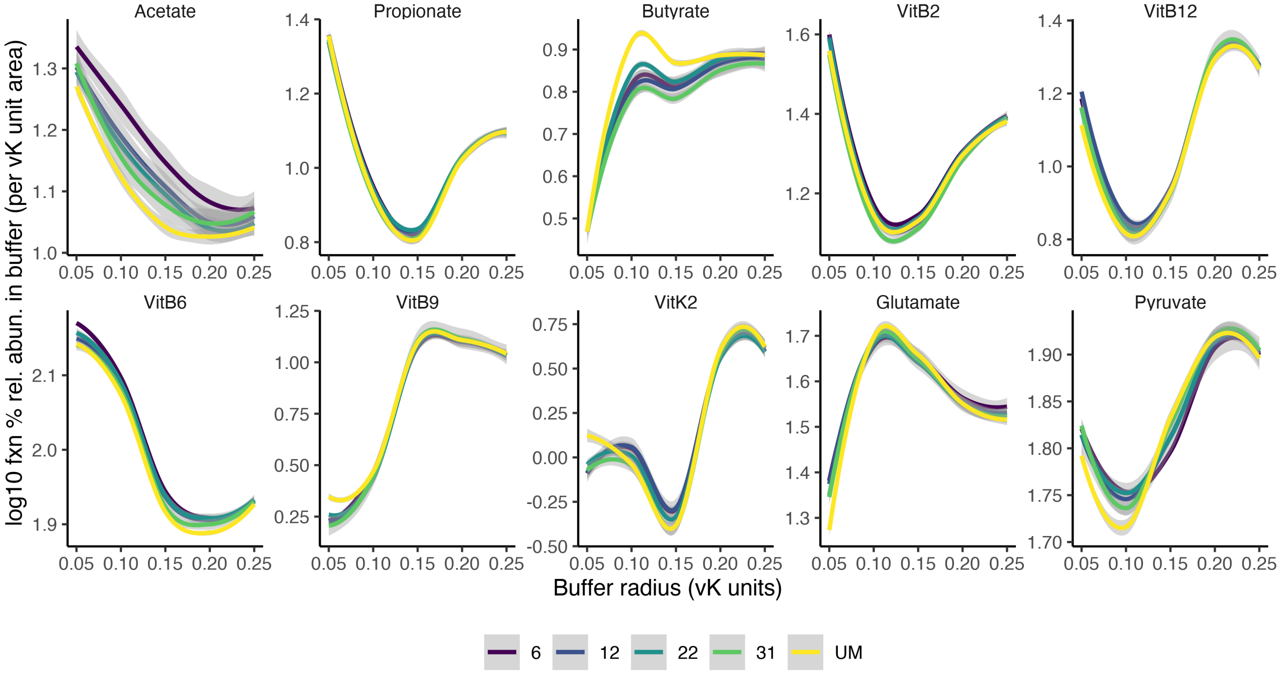


**Fig. S31.** Compound processing potential densities ($CPP_{density}$) in proximity to key health-associated biomolecules at post-mining forest ecosystem restoration sites (n = groups of 3, across 5 revegetation age groups). Based on log10 % function relative abundance mapped to radial buffers surrounding selected biomolecules, normalized (divided) by buffer area. Lines denote the age-based classes of forest ecosystem restoration sites.

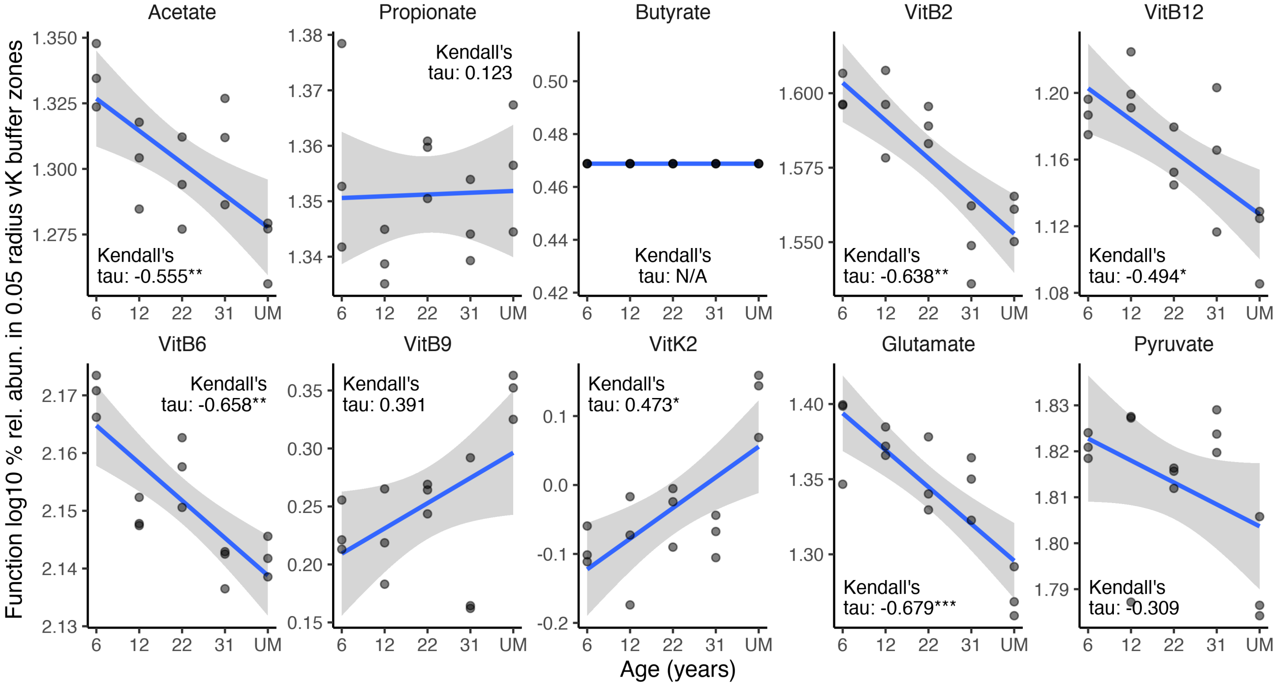


**Fig. S32.** Compound processing potential density ($CPP_{density}$) corresponding to key health-associated biomolecules at post-mining forest ecosystem restoration sites (n = groups of 3, across 5 revegetation age groups). Based on log10 % functional relative abundance mapped to the smallest radial buffer (0.05 vK units) normalized (divided) by buffer area. Age-based classes of forest ecosystem restoration are displayed on the x-axis.

Australian Microbiome Initiative (AMI) disturbed vs natural soil case study data (raw sequences from Bissett et al., 2016)

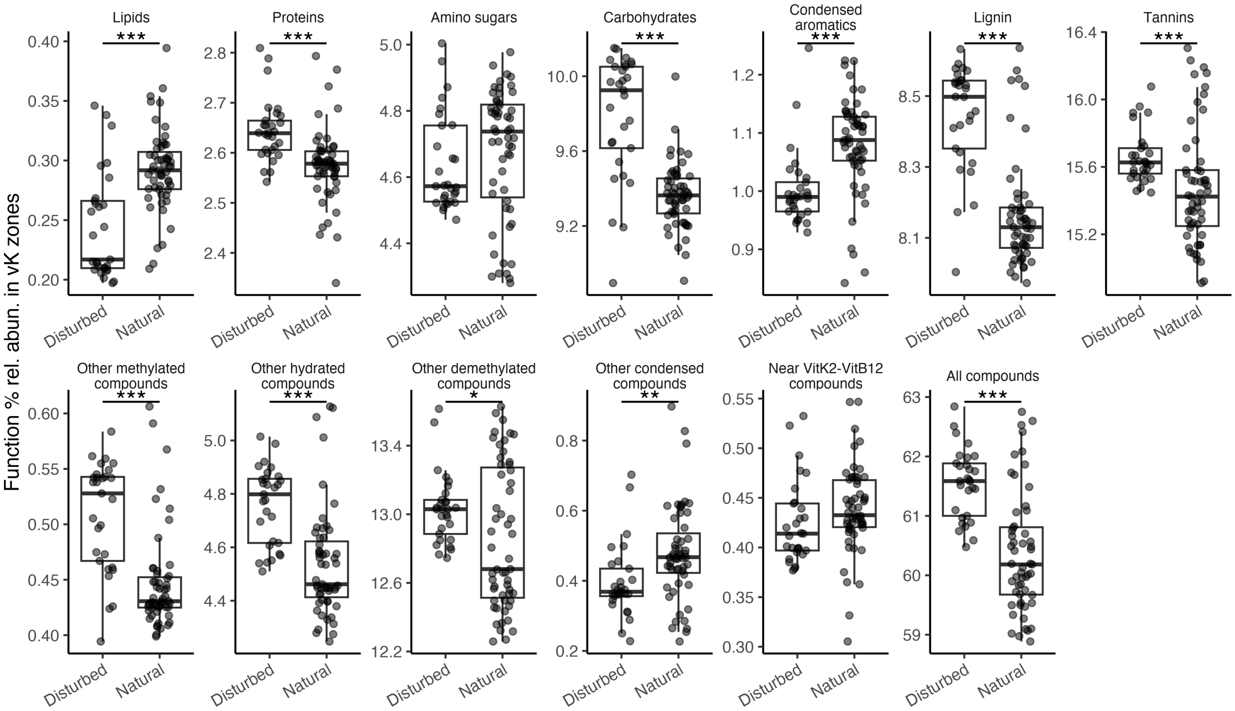


**Fig. S33.** Compound processing potential ($CPP_{class}$) in AMI disturbed (n = 29) versus natural (n = 55) soil samples, across compound classes, based on mapping of % functional relative abundances to compound-associated vK zones.

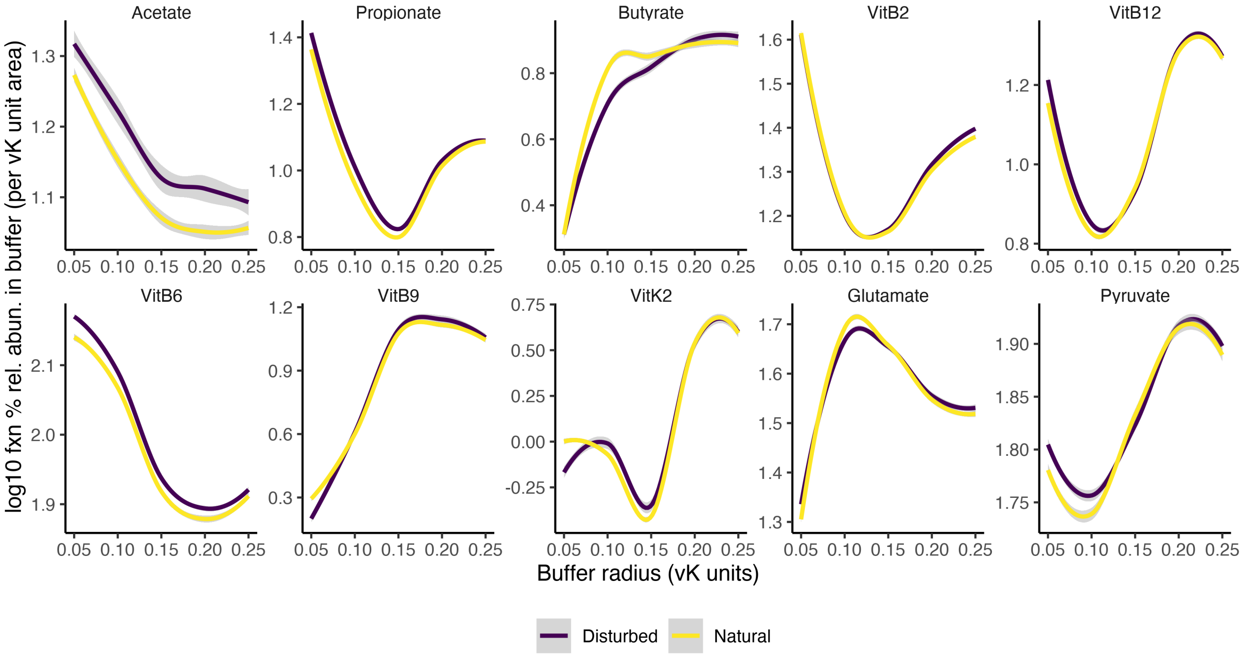


**Fig. S34.** Compound processing potential densities ($CPP_{density}$) in proximity to key health-associated biomolecules in AMI disturbed (n = 29) versus natural (n = 55) soil samples. Based on log10 % function relative abundance mapped to radial buffers surrounding selected biomolecules, normalized (divided) by buffer area.

AMI disturbed vs natural soil case study data (continued)

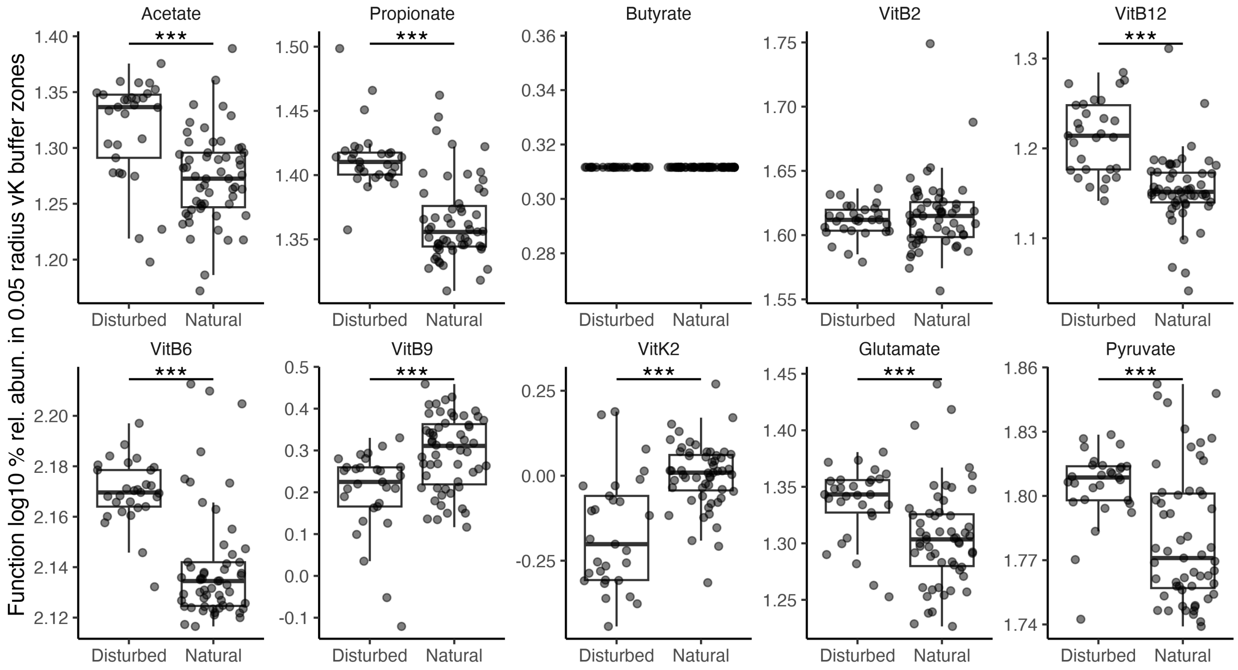


**Fig. S35.** Compound processing potential density ($CPP_{density}$) corresponding to key health-associated biomolecules in AMI disturbed (n = 29) versus natural (n = 55) soil samples, based on log10 % functional relative abundance mapped to the smallest radial buffer (0.05 vK units) normalized (divided) by buffer area, comparing disturbed versus natural samples.

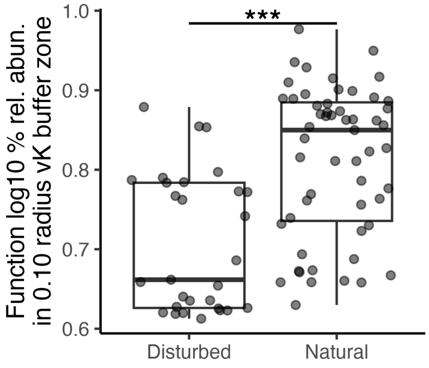


**Fig. S36.** Near butyrate compound processing potential density ($CPP_{density}$) in AMI disturbed (n = 29) versus natural (n = 55) soil samples, based on log10 % functional relative abundance mapped to the 0.1 vK unit radial buffer, normalized by buffer area.

**Table S1.** Van Krevelen (vK) compound zones adapted from (Wu et al., 2018).

| Compound class | O:C molar ratio (x-axis) | | H:C molar ratio (y-axis) | |
|---|---|---|---|---|
| | xmin | xmax | ymin | ymax |
| Lipids | ≥ 0 | ≤ 0.2 | ≥ 1.5 | ≤ 2.3 |
| Proteins | > 0.2 | ≤ 0.52 | ≥ 1.5 | ≤ 2.2 |
| Amino sugars | > 0.52 | ≤ 0.7 | ≥ 1.5 | ≤ 2.2 |
| Carbohydrates | > 0.7 | ≤ 1.1 | ≥ 1.5 | ≤ 2.4 |
| Condensed aromatics | ≥ 0 | ≤ 0.25 | ≥ 0.5 | ≤ 1.25 |
| Lignin | > 0.25 | ≤ 0.67 | ≥ 0.75 | < 1.5 |
| Tannins | > 0.67 | ≤ 0.97 | ≥ 0.53 | < 1.5 |
| Additional classes below were created to exhaustively cover the vK coordinate space – these were applied only for remaining vK-coordinates not already assigned to the classes above: | | | | |
| Other methylated | – | ≤ 1.1 | > 2.2 | – |
| Other hydrated | > 1.1 | – | ≥ 1.5 | – |
| Other demethylated | > 0.97 | – | – | < 1.5 |
| Other condensed | > 0 | ≤ 0.97 | > 0 | < 0.75 |
| Near Vit K2 – Vit B12 | ≥ 0 | ≤ 0.25 | > 1.25 | < 1.5 |

**Table S2.** Chemical formulae, O:C and H:C ratios of health-associated biomolecules and dietary or environmental substrates featured in Fig. 1 main article. Focus biomolecules in this study are highlighted in bold.

| Compound | Chemical formula | Atomic carbon | Atomic hydrogen | Atomic oxygen | O:C ratio | H:C ratio | Category (Fig. 1 main article)* | Source |
|---|---|---|---|---|---|---|---|---|
| **Acetate** | $C_2H4O_2$ | 6 | 12 | 6 | 1.0000 | 2.0000 | biomolecule | C00033[†] |
| **Propionate** | $C_3H_6O_2$ | 3 | 6 | 2 | 0.6667 | 2.0000 | biomolecule | C00163[†] |
| **Butyrate** | $C_4H_8O_2$ | 4 | 8 | 2 | 0.5000 | 2.0000 | biomolecule | C00246[†] |
| **Riboflavin (Vit B2)** | $C_{17}H_{20}N_4O_6$ | 17 | 20 | 6 | 0.3529 | 1.1765 | biomolecule | C00255[†] |
| **Cobalamin (Vit B12)** | $C_{62}H_{88}CoN_{13}O_{14}PR$ | 62 | 88 | 14 | 0.2258 | 1.4194 | biomolecule | C05776[†] |
| **Pyridoxal 5'-phosphate (Vit B6)** | $C_8H_{10}NO_6P$ | 8 | 10 | 6 | 0.7500 | 1.2500 | biomolecule | C00018[†] |
| **Folate (Vit B9)** | $C_{19}H_{19}N_7O_6$ | 19 | 19 | 6 | 0.3158 | 1.0000 | biomolecule | C00504[†] |
| **Menaquinone (Vit K2)** | $C_{16}H_{16}O_2(C_5H_8)n$ | 31 | 40 | 2 | 0.0645 | 1.2903 | biomolecule | C00828[†] |
| **Pyruvate** | $C_3H_4O_3$ | 3 | 4 | 3 | 1.0000 | 1.3333 | biomolecule | C00022[†] |
| **Glutamate** | $C_5H_9NO_4$ | 5 | 9 | 4 | 0.8000 | 1.8000 | biomolecule | C00025[†] |
| Trimethylamine | $C_3H_9N$ | 3 | 9 | 0 | 0.0000 | 3.0000 | biomolecule | C00565[†] |
| GABA | $C_4H_9NO_2$ | 4 | 9 | 2 | 0.5000 | 2.2500 | biomolecule | C00334[†] |
| Acetyl-CoA | $C_{23}H_{38}N_7O_{17}P_3S$ | 23 | 38 | 17 | 0.7391 | 1.6522 | biomolecule | C00024[†] |
| Noradrenaline | $C_8H_{11}NO_3$ | 8 | 11 | 3 | 0.3750 | 1.3750 | biomolecule | C00547[†] |
| Serotonin | $C_{10}H_{12}N_2O$ | 10 | 12 | 1 | 0.1000 | 1.2000 | biomolecule | C00780[†] |
| p-cresol | $C_7H_8O$ | 7 | 8 | 1 | 0.1429 | 1.1429 | biomolecule | C01468[†] |
| Dopamine | $C_8H_{11}NO_2$ | 8 | 11 | 2 | 0.2500 | 1.3750 | biomolecule | C03758[†] |
| Indole | $C_8H_7N$ | 8 | 7 | 0 | 0.0000 | 0.8750 | biomolecule | C00463[†] |
| Sucrose; Maltose; Lactose | $C_{12}H_{22}O_{11}$ | 12 | 22 | 11 | 0.9167 | 1.8333 | substrate | C00089; C00208; C00243[†] |
| Glucose | $C_6H_{12}O_6$ | 6 | 12 | 6 | 1.0000 | 2.0000 | substrate | C00031[†] |
| Glycogen | $C_{24}H_{42}O_{21}$ | 24 | 42 | 21 | 0.8750 | 1.7500 | substrate | C00182[†] |
| Cellulose; Amylose | $(C_6H_{10}O_5)n$ | 6 | 10 | 5 | 0.8333 | 1.6667 | substrate | C00760; C00718[†] |
| Chitin | $(C_8H_{13}NO_5)n$ | 8 | 13 | 5 | 0.6250 | 1.6250 | substrate | C00461[†] |
| Stearic acid | $C_{18}H_{36}O_2$ | 18 | 36 | 2 | 0.1111 | 2.0000 | substrate | C01530[†] |
| Oleic acid | $C_{18}H_{34}O_2$ | 18 | 34 | 2 | 0.1111 | 1.8889 | substrate | C00712[†] |
| Linoleic acid | $C_{18}H_{32}O_2$ | 18 | 32 | 2 | 0.1111 | 1.7778 | substrate | C01595[†] |
| Ethanol | $C_2H_6O$ | 2 | 6 | 1 | 0.5000 | 3.0000 | substrate | C00469[†] |
| Nicotine | $C_{10}H_{14}N_2$ | 10 | 14 | 0 | 0.0000 | 1.4000 | substrate | C16150[†] |
| a-pinene; Camphene | $C_{10}H_{16}$ | 10 | 16 | 0 | 0.0000 | 1.6000 | substrate | C09880; C06076[†] |
| Benzaldehyde | $C_7H_6O$ | 7 | 6 | 1 | 0.1429 | 0.8571 | substrate | C00261[†] |
| Wheat straw | | 4.4583 | 6.1 | 2.51875 | 0.5650 | 1.3682 | substrate | (De et al., 2018) |

| **Compound** | **Chemical formula** | **Atomic carbon** | **Atomic hydrogen** | **Atomic oxygen** | **O:C ratio** | **H:C ratio** | **Category (Fig. 1 main article)*** | **Source** |
|---|---|---|---|---|---|---|---|---|
| Corncob | | 4.0833 | 5.1 | 2.86875 | 0.7026 | 1.2490 | substrate | (De et al., 2018) |
| Softwood (Pine chips) | | 4.3417 | 6.1 | 2.59375 | 0.5974 | 1.4050 | substrate | (De et al., 2018) |
| Switchgrass | | 4.0167 | 6 | 2.8375 | 0.7064 | 1.4938 | substrate | (De et al., 2018) |
| Lignin | | 4.425 | 5.9 | 2.4875 | 0.5621 | 1.3333 | substrate | (De et al., 2018) |
| Cellulose | | 3.6333 | 6.5 | 3.10625 | 0.8549 | 1.7890 | substrate | (De et al., 2018) |
| Wood chips | | 4.8417 | 6.2 | 2.21875 | 0.4583 | 1.2806 | substrate | (De et al., 2018) |
| Wood | | 4.2917 | 5.3 | 2.6875 | 0.6262 | 1.2350 | substrate | (De et al., 2018) |
| Hardwood (Casuarina equisetifolia) | | 4.575 | 6.7 | 2.23125 | 0.4877 | 1.4645 | substrate | (De et al., 2018) |
| Switchgrass | | 3.6 | 5.7 | 3.1375 | 0.8715 | 1.5833 | substrate | (De et al., 2018) |
| Corncob | | 3.9 | 6 | 2.91875 | 0.7484 | 1.5385 | substrate | (De et al., 2018) |
| Sewage sludge | | 4.5667 | 7.5 | 1.65 | 0.3613 | 1.6423 | substrate | (De et al., 2018) |
| Sewage sludge | | 4.3583 | 8 | 2.01875 | 0.4632 | 1.8356 | substrate | (De et al., 2018) |
| Hemicellulose | $C_5H_{10}O_4$ | 6 | 10 | 4 | 0.6667 | 1.6667 | substrate | (Anca-Couce and Obernberger, 2016) |
| Hardwood (Beech chips) | | 4.0367 | 6.03 | 2.77875 | 0.6884 | 1.4938 | substrate | (Anca-Couce and Obernberger, 2016) |
| Softwood (Spruce chips) | | 4.1783 | 6.16 | 2.70125 | 0.6465 | 1.4743 | substrate | (Anca-Couce and Obernberger, 2016) |
| Humic acid (soil) | | | | | 0.5000 | 1.0400 | substrate | (Rice and MacCarthy, 1991; Sposito, 2008) |
| Fulvic acid (soil) | | | | | 0.7800 | 1.3500 | substrate | (Rice and MacCarthy, 1991; Sposito, 2008) |
| Meat protein | | 4.3333 | 6 | 1.1875 | 0.2740 | 1.3850 | substrate | (Feiner, 2006) |

*Health-associated biomolecule, or dietary or environmental substrate. †Compound number from KEGG Compound Database: https://www.genome.jp/kegg/compound/

**Table S3.** Summary results for $CPP_{class}$ data, i.e., total function % relative abundance mapped to compound-associated vK zones. Corresponding plots are noted. Human gut results indicate increased (+) or decreased (–) compound processing potential ($CPP_{class}$) observed in the disease state compared to the normal healthy state. Soil results indicate increased (+) or decreased (–) $CPP_{class}$ values with ecosystem maturity.

<table>
<tr><th></th><th colspan="7">Human gut</th><th colspan="3">Soils</th></tr>
<tr><td></td><td colspan="2">ACVD cf. Normal (Jie et al., 2017) (Fig. S4) [†]</td><td colspan="2">Colorectal cancer cf. Normal (Zeller et al., 2014) (Fig. S8) [†]</td><td>T2D Met- / T2D Met+/ IGT/ Normal (Forslund et al., 2015) (Fig. S12) [‡]</td><td colspan="2">Trend with PC1 of problem behaviors (Flannery et al., 2020) (Fig. S18) [§, #]</td><td>PCaN: Trend with Reveg age (Barnes et al., 2020) (Fig. S26) [¶, #]</td><td>Post-mining: Trend with Reveg age (Sun and Badgley, 2019) (Fig. S30) [¶, #]</td><td>AMI: Natural cf. disturbed (Bissett et al., 2016) (Fig. S33) [†]</td></tr>
<tr><td>Sex / pH class</td><td>F</td><td>M</td><td>F</td><td>M</td><td>F</td><td>F</td><td>M</td><td>1) | 2) [‖]</td><td></td><td></td></tr>
<tr><td>N</td><td>ACVD 53<br>Norm 101</td><td>ACVD 157<br>Norm 69</td><td>Cancer 24<br>Normal 33</td><td>Cancer 29<br>Normal 27</td><td>T2D Met- 33,<br>T2D Met+ 20,<br>IGT 49,<br>Norm 43</td><td>20</td><td>17</td><td>8 | 10</td><td>15 (=5 age-based groups, each with 3 replicates)</td><td>Disturbed 29<br>Natural 55</td></tr>
<tr><td>Df</td><td>152</td><td>224</td><td>55</td><td>54</td><td>3</td><td>18</td><td>15</td><td>N/A | N/A</td><td>N/A</td><td>82</td></tr>
<tr><td>Lipids</td><td>(+)***<br>W = 4053</td><td>(+)***<br>W = 8281</td><td>ns<br>W = 400</td><td>(+)*<br>W = 533</td><td>ns<br>F = 1.89</td><td>ns<br>t = 0.91</td><td>(+)*<br>t = 1.89</td><td>ns | ns<br>z = 1.54 | -1.09<br>tau = 0.46 | -0.28</td><td>(+)**<br>z = 3.14<br>tau = 0.64</td><td>(+)***<br>t = -5.51</td></tr>
<tr><td>Proteins</td><td>ns<br>t = -0.58</td><td>(+)**<br>W = 6520</td><td>(–)*<br>t = -1.79</td><td>(–)***<br>t = -3.36</td><td>ns<br>F = 2.07</td><td>ns<br>t = 0.25</td><td>ns<br>t = 0.97</td><td>ns | ns<br>z = 0 | -0.54<br>tau = 0 | -0.14</td><td>(–)*<br>z = -2.43<br>tau = -0.49</td><td>(–)***<br>t = 4.30</td></tr>
<tr><td>Amino sugars</td><td>ns<br>W = 2504</td><td>ns<br>W = 6093</td><td>(+)**<br>t = 2.51</td><td>(+)***<br>W = 581</td><td>ns<br>F = 0.93</td><td>(+)*<br>t = 1.80</td><td>(+)<br>t = 1.72</td><td>ns | ns<br>z = 0 | -0.18<br>tau = 0 | -0.05</td><td>ns<br>z = -1.22<br>tau = -0.25</td><td>ns<br>t = -0.88</td></tr>
<tr><td>Carbohydrates</td><td>(–)***<br>W = 1587</td><td>(–)***<br>W = 2502</td><td>ns<br>t = 0.33</td><td>ns<br>t = -0.28</td><td>ns<br>F = 1.99</td><td>ns<br>t = -0.84</td><td>(–)*<br>t = -3.06</td><td>(–) | (–)<br>z = -1.80 | -1.82<br>tau = -0.54 | -0.46</td><td>(–)***<br>z = -4.25<br>tau = -0.86</td><td>(–)***<br>W = 1385</td></tr>
<tr><td>Condensed aromatics</td><td>(+)***<br>t = 6.28</td><td>(+)***<br>t = 8.80</td><td>ns<br>t = 1.02</td><td>(+)*<br>t = 1.92</td><td>ns<br>F = 0.86</td><td>ns<br>t = 0.90</td><td>ns<br>t = 0.58</td><td>ns | ns<br>z = -0.26 | 1.27<br>tau = -0.08 | 0.32</td><td>(+)***<br>z = 3.44<br>tau = 0.70</td><td>(+)***<br>t = -4.30</td></tr>
<tr><td>Lignin</td><td>(+)**<br>W = 3438</td><td>(+)***<br>W = 7633</td><td>ns<br>t = 0.59</td><td>(+)**<br>t = 2.80</td><td>ns<br>F = 1.55</td><td>ns<br>t = 0.14</td><td>ns<br>t = 0.78</td><td>ns | ns<br>z = -0.51 | 0.54<br>tau = -0.15 | 0.14</td><td>(–)**<br>z = -3.14<br>tau = -0.64</td><td>(–)***<br>t = 7.91</td></tr>
<tr><td>Tannins</td><td>ns<br>W = 2422</td><td>(+)**<br>t = 3.02</td><td>(–)**<br>t = -2.30</td><td>(–)***<br>W = 176</td><td>ns<br>F = 0.85</td><td>ns<br>t = 0.02</td><td>(+)<br>t = 1.38</td><td>(+)* | ns<br>z = 2.05 | 1.09<br>tau = 0.62 | 0.28</td><td>ns<br>z = -0.30<br>tau = -0.06</td><td>(–)***<br>W = 1193</td></tr>
<tr><td>Other methylated compounds</td><td>ns<br>W = 3000</td><td>ns<br>W = 4791</td><td>ns<br>t = 1.34</td><td>ns<br>t = 0.89</td><td>(–)* IGT cf. Normal<br>F = 2.77</td><td>(+)<br>t = 3.12</td><td>ns<br>t = 0.81</td><td>ns | ns<br>z = 0 | -0.36<br>tau = 0 | -0.09</td><td>(–)***<br>z = -3.75<br>tau = -0.76</td><td>(–)***<br>t = 5.78</td></tr>
<tr><td>Other hydrated compounds</td><td>(+)*<br>W = 3115</td><td>(+)***<br>W = 7980</td><td>ns<br>t = 1.09</td><td>ns<br>t = -1.17</td><td>ns<br>F = 1.64</td><td>ns<br>t = -0.73</td><td>ns<br>t = 1.15</td><td>ns | (–)**<br>z = 1.28 | -3.27<br>tau = 0.39 | -0.83</td><td>ns<br>z = 0.81<br>tau = 0.16</td><td>(–)***<br>t = 5.20</td></tr>
<tr><td>Other demethylated compounds</td><td>ns<br>W = 2942</td><td>(+)***<br>W = 8113</td><td>ns<br>W = 455</td><td>ns<br>W = 479</td><td>ns<br>F = 1.08</td><td>ns<br>t = -0.21</td><td>(+)*<br>t = 2.31</td><td>ns | ns<br>z = 1.03 | -0.54<br>tau = 0.31 | -0.14</td><td>ns<br>z = 1.01<br>tau = 0.21</td><td>(–)*<br>W = 1037</td></tr>
<tr><td>Other condensed compounds</td><td>ns<br>t = 1.52</td><td>(–)*<br>t = -1.98</td><td>(+)*<br>t = 1.95</td><td>(+)**<br>t = 2.56</td><td>ns<br>F = 0.40</td><td>ns<br>t = 0.06</td><td>ns<br>t = -1.04</td><td>ns | ns<br>z = 0.26 | 1.63<br>tau = 0.08 | 0.41</td><td>ns<br>z = 1.52<br>tau = 0.31</td><td>(+)**<br>t = -2.81</td></tr>
<tr><td>Near VitK2-VitB12 compounds</td><td>(+)*<br>t = 2.30</td><td>ns<br>t = 0.81</td><td>ns<br>t = 0.19</td><td>(+)***<br>W = 587</td><td>(–)* IGT cf. Normal<br>F = 3.55</td><td>(+)***<br>t = 4.13</td><td>ns<br>t = -0.56</td><td>(+)* | (–)*<br>z = 2.05 | -2.18<br>tau = 0.62 | -0.55</td><td>(–)*<br>z = 2.43<br>tau = 0.49</td><td>ns<br>t = -1.35</td></tr>
<tr><td>All compounds</td><td>(–)***<br>W = 1675</td><td>ns<br>W = 5245</td><td>ns<br>t = -1.07</td><td>(–)**<br>W = 223</td><td>(–)* Normal cf. others<br>F = 2.84</td><td>ns<br>t = -0.34</td><td>ns<br>t = 0.49</td><td>(+)* | ns<br>z = 2.05 | -1.27<br>tau = 0.62 | -0.32</td><td>(–)<br>z = -1.82<br>tau = -0.37</td><td>(–)***<br>W = 1323</td></tr>
<tr><td>Total functional % relative abundances mapped into vK space (sample level)</td><td colspan="2">Range:<br>54.4–84.1%<br>Mean ± s.d.:<br>63.2 ± 3.5 %</td><td colspan="2">Range:<br>57.2–74.9%<br>Mean ± s.d.:<br>66.7 ± 2.7 %</td><td>Range:<br>55.8–69.5%<br>Mean ± s.d.:<br>64.4 ± 2.2 %</td><td colspan="2">Range:<br>51.7–57.0%<br>Mean ± s.d.:<br>54.9 ± 1.3 %</td><td>Range:<br>60.9–62.5%<br>Mean ± s.d.:<br>61.7 ± 0.3 %</td><td>Range:<br>61.3–62.3%<br>Mean ± s.d.:<br>61.9 ± 0.3 %</td><td>Range:<br>58.9–62.8%<br>Mean ± s.d.:<br>60.8 ± 1.0 %</td></tr>
</table>

Table Notes – see next page

Table Notes:

(+)/(–) = Increasing / decreasing pattern in the direction specified.

Significance: *** = $P$-value < 0.001, ** = 0.001 ≤ $P$-value < 0.01, * = 0.01 ≤ $P$-value ≤ 0.05, ns = $P$-value > 0.5.

† T-tests were used for testing differences between two groups (t values provided), except when pre-testing for homogeneity of variances between samples was significant, in which case we used the Wilcoxon rank sum test (W values provided).

‡ In testing differences between three or more groups Levene's test was first used to check for common variance among samples, followed by analysis of variance (F values reported), then Tukey's 'Honest Significant Difference' method for pairwise differences. Except where sample variance was different, then Kruskal-Wallis rank sum tests were used ($\chi^2$ values reported), followed by post-hoc Dunn tests to detect pairwise differences.

§ Associations with numeric response data were assessed using Pearson r correlation tests.

¶ Associations with ordinal response data were assessed using Kendall tau correlation tests.

### Marginally significant indicative correlations in soil samples and problem behavior data are reported by sign only where abs(Kendall tau or Pearson r) > 0.3 and P-value < 0.1.

|| PCaN soil data are analyzed separately in pH classes: 1) strongly acidic (pH < 4.5), and 2) acidic to neutral (4.5 ≤ pH ≤ 7)

Abbreviations:

ACVD = atherosclerotic cardiovascular disease

AMI = Australian Microbiome Initiative

cf. = compared to

CPP = compound processing potential

Df = Degrees of freedom

IGT = impaired glucose tolerance

N/A = not applicable

ns = no significant difference

PC1 = first principal component of problem (anxious, depressive, etc.) behaviors

PCaN = People Cities and Nature urban ecosystem restoration research program

T2D = type 2 diabetes

**Table S4.** Summary results for $CPP_{density}$ data, i.e., log10-transformed function % relative abundance per unit vK area, mapped within 0.05 vK unit buffer radii of key biomolecules. Corresponding plots are noted. Human gut results indicate increased (+) or decreased (–) compound processing potential ($CPP_{density}$) observed in the disease state compared to the normal healthy state. Soil results indicate increased (+) or decreased (–) $CPP_{density}$ values with ecosystem maturity.

| | Human gut | | | | | | | Soils | | |
|---|---|---|---|---|---|---|---|---|---|---|
| | ACVD cf. Normal (Jie et al., 2017) (Fig. S6) [†] | | Colorectal cancer cf. Normal (Zeller et al., 2014) (Fig. S10) [†] | | T2D Met- / T2D Met+/ IGT/ Normal (Forslund et al., 2015) (Fig. S14) [‡] | Trend with PC1 of problem behaviors (Flannery et al., 2020) (Fig. S20) [§, #] | | PCaN: Trend with Reveg age (Barnes et al., 2020) (Fig. S28) [¶, #] | Post-mining: Trend with Reveg age (Sun and Badgley, 2019) (Fig. S32) [¶, #] | AMI: Natural cf. disturbed (Bissett et al., 2016) (Fig. S35) [†] |
| Sex / pH class | F | M | F | M | F | F | M | 1) \| 2) [‖] | | |
| N | ACVD 53 Norm 101 | ACVD 157 Norm 69 | Cancer 24 Normal 33 | Cancer 29 Normal 27 | T2D Met- 33, T2D Met+ 20, IGT 49, Norm 43 | 23 | 17 | 8 \| 10 | 15 (=5 age-based groups, each with 3 replicates) | Disturbed 29 Natural 55 |
| Df. | | | 55 | 54 | 3 | 18 | 15 | 6 \| 8 | | 82 |
| Acetate | ns W = 2888 | (–)*** W = 3418 | ns t = -0.33 | ns W = 403 | ns F = 0.70 | ns t = -0.35 | (–)** t = -3.15 | ns \| (–)** z = 0.77 \| -2.72 tau = 0.23 \| -0.69 | (–)** z = -2.73 tau = -0.56 | (–)*** t = 4.54 |
| Propionate | (+)* W = 3228 | ns W = 6117 | ns t = -1.01 | ns t = -0.62 | ns F = 0.68 | (+) t = 1.40 | ns t = 0.90 | ns \| ns z = 0.26 \| 1.27 tau = 0.08 \| 0.32 | ns z = 0.61 tau = 0.12 | (–)*** t = 7.23 |
| Butyrate | N/A | N/A | N/A | N/A | N/A | N/A | N/A | N/A \| N/A | N/A | N/A |
| VitB2 | ns W = 2823 | ns W = 5379 | ns t = 0.54 | (+)* t = 2.15 | ns F = 0.79 | ns t = 0.66 | ns t = -0.41 | (–) \| ns z = -1.80 \| -0.91 tau = -0.54 \| -0.23 | (–)** z = -3.14 tau = -0.64 | ns W = 772 |
| VitB12 | (+)*** W = 3626 | ns W = 6154 | (+)** t = 2.73 | (+)** t = 3.20 | ns F = 1.99 | ns t = 1.32 | (–)* t = -2.31 | (–)* \| ns z = -2.05 \| 0.36 tau = -0.62 \| 0.09 | (–)* z = -2.43 tau = -0.49 | (–)*** t = 6.30 |
| VitB6 | ns W = 3047 | (+)*** W = 7352 | ns W = 448 | ns W = 395 | ns F = 2.14 | ns t = 0.34 | (+)* t = 1.78 | ns \| ns z = 1.28 \| -1.45 tau = 0.39 \| -0.37 | (–)** z = -3.24 tau = -0.66 | (–)*** W = 1384 |
| VitB9 | (+)*** W = 3803 | (+)*** W = 7416 | (+)* t = 2.20 | (+)** t = 2.91 | ns F = 0.43 | ns t = -0.69 | (+)* t = 2.20 | (+) \| ns z = 1.80 \| 0.18 tau = 0.54 \| 0.05 | (+) z = 1.92 tau = 0.39 | (+)*** t = -4.36 |
| VitK2 | (+)*** W = 4209 | (+)*** t = 5.42 | ns W = 396 | ns W = 419 | ns $\chi^2$ = 7.15 | ns t = 0.22 | N/A | (+) \| ns z = 1.80 \| 0.36 tau = 0.54 \| 0.09 | (+)* z = 2.33 tau = 0.47 | (+)*** W = 305 |
| Glutamate | (–)*** W = 1758 | (–)*** W = 2641 | ns t = 1.01 | ns t = 0.89 | ns F = 0.38 | ns t = 0.23 | (–)** t = -2.77 | ns \| (–)* z = 0 \| -2.36 tau = 0 \| -0.60 | (–)*** z = -3.34 tau = -0.68 | (–)*** t = 3.32 |
| Pyruvate | ns W = 2817 | (+)*** W = 7544 | ns W = 490 | (+)* W = 519 | ns F = 1.31 | ns t = -0.37 | ns t = -0.68 | ns \| ns z = 0.77 \| -1.45 tau = 0.23 \| -0.37 | ns z = -1.52 tau = -0.31 | (–)*** W = 1188 |

Table Note: Symbols and abbreviations are detailed as per Table S3.

**Table S5.** Summary results for putative activity-normalized, or amino sugar-adjusted log ratio ($CPP_{ASALR}$) data as displayed in Figs. 3, 5 of the main paper. Human gut results indicate increased (+) or decreased (–) compound processing potential ($CPP_{ASALR}$) observed in the disease state compared to the normal healthy state. Soil results indicate increased (+) or decreased (–) $CPP_{ASALR}$ values with ecosystem maturity. Samples with zero measure of $CPP_{class\ =\ amino\ sugars}$ were excluded due to undefined values (division by zero).

| | Human gut | | | | | | | Soils | | |
|---|---|---|---|---|---|---|---|---|---|---|
| | ACVD cf. Normal (Jie et al., 2017) † | | Colorectal cancer cf. Normal (Zeller et al., 2014) † | | T2D Met- / T2D Met+/ IGT/ Normal (Forslund et al., 2015) ‡ | High cf. Low PC1 of problem behaviors (Flannery et al., 2020) † | | PCaN: Trend with Reveg age (Barnes et al., 2020) ¶, # | Post-mining: Trend with Reveg age (Sun and Badgley, 2019) ¶, # | AMI: Natural cf. disturbed (Bissett et al., 2016) † |
| Sex / pH class | F | M | F | M | F | F | M | 1) \| 2) ‖ | | |
| N | ACVD 53 Norm 99 | ACVD 157 Norm 66 | Cancer 24 Normal 33 | Cancer 29 Normal 27 | T2D Met- 33, T2D Met+ 20, IGT 49, Norm 43 | High PC1 10, Low PC1 10 | High PC1 8, Low PC1 9 | 8 \| 10 | 15 (=5 age-based groups, each with 3 replicates) | Disturbed 29 Natural 55 |
| Df | 152 | 224 | 55 | 54 | 3 | 18 | 15 | N/A \| N/A | N/A | 82 |
| Lipids | (+)*** W = 3910 | (+)*** W = 7589 | ns W = 373 | (+)* W = 499 | ns F = 1.15 | ns t = 0.15 | ns t = 1.36 | (+) \| ns z = 1.80 \| -0.73 tau = 0.54 \| -0.18 | (+)** z = 3.04 tau = 0.62 | (+)*** t = -5.29 |
| Proteins | ns W = 2546 | ns W = 5805 | (–)** t = -2.43 | (–)*** t = -4.09 | ns F = 1.81 | ns t = 0.05 | ns t = -0.40 | ns \| ns z = 0.77 \| -0.73 tau = 0.23 \| -0.18 | (–) z = -1.92 tau = -0.39 | (–)*** t = 4.65 |
| Carbohydrates | (–)** W = 1880 | (–)*** W = 2866 | ns t = -1.55 | (–)** W = 225 | ns F = 1.45 | ns t = -0.74 | (–)* t = -1.85 | ns \| (–)* z = -0.51 \| -2.0 tau = -0.15 \| -0.51 | (–)* z = -2.43 tau = -0.49 | (–)*** t = 4.56 |
| Condensed aromatics | (+)*** W = 4036 | (+)*** W = 7989 | ns t = 0.81 | ns t = 0.59 | ns F = 0.79 | ns t = -0.48 | ns t = 0.31 | ns \| ns z = 0 \| 1.45 tau = 0 \| 0.37 | (+)*** z = 3.34 tau = 0.68 | (+)*** t = -3.44 |
| Lignin | (+)* W = 3061 | (+)* W = 6152 | (–)* t = -2.0 | (–)* W = 283 | ns F = 1.33 | ns t = -0.64 | ns t = 0.35 | ns \| ns z = 0 \| 0 tau = 0 \| 0 | ns z = -0.20 tau = -0.04 | (–)*** t = 3.95 |
| Tannins | ns W = 2300 | ns W = 5416 | (–)** t = -2.86 | (–)*** W = 170 | ns F = 0.44 | ns t = -0.38 | ns t = 0.44 | ns \| ns z = 0.51 \| 0.73 tau = 0.15 \| 0.18 | ns z = 0.81 tau = 0.16 | (–)* W = 995 |
| Other methylated compounds | (+)* W = 3093 | ns W = 4671 | ns t = -0.33 | (–)* W = 260 | ns F = 1.41 | (+)* t = 1.81 | ns t = 0.21 | ns \| ns z = 0.26 \| -0.91 tau = 0.08 \| -0.23 | (–)*** z = -4.05 tau = -0.82 | (–)*** t = 5.13 |
| Other hydrated compounds | (+)* W = 3084 | (+)*** W = 7445 | ns t = -1.13 | (–)** W = 210 | ns F = 2.31 | ns t = -0.68 | ns t = 0.37 | ns \| ns z = 0 \| -1.45 tau = 0 \| -0.37 | ns z = 1.11 tau = 0.23 | (–)*** W = 1178 |
| Other demethylated compounds | ns W = 2825 | (+)*** W = 7404 | (–)* t = -1.86 | (–)** W = 216 | ns F = 2.40 | ns t = -0.37 | ns t = 1.18 | ns \| ns z = 0.26 \| 0.18 tau = 0.08 \| 0.05 | ns z = 1.11 tau = 0.23 | (–)* W = 999 |
| Other condensed compounds | ns W = 2890 | (–)** W = 3835 | ns t = 0.83 | ns t = 1.03 | ns F = 0.19 | ns W = 41 | ns t = -1.25 | ns \| ns z = 0.51 \| 1.63 tau = 0.15 \| 0.41 | ns z = 1.52 tau = 0.31 | (+)** t = -2.53 |
| Near VitK2-VitB12 compounds | (+)* W = 3199 | ns W = 4778 | ns W = 383 | ns W = 486 | (–)* IGT cf. Normal & T2D Met+ F = 4.34 | ns t = 1.36 | ns t = -0.29 | ns \| (–) z = 1.54 \| -1.82 tau = 0.46 \| -0.46 | (+)* z = 2.43 tau = 0.49 | ns W = 652 |

Table Note: Symbols and abbreviations are detailed as per Table S3.

**Atherosclerotic Cardiovascular disease (ACVD) case study data**

**Table S6.** PERMANOVA and beta-dispersion results testing for differences in weighted-mean vK coordinates between ACVD (n = 53) and normal (n = 101) diagnoses, in female subjects.

| | PERMANOVA | | | | Beta-dispersion | | |
|---|---|---|---|---|---|---|---|
| Compound class | Df | $R^2$ | F-value | P-value | Df | F-value | P-value |
| Lipids | 1 | 0.057 | 8.34 | **0.005** ** | 1 | 1.11 | 0.307 (ns) |
| Proteins | 1 | 0.010 | 1.56 | 0.199 (ns) | 1 | 10.1 | **0.002** ** |
| Amino sugars | 1 | 0.022 | 3.45 | 0.052 (ns) | 1 | 6.33 | **0.017** * |
| Carbohydrates | 1 | 0.059 | 9.52 | **0.003** ** | 1 | 5.54 | **0.026** * |
| Condensed aromatics | 1 | 0.030 | 4.32 | **0.035** * | 1 | 1.33 | 0.264 (ns) |
| Lignin | 1 | 0.021 | 3.21 | **0.038** * | 1 | 9.45 | **0.004** ** |
| Tannins | 1 | 0.089 | 14.8 | **0.001** *** | 1 | 10.35 | **0.004** ** |

**Table S7.** PERMANOVA and beta-dispersion results testing for differences in weighted-mean vK coordinates between ACVD (n = 157) and normal (n = 69) diagnoses, in male subjects.

| | PERMANOVA | | | | Beta-dispersion | | |
|---|---|---|---|---|---|---|---|
| Compound class | Df | $R^2$ | F-value | P-value | Df | F-value | P-value |
| Lipids | 1 | 0.0032 | 0.691 | 0.447 (ns) | 1 | 8.24 | **0.005** ** |
| Proteins | 1 | 0.057 | 13.4 | **0.001** *** | 1 | 23.6 | **0.001** *** |
| Amino sugars | 1 | 0.04 | 9.26 | **0.001** *** | 1 | 18.5 | **0.001** *** |
| Carbohydrates | 1 | 0.225 | 65 | **0.001** *** | 1 | 7.08 | **0.012** * |
| Condensed aromatics | 1 | 0.104 | 25.0 | **0.001** *** | 1 | 2.48 | 0.114 (ns) |
| Lignin | 1 | 0.037 | 8.52 | **0.001** *** | 1 | 27.1 | **0.001** *** |
| Tannins | 1 | 0.203 | 57.0 | **0.001** *** | 1 | 50.9 | **0.001** *** |

#### Colorectal cancer case study data

**Table S8.** PERMANOVA and beta-dispersion results testing for differences in weighted-mean vK coordinates between colorectal cancer (n = 24) and normal (n = 33) diagnoses, in female subjects.

| | PERMANOVA | | | | Beta-dispersion | | |
|---|---|---|---|---|---|---|---|
| Compound class | Df | $R^2$ | F-value | P-value | Df | F-value | P-value |
| Lipids | 1 | 0.075 | 3.97 | 0.068 (ns) | 1 | 0.05 | 0.818 (ns) |
| Proteins | 1 | 0.003 | 0.175 | 0.717 (ns) | 1 | 5.23 | **0.018** * |
| Amino sugars | 1 | 0.027 | 1.516 | 0.217 (ns) | 1 | 0.366 | 0.563 (ns) |
| Carbohydrates | 1 | 0.023 | 1.31 | 0.252 (ns) | 1 | 1.588 | 0.228 (ns) |
| Condensed aromatics | 1 | 0.062 | 3.37 | **0.05** * | 1 | 6.74 | **0.011** * |
| Lignin | 1 | 0.027 | 1.55 | 0.212 (ns) | 1 | 0.0045 | 0.94 (ns) |
| Tannins | 1 | 0.033 | 1.89 | 0.156 (ns) | 1 | 1.60 | 0.224 (ns) |

**Table S9.** PERMANOVA and beta-dispersion results testing for differences in weighted-mean vK coordinates between colorectal cancer (n = 29) and normal (n = 27) diagnoses, in male subjects.

| | PERMANOVA | | | | Beta-dispersion | | |
|---|---|---|---|---|---|---|---|
| Compound class | Df | $R^2$ | F-value | P-value | Df | F-value | P-value |
| Lipids | 1 | 0.031 | 1.55 | 0.216 (ns) | 1 | 3.10 | 0.093 (ns) |
| Proteins | 1 | 0.006 | 0.303 | 0.609 (ns) | 1 | 4.34 | **0.04** * |
| Amino sugars | 1 | 0.051 | 2.89 | 0.075 (ns) | 1 | 3.154 | 0.066 (ns) |
| Carbohydrates | 1 | 0.017 | 0.937 | 0.377 (ns) | 1 | 1.106 | 0.286 (ns) |
| Condensed aromatics | 1 | 0.041 | 2.01 | 0.165 (ns) | 1 | 5.13 | **0.018** * |
| Lignin | 1 | 0.101 | 6.03 | **0.005** ** | 1 | 0.319 | 0.572 |
| Tannins | 1 | 0.111 | 6.75 | **0.008** ** | 1 | 1.93 | 0.185 (ns) |

#### Type 2 diabetes case study data

**Table S10.** PERMANOVA and beta-dispersion results testing for differences in weighted-mean vK coordinates between T2D Met- (n = 33), T2D Met+ (n = 20), IGT (n = 49), and Normal (n = 43) in female subjects only.

| | PERMANOVA | | | | Beta-dispersion | | |
|---|---|---|---|---|---|---|---|
| Compound class | Df | $R^2$ | F-value | P-value | Df | F-value | P-value |
| Lipids | 3 | 0.039 | 1.92 | 0.128 (ns) | 3 | 1.71 | 0.158 (ns) |
| Proteins | 3 | 0.070 | 3.52 | **0.013** * | 3 | 1.09 | 0.349 (ns) |
| Amino sugars | 3 | 0.030 | 1.44 | 0.215 (ns) | 3 | 0.564 | 0.63 (ns) |
| Carbohydrates | 3 | 0.009 | 0.4223 | 0.716 (ns) | 3 | 0.836 | 0.46 (ns) |
| Condensed aromatics | 3 | 0.008 | 0.363 | 0.841 (ns) | 3 | 0.561 | 0.646 (ns) |
| Lignin | 3 | 0.039 | 1.89 | 0.07 (ns) | 3 | 0.899 | 0.431 (ns) |
| Tannins | 3 | 0.036 | 1.75 | 0.146 (ns) | 3 | 0.078 | 0.974 (ns) |

**Problem behavior case study data**

**Table S11.** PERMANOVA and beta-dispersion results testing for differences in weighted-mean vK coordinates between low (n = 10) and high (n = 10) PC1 of problem behaviors, in female subjects.

| | PERMANOVA | | | | Beta-dispersion | | |
|---|---|---|---|---|---|---|---|
| Compound class | Df. | $R^2$ | F-value | P-value | Df | F-value | P-value |
| Lipids | 1 | 0.009 | 0.163 | 0.73 (ns) | 1 | 0.098 | 0.755 (ns) |
| Proteins | 1 | 0.048 | 0.916 | 0.377 (ns) | 1 | 1.94 | 0.183 (ns) |
| Amino sugars | 1 | 0.035 | 0.660 | 0.429 (ns) | 1 | 0.099 | 0.777 (ns) |
| Carbohydrates | 1 | 0.0008 | 0.014 | 0.956 (ns) | 1 | 4.10 | 0.057 (.) |
| Condensed aromatics | 1 | 0.007 | 0.123 | 0.812 (ns) | 1 | 4.21 | **0.046 *** |
| Lignin | 1 | 0.025 | 0.467 | 0.631 (ns) | 1 | 0.194 | 0.718 (ns) |
| Tannins | 1 | 0.064 | 1.23 | 0.298 (ns) | 1 | 0.488 | 0.491 (ns) |

**Table S12.** PERMANOVA and beta-dispersion results testing for differences in weighted-mean vK coordinates between low (n = 9) and high (n = 8) PC1 of problem behaviors, in male subjects.

| | PERMANOVA | | | | Beta-dispersion | | |
|---|---|---|---|---|---|---|---|
| Compound class | Df | $R^2$ | F-value | P-value | Df | F-value | P-value |
| Lipids | 1 | 0.045 | 0.706 | 0.431 (ns) | 1 | 1.82 | 0.213 (ns) |
| Proteins | 1 | 0.158 | 2.80 | 0.094 (.) | 1 | 0.151 | 0.724 (ns) |
| Amino sugars | 1 | 0.075 | 1.21 | 0.253 (ns) | 1 | 0.0008 | 0.984 (ns) |
| Carbohydrates | 1 | 0.109 | 1.84 | 0.205 (ns) | 1 | 0.231 | 0.641 (ns) |
| Condensed aromatics | 1 | 0.018 | 0.277 | 0.649 (ns) | 1 | 0.835 | 0.37 (ns) |
| Lignin | 1 | 0.005 | 0.078 | 0.887 (ns) | 1 | 3.60 | 0.067 (.) |
| Tannins | 1 | 0.052 | 0.815 | 0.432 (ns) | 1 | 0.299 | 0.602 (ns) |

**People Cities and Nature (PCaN) urban ecosystem restoration case study data**

**Table S13.** PERMANOVA results testing for trend in weighted-mean vK coordinates for PCaN urban ecosystem restoration (strongly acidic soils) based on ordinal (ranked) values of vegetation age.

| | PERMANOVA | | | | Beta-dispersion | | |
|---|---|---|---|---|---|---|---|
| variable | Df | $R^2$ | F-value | P-value | Df | F-value | P-value |
| Lipids | 1 | 0.005 | 0.032 | 0.912 (ns) | Beta dispersion values were not tested due to low sample sizes | | |
| Proteins | 1 | 0.062 | 0.399 | 0.529 (ns) | | | |
| Amino sugars | 1 | 0.011 | 0.067 | 0.813 (ns) | | | |
| Carbohydrates | 1 | 0.085 | 0.559 | 0.628 (ns) | | | |
| Condensed aromatics | 1 | 0.003 | 0.017 | 0.919 (ns) | | | |
| Lignin | 1 | 0.303 | 2.61 | 0.139 (ns) | | | |
| Tannins | 1 | 0.194 | 1.44 | 0.288 (ns) | | | |

**Table S14.** PERMANOVA results testing for differences in weighted-mean vK coordinates for PCaN urban ecosystem restoration (acidic to neutral soils) based on ordinal (ranked) values of vegetation age.

| | PERMANOVA | | | | Beta-dispersion | | |
|---|---|---|---|---|---|---|---|
| variable | Df | $R^2$ | F-value | P-value | Df | F-value | P-value |
| Lipids | 1 | 0.039 | 0.328 | 0.653 (ns) | Beta dispersion values were not tested due to low sample sizes | | |
| Proteins | 1 | 0.122 | 1.11 | 0.344 (ns) | | | |
| Amino sugars | 1 | 0.016 | 0.134 | 0.733 (ns) | | | |
| Carbohydrates | 1 | 0.569 | 10.57 | **0.008 **** | | | |
| Condensed aromatics | 1 | 0.541 | 9.44 | **0.019 *** | | | |
| Lignin | 1 | 0.086 | 0.750 | 0.407 (ns) | | | |
| Tannins | 1 | 0.238 | 2.50 | 0.117 (ns) | | | |

#### Post-mining forest restoration case study data

**Table S15.** PERMANOVA and beta-dispersion results testing for differences in weighted-mean vK coordinates for post-mining forest ecosystem restoration soils across age-based groups (n = 5 groups of 3 each).

| | PERMANOVA | | | | Beta-dispersion | | |
|---|---|---|---|---|---|---|---|
| variable | Df | $R^2$ | F-value | P-value | Df | F-value | P-value |
| Lipids | 4 | 0.655 | 4.75 | **0.014 *** | 4 | 0.141 | 0.944 (ns) |
| Proteins | 4 | 0.884 | 19.0 | **0.002 **** | 4 | 0.812 | 0.546 (ns) |
| Amino sugars | 4 | 0.854 | 14.6 | **0.01 **** | 4 | 0.515 | 0.74 (ns) |
| Carbohydrates | 4 | 0.607 | 3.86 | **0.008 **** | 4 | 0.165 | 0.947 (ns) |
| Condensed aromatics | 4 | 0.925 | 30.7 | **0.002 **** | 4 | 0.5661 | 0.686 (ns) |
| Lignin | 4 | 0.841 | 13.2 | **0.001 ***** | 4 | 0.181 | 0.942 (ns) |
| Tannins | 4 | 0.795 | 9.70 | **0.001 ***** | 4 | 1.01 | 0.46 (ns) |

#### Australian Microbiome Initiative (AMI) disturbed vs. natural soil case study data

**Table S16.** PERMANOVA and beta-dispersion results testing for differences in weighted-mean vK coordinates between AMI disturbed (n = 29) and natural (n = 55) soils.

| | PERMANOVA | | | | Beta-dispersion | | |
|---|---|---|---|---|---|---|---|
| variable | Df | $R^2$ | F-value | P-value | Df | F-value | P-value |
| Lipids | 1 | 0.192 | 19.5 | **0.001 ***** | 1 | 0.023 | 0.887 (ns) |
| Proteins | 1 | 0.112 | 10.3 | **0.001 ***** | 1 | 13.04 | **0.002 **** |
| Amino sugars | 1 | 0.266 | 29.7 | **0.001 ***** | 1 | 8.30 | **0.008 **** |
| Carbohydrates | 1 | 0.153 | 14.83 | **0.001 ***** | 1 | 0.207 | 0.671 (ns) |
| Condensed aromatics | 1 | 0.195 | 19.84 | **0.001 ***** | 1 | 3.81 | 0.056 (ns) |
| Lignin | 1 | 0.065 | 5.69 | **0.014 *** | 1 | 0.485 | 0.506 (ns) |
| Tannins | 1 | 0.091 | 8.24 | **0.002 **** | 1 | 4.21 | **0.039 *** |

**Table S17.** Summary statistics for CCLasso-correlation relationship networks based on compound-associated van Krevelen coordinate profiles of gut metagenomes from the type 2 diabetes (T2D) case study (see Fig. 4 in main article, and Fig. S17). Diagnosis groups of normal healthy, impaired glucose tolerance (IGT), and T2D with and without Metformin (Met+, Met-) (n = 20 female subjects in each group) are compared to bootstrap (B = 1000) randomized network probability density distributions. Significant results ($P \leq 0.05$) are in bold; borderline results in italics; based on comparison to the bootstrap distribution. Network analyses were based on commonly observed vK coordinates (specified by presence in at least 60% of samples and minimum of 2% total functional relative abundance summed across samples).

| Diagnosis group | No. of vertices (nodes) | No. of edges (links) | Edge density | Fraction of negative edges | Degree centralization | Closeness centralization | Betweenness centralization | Mean distance | Modularity |
|---|---|---|---|---|---|---|---|---|---|
| Normal | 48 | 96 | 0.0851 | **0.1042*** | **0.5106*** | 1.1099 | 0.3803 | 2.4807 | 0.2176 |
| IGT | 49 | 67 | 0.0570 | 0.3433 | 0.2347 | 1.3493 | 0.3883 | 3.6220 | 0.5939 |
| T2D Met– | *173* | 4028 | 0.2707 | *0.4814* | 0.1944 | **0.1647*** | *0.0101* | *1.7666* | 0.2610 |
| T2D Met+ | 103 | 352 | 0.0670 | 0.3295 | 0.1879 | 1.3976 | 0.3869 | 3.4421 | 0.4625 |
| 2.5th, 97.5th percentiles of bootstrap distribution (B=1000) | 17, 174 | 13, 4524 | 0.0404, 0.3194 | 0.1291, 0.4971 | 0.1075, 0.4656 | 0.1867, 1.4243 | 0.0098, 0.5352 | 1.6674, 4.1135 | 0.0645, 0.7189 |

**Table S18.** Summary of differentially abundant functions, comparing **female** subjects with high (n = 10) versus low (n = 10) PC1 of problem behaviors

| Fxn ID | ANCOMBC Intercept | ANCOMBC Log fold change High PC1 cf. Low PC1 | OC_x | HC_y | Compound-associated vK mapping zone | SEED Subsytem Level 1 | SEED Subsytem Level 2 | SEED Subsytem Level 3 | Function |
|---|---|---|---|---|---|---|---|---|---|
| fxn_118 | 4.60E-16 | -0.5108 | 0.6539 | 1.7471 | Amino sugars | Amino Acids and Derivatives | Alanine, serine, and glycine | Glycine Biosynthesis | Serine_hydroxymethyltransferase_(EC_2.1.2.1) |
| fxn_341 | 2.44E-16 | 0.5932 | 0.7879 | 1.9606 | Carbohydrates | Amino Acids and Derivatives | Arginine; urea cycle, polyamines | Polyamine Metabolism | Arginine_decarboxylase_(EC_4.1.1.19);_Lysine_decarboxylase_(EC_4.1.1.18);_Ornithine_decarboxylase_(EC_4.1.1.17) |
| fxn_532 | -3.84E-16 | 0.7338 | 1.1429 | 1.3571 | Other demethylated | Amino Acids and Derivatives | Aromatic amino acids and derivatives | Chorismate Synthesis | 3-dehydroquinate_synthase_(EC_4.2.3.4)_#_AroB |
| fxn_680 | -3.36E-16 | 0.7591 | 1.1429 | 1.3571 | Other demethylated | Amino Acids and Derivatives | Aromatic amino acids and derivatives | Common Pathway For Synthesis of Aromatic Compounds (DAHP synthase to chorismate) | 3-dehydroquinate_synthase_(EC_4.2.3.4)_#_AroB |
| fxn_2426 | -2.23E-16 | 0.4815 | 0.7663 | 0.9250 | Tannins | Carbohydrates | Central carbohydrate metabolism | acinetobacter tca | Succinate_dehydrogenase_iron-sulfur_protein_(EC_1.3.99.1) |
| fxn_4797 | 1.43E-16 | 0.6131 | 1.0357 | 1.2857 | Other demethylated | Carbohydrates | Organic acids | Glycerate metabolism | 2-hydroxy-3-oxopropionate_reductase_(EC_1.1.1.60) |
| fxn_5232 | -8.28E-17 | 0.5436 | NA | NA | NA | Cell Division and Cell Cycle | - | Bacterial Cytoskeleton | Chromosome_(plasmid)_partitioning_protein_ParB_/_Stage_0_sporulation_protein_J |
| fxn_6310 | -2.84E-16 | 0.7211 | NA | NA | NA | Cell Wall and Capsule | Gram-Negative cell wall components | Major Outer Membrane Proteins | Outer_membrane_chaperone_Skp_(OmpH)_precursor_@_Outer_membrane_protein_H_precursor |
| fxn_7446 | 3.69E-16 | 0.5585 | NA | NA | NA | Clustering-based subsystems | Probably GTP or GMP signaling related | CBSS-176299.4.peg.1292 | Ribonuclease_III_(EC_3.1.26.3) |
| fxn_7724 | -1.06E-16 | 0.6400 | 0.4672 | 1.4495 | Lignin | Cofactors, Vitamins, Prosthetic Groups, Pigments | - | CLO thiaminPP biosynthesis | 2-iminoacetate_synthase_(ThiH)_(EC_4.1.99.19) |
| fxn_8150 | -8.46E-17 | -0.7967 | 0.8910 | 1.4072 | Tannins | Cofactors, Vitamins, Prosthetic Groups, Pigments | - | Thiamin biosynthesis LDP | Sulfur_carrier_protein_adenylyltransferase_ThiF |
| fxn_8356 | -3.16E-16 | -0.7595 | 0.8910 | 1.4072 | Tannins | Cofactors, Vitamins, Prosthetic Groups, Pigments | - | Thiamin Copy RZ | Sulfur_carrier_protein_adenylyltransferase_ThiF |
| fxn_9046 | -1.72E-16 | 0.9616 | 0.3267 | 1.6467 | Proteins | Cofactors, Vitamins, Prosthetic Groups, Pigments | Folate and pterines | YgfZ | Biotin_synthase_(EC_2.8.1.6) |
| fxn_9072 | 4.20E-16 | 0.5429 | 0.7333 | 1.0310 | Tannins | Cofactors, Vitamins, Prosthetic Groups, Pigments | Folate and pterines | YgfZ | Dihydroorotate_dehydrogenase_(NAD(+)),_electron_transfer_subunit_(EC_1.3.1.14) |
| fxn_10516 | -6.29E-17 | -0.4833 | NA | NA | NA | DNA Metabolism | DNA repair | DNA repair, bacterial RecFOR pathway | RecA_protein |
| fxn_11346 | 3.54E-16 | -0.6258 | 0.7757 | 1.6784 | Carbohydrates | Fatty Acids, Lipids, and Isoprenoids | Isoprenoids | Isoprenoid Biosynthesis | Octaprenyl-diphosphate_synthase_(EC_2.5.1.-)_/_Dimethylallyltransferase_(EC_2.5.1.1)_/_Geranyltranstransferase_(farnesyldiphosphate_synthase)_(EC_2.5.1.10)_/_Geranylgeranyl_pyrophosphate_synthetase_(EC_2.5.1.29) |
| fxn_11530 | -4.29E-16 | 0.6331 | 0.6898 | 2.1205 | Amino sugars | Fatty Acids, Lipids, and Isoprenoids | Triacylglycerols | Triacylglycerol metabolism | Lysophospholipase_L2_(EC_3.1.1.5) |
| fxn_11660 | -2.71E-16 | 0.4524 | NA | NA | NA | Iron acquisition and metabolism | - | Iron acquisition in Streptococcus | Ferric_iron_ABC_transporter,_ATP-binding_protein |
| fxn_16658 | 7.76E-17 | -1.0632 | NA | NA | NA | Regulation and Cell signaling | Programmed Cell Death and Toxin-antitoxin Systems | Murein hydrolase regulation and cell death | Autolysis_histidine_kinase_LytS |
| fxn_18172 | -2.81E-16 | 0.6666 | 1.3333 | 1.1111 | Other demethylated | RNA Metabolism | RNA processing and modification | RNA pseudouridine syntheses | Ribosomal_large_subunit_pseudouridine(746)_synthase_(EC_5.4.99.29)_@_tRNA_pseudouridine(32)_synthase_(EC_5.4.99.28) |
| fxn_18378 | 3.64E-16 | 0.6693 | NA | NA | NA | RNA Metabolism | RNA processing and modification | Threonylcarbamoyladenosine | TsaC_protein_(YrdC-Sua5_domains)_required_for_threonylcarbamoyladenosine_t(6)A37_modification_in_tRNA |
| fxn_18462 | -3.82E-16 | 0.6840 | 1.2421 | 1.1930 | Other demethylated | RNA Metabolism | RNA processing and modification | tRNA modification Archaea | tRNA_pseudouridine(38-40)_synthase_(EC_5.4.99.12)_##_tRNA_Psi38,_Psi39_and_Psi40 |

**Table S19.** Summary of differentially abundant functions, comparing **male** subjects with high (n = 8) versus low (n = 9) PC1 of problem behaviors

| Fxn ID | ANCOMBC Intercept | ANCOMBC Log fold change High PC1 cf. Low PC1 | OC_x | HC_y | Compound-associated vK mapping zone | SEED Subsytem Level 1 | SEED Subsytem Level 2 | SEED Subsytem Level 3 | Function |
|---|---|---|---|---|---|---|---|---|---|
| fxn_303 | 0.0334 | 0.7756 | 0.4167 | 2.3333 | Other methylated | Amino Acids and Derivatives | Arginine; urea cycle, polyamines | Arginine Deiminase Pathway | Arginine_deiminase_(EC_3.5.3.6) |
| fxn_3075 | 0.0395 | 0.9227 | 0.4130 | 1.5697 | Proteins | Carbohydrates | Central carbohydrate metabolism | TCA Cycle | Dihydrolipoamide_dehydrogenase_of_branched-chain_alpha-keto_acid_dehydrogenase_(EC_1.8.1.4)_/_Dihydrolipoamide_dehydrogenase_(EC_1.8.1.4) |
| fxn_9200 | 0.0361 | 0.8402 | 0.8140 | 1.6031 | Carbohydrates | Cofactors, Vitamins, Prosthetic Groups, Pigments | Lipoic acid | BEY LIP | Glycine_dehydrogenase_[decarboxylating]_(glycine_cleavage_system_P2_protein)_(EC_1.4.4.2) |
| fxn_10191 | 0.0384 | 0.8960 | NA | NA | NA | Cofactors, Vitamins, Prosthetic Groups, Pigments | Tetrapyrroles | Coenzyme B12 biosynthesis | Duplicated_ATPase_component_CbrU_of_energizing_module_of_predicted_cobalamin_ECF_transporter_/_Substrate-specific_component_CbrT_of_predicted_cobalamin_ECF_transporter |
| fxn_11908 | 0.0384 | 0.8960 | NA | NA | NA | Membrane Transport | - | ECF class transporters | Duplicated_ATPase_component_CbrU_of_energizing_module_of_predicted_cobalamin_ECF_transporter_/_Substrate-specific_component_CbrT_of_predicted_cobalamin_ECF_transporter |
| fxn_14887 | 0.0313 | 0.7258 | NA | NA | NA | Phosphorus Metabolism | - | High affinity phosphate transporter and control of PHO regulon | response_regulator_DrrA |

**Table S20.** Summary of (a) differentially abundant vK-coordinates, and (b) corresponding functional information comparing **female** subjects with high (n = 10) versus low (n = 10) PC1 of problem behaviors

(a) vK-coordinates

| vK-coordinate | ANCOMBC Intercept | ANCOMBC Log fold change High PC1 cf. Low PC1 | OC_x | HC_y | Compound-associated vK mapping zone | Linked Function IDs |
|---|---|---|---|---|---|---|
| 0.593253968253968__1.59325396825397 | -3.21E-16 | -0.4083 | 0.5933 | 1.5933 | Amino sugars | fxn_12938;fxn_12942 |
| 0.922222222222222__1.37777777777778 | -1.20E-16 | 0.4173 | 0.9222 | 1.3778 | Tannins | fxn_14491 |

(b) linked functions

| Fxn ID | SEED Subsytem Level 1 | SEED Subsystem Level 2 | SEED Subsystem Level 3 | Function |
|---|---|---|---|---|
| fxn_12938 | Miscellaneous | - | EC 4.1.2.- Aldehyde-lyases | Dihydroneopterin_aldolase_(EC_4.1.2.25) |
| fxn_12942 | Miscellaneous | - | EC 4.1.2.- Aldehyde-lyases | Dihydroneopterin_phosphate_phosphatase_/_Dihydroneopterin_aldolase_(EC_4.1.2.25) |
| fxn_14491 | Nucleosides and Nucleotides | Pyrimidines | pyrimidine conversions | Uridine_phosphorylase_(EC_2.4.2.3) |

**Table S21.** Summary of (a) differentially abundant vK-coordinates, and (b) corresponding functional information comparing **male** subjects with high (n = 8) versus low (n = 9) PC1 of problem behaviors

(a) vK-coordinates

| vK-coordinate | ANCOMBC Intercept | ANCOMBC Log fold change High PC1 cf. Low PC1 | OC_x | HC_y | Compound-associated vK mapping zone | Linked Function IDs |
|---|---|---|---|---|---|---|
| 1.55__0.753333333333333 | 3.73E-02 | 0.8894 | 1.5500 | 0.7533 | demethylated | fxn_2926 |
| 0.583308133090742__1.44842868819318 | 2.00E-02 | 0.4742 | 0.5833 | 1.4484 | lignin | fxn_821;fxn_2958;fxn_2973; fxn_12703;fxn_12705;fxn_12786; fxn_12788 |

(b) linked functions

| Fxn ID | SEED Subsytem Level 1 | SEED Subsystem Level 2 | SEED Subsystem Level 3 | Function |
|---|---|---|---|---|
| fxn_12703 | Metabolism of Aromatic Compounds | Metabolism of central aromatic intermediates | Central meta-cleavage pathway of aromatic compound degradation | Acetaldehyde_dehydrogenase_(EC_1.2.1.10)_/_Alcohol_dehydrogenase_(EC_1.1.1.1) |
| fxn_12705 | Metabolism of Aromatic Compounds | Metabolism of central aromatic intermediates | Central meta-cleavage pathway of aromatic compound degradation | Alcohol_dehydrogenase_(EC_1.1.1.1)_@_Acetaldehyde_dehydrogenase_(EC_1.2.1.10)_@_Pyruvate-formate-lyase_deactivase |
| fxn_12786 | Metabolism of Aromatic Compounds | Peripheral pathways for catabolism of aromatic compounds | Biphenyl Degradation | Acetaldehyde_dehydrogenase_(EC_1.2.1.10)_/_Alcohol_dehydrogenase_(EC_1.1.1.1) |
| fxn_12788 | Metabolism of Aromatic Compounds | Peripheral pathways for catabolism of aromatic compounds | Biphenyl Degradation | Alcohol_dehydrogenase_(EC_1.1.1.1)_@_Acetaldehyde_dehydrogenase_(EC_1.2.1.10)_@_Pyruvate-formate-lyase_deactivase |
| fxn_2926 | Carbohydrates | Central carbohydrate metabolism | Pyruvate metabolism I: anaplerotic reactions, PEP | Phosphoenolpyruvate_carboxykinase_[GTP]_(EC_4.1.1.32) |
| fxn_2958 | Carbohydrates | Central carbohydrate metabolism | Pyruvate metabolism II: acetyl-CoA, acetogenesis from pyruvate | Acetaldehyde_dehydrogenase_(EC_1.2.1.10)_/_Alcohol_dehydrogenase_(EC_1.1.1.1) |
| fxn_2973 | Carbohydrates | Central carbohydrate metabolism | Pyruvate metabolism II: acetyl-CoA, acetogenesis from pyruvate | Alcohol_dehydrogenase_(EC_1.1.1.1)_@_Acetaldehyde_dehydrogenase_(EC_1.2.1.10)_@_Pyruvate-formate-lyase_deactivase |
| fxn_821 | Amino Acids and Derivatives | Aromatic amino acids and derivatives | Tryptophan catabolism | Acetaldehyde_dehydrogenase_(EC_1.2.1.10)_/_Alcohol_dehydrogenase_(EC_1.1.1.1) |
